# Structural basis of ligand-selective transcriptional activation in the MerR-family antibiotic resistance regulator AlbA

**DOI:** 10.64898/2026.09.21.753095

**Authors:** M. Di Palma, A. Grinzato, P. Andriollo, L. Kleebauer, M. Leusciatti, G. Morra, R.D. Süssmuth, J.M. Sutton, K.M. Rahman, R. A. Steiner

**Affiliations:** Department of Biomedical Sciences, University of Padova, via Ugo Bassi 58/B, Padova, 35131, Italy; Institute of Pharmaceutical Science, School of Cancer & Pharmaceutical Sciences, King’s College London, Franklin-Wilkins Building, 150 Stamford Street, London, SE1 9NH, UK; Department of Chemistry, Technische Universität Berlin, Berlin, 10623, Germany; Istituto di Scienze e Tecnologie Chimiche ‘G. Natta’ SCITEC, CNR, Milano, Italy; UK Health Security Agency, Research and Evaluation, Porton Down, Salisbury, SP4 0JG Wiltshire, UK; Randall Centre for Cell and Molecular Biophysics, King’s College London, New Hunt’s House, Guy’s Campus, London SE1 1UL, UK

## Abstract

Multidrug-resistant pathogens demand novel resistance-breaking strategies. The oligoarylamide albicidin and pyrrolobenzodiazepines (PBDs) are potent antibacterials with unrelated scaffolds, yet both are neutralized by *albA* gene products in many Gram-negatives: AlbA, an albicidin-responsive self-upregulating MerR-family factor, and the smaller AlbAS, its unique ligand binding domain (LBD) alone. How AlbA couples ligand sensing to transcriptional activation has remained elusive. Here, using an integrative multidisciplinary approach, we define this mechanism and show that albicidin and PBDs elicit different responses. Crystal structures reveal C8-linked PBDs bound to AlbAS at the N- and C-terminal subdomains (NTD, CTD) of the central tunnel in poses dictated by C8-tail chemistry. This plasticity sequesters diverse PBDs with nanomolar affinities. Cryo-EM shows AlbA as an autoinhibited dimer in which a reciprocal arm closes the NTD end of the partner’s tunnel, restricting PBD binding to the CTD while the DNA binding domains (DBDs) are mobile. Albicidin binding is incompatible with this arrangement, whereas PBDs, as CTD plugs, do not perturb the native equilibrium between autoinhibited and promoter-competent states. We visualized the latter by cryo-EM in the RNAP-DNA-AlbA complex. By blocking albicidin-mediated enhancement *in vitro*, PBDs act as resistance-breaking partners for albicidin and a route to overcoming *albA*-dependent resistance to oligoarylamide antibiotics.

## Introduction

Antimicrobial resistance (AMR) was associated with an estimated 4.95 million deaths in 2019, with numbers projected to rise sharply by 2050 if current trends persist^1–3^. Of greatest concern are the multidrug-resistant ESKAPE pathogens responsible for most hospital-acquired infections^4,5^, which feature prominently on the 2024 World Health Organization (WHO) bacterial priority pathogens list^6,7^. New druggable targets and chemical scaffolds are therefore crucial for next-generation antibiotic discovery, especially against Gram-negative pathogens^8–12^.

Pyrrolobenzodiazepines (PBDs) are sequence-selective DNA minor groove-binding agents with promising antitumor and antibacterial activities^13–17^. Their pyrrolo[2,1-c][1,4]benzodiazepine core (**Fig. 1a**) carries an *S*-chiral center at C11a that provides optimal complementarity for the DNA minor groove. Synthetic PBDs modified at C8 are promising potent antimicrobials against both Gram-positive and Gram-negative bacteria^15–17^. Their antibacterial mechanism is thought to combine DNA binding with inhibition of DNA gyrase^17,18^. In *Klebsiella pneumoniae*, PBD resistance to PBDs has been linked to sequence changes in the Tsx transporter and in the albicidin-binding protein AlbA, a MerR-family transcriptional regulator^19^. AlbA was originally discovered as a key driver of resistance to albicidin **(Fig. 1b)**, a potent natural antibiotic from *Xanthomonas albilineans*^20–25^. Albicidin binding triggers robust upregulation of *albA*, conferring resistance through a positive-feedback sequestration mechanism^22^. AlbA is therefore a critical determinant of convergent resistance to the structurally unrelated PBD and albicidin classes. As AlbA may also confer resistance to cystobactamids, promising natural antibiotics structurally related to albicidins^26–30^, its role in transcription regulation needs to be understood.

**Figure 1.**
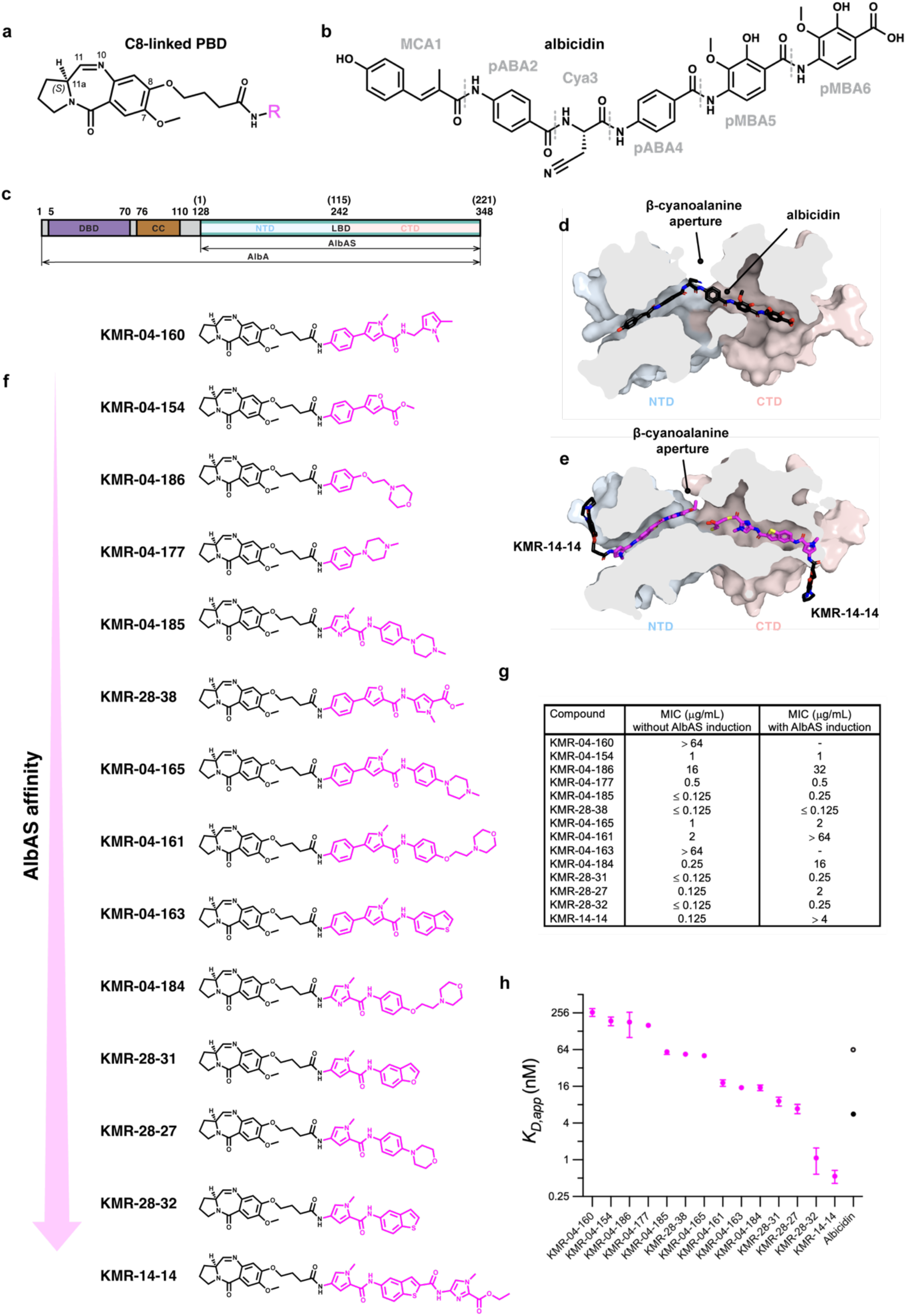
C8-linked pyrrolobenzodiazepine (PBD) conjugates bind AlbAS with nanomolar affinity and their antibacterial potency in *E. coli* is affected by AlbAS overexpression. (**a**) General structure of C8-linked PBD conjugates. The core is shown with standard ring numbering, and C8 is linked through an ether oxygen to a propyl-amide spacer connecting to the variable tail region R (magenta), which differs across the series. (**b**) Chemical structure of albicidin, a potent broad-spectrum peptide antibiotic from the plant pathogen *Xanthomonas albilineans*. Its six building blocks are: MCA1, methyl p-coumaric acid, pABA2 and pABA4, para-aminobenzoic acid, Cya3, β-cyano-l-alanine, and pMBA5 and pMBA6, 4-amino-2-hydroxy-3-methoxybenzoic acid. (**c**) Domain architecture of AlbA and AlbAS, with residue boundaries, those in brackets for AlbAS. AlbA comprises the N-terminal DNA-binding domain (DBD), the dimerization coiled-coil (CC) and the ligand-binding domain (LBD), itself a tandem repeat of two structurally similar N- and C-terminal domains (NTD and CTD). AlbAS is an alternative translation product comprising the LBD alone. (**d**) Surface of AlbAS bound to albicidin (black sticks, PDB 6ET8), in cross-section to reveal the internal binding channel, with the NTD and CTD lobes colored blue and pink, respectively. The cyanoalanine aperture, a solvent-exposed opening accommodating the central Cya3 moiety, is indicated. (**e**) Sliced surface of AlbAS bound to two KMR-14-14 molecules (black and magenta sticks, PDB 8RKY), showing the “tail-to-tail” binding mode in which both PBD heads protrude from opposite channel exits while the ester termini face each other within the protein. (**f**) Structures of all C8-linked PBD conjugates evaluated here, ordered from lowest (top) to highest (bottom) AlbAS affinity, with the variable tails in magenta. (**g**) Minimum inhibitory concentrations (MICs, µg/mL) against *E. coli* BL21(DE3) under non-inducing and AlbAS-inducing conditions. Dashes mark compounds not tested under induction because their control MIC exceeded 64 µg/mL. (**h**) Equilibrium dissociation constants of each conjugate for AlbAS, measured by tryptophan fluorescence quenching, on a log-2 *y*-axis. Published albicidin values are included for reference. Error bars are standard errors of the fitted *K*D,app (**Suppl. Table 1**).

Like other MerR family members, AlbA comprises a conserved N-terminal DNA binding domain (DBD) a coiled-coil (CC) dimerization region and a C-terminal ligand binding domain (LBD) specific to the effector recognized (**Fig. 1c**)^31–36^. Besides full-length AlbA (also known as AlbAL, L for long, 40 kDa, residues 1-348 in *K. oxytoca*), the *albA* gene produces a second in-frame translation product, AlbAS (25.8 kDa, S for short, residues 1-221 corresponding to residues 128-348 of *K. oxytoca* AlbA) consisting solely of the LBD^24^. The latter is a unique all-helical architecture of two structurally similar N-terminal and C-terminal domains sharing only 15.6% sequence identity (NTD, residues 1-115, and CTD, residues 116–221). In the AlbAS:albicidin complex the ligand straddles both domains within the ∼40 Å-long central tunnel (**Fig. 1d**), placing its MCA-1 and pMBA6 ends within the NTD and CTD, respectively^20,21^. In contrast, we have shown that two PBD molecules (KMR-14-14) bind AlbAS ‘tail-to-tail’, each independently stabilized by the NTD and CTD (**Fig. 1e**)^19^.

Here, using an integrative multidisciplinary approach that combines microbiology, affinity measurements, X-ray crystallography, AI-based structural predictions, single-particle cryo-EM, X-ray solution scattering and *in vitro* transcription assays, we define how AlbA senses its two antibiotic classes and translates this into a differential transcriptional response. While C8-linked PBD molecules bind with variable poses at the NTD and CTD ends of the AlbAS central tunnel depending on their tail chemistry, in the AlbA dimer the LBDs are autoinhibited, each protomer having its NTD end occluded by a ‘switch’ arm of its partner, which restricts PBD binding to the CTD. The DBDs and the CC remain conformationally heterogeneous. The rigid albicidin antibiotic cannot decouple NTD and CTD binding, shifting the equilibrium toward the ‘switch’-open state in which the LBDs separate and the DBDs become available for promoter binding. The F143A ‘switch’-open variant reproduced this shift without ligand, and we observed the promoter-competent state by cryo-EM at 2.4 Å in the RNAP:DNA:AlbA complex. Thus, AlbA discriminates its two ligand classes by tunnel reach rather than affinity, with PBDs as ‘CTD plugs’ unable to enhance transcriptional activation like albicidin. Our work shows that AlbA employs an ‘equilibrium-shift’ mechanism unconventional within the MerR-family. It also indicates that PBDs are resistance-breaking partners for albicidin and a route to overcoming AlbA-dependent resistance to oligoarylamide antibiotics.

## Results

### The C8-linked tail of PBDs modulates AlbAS recognition and AlbAS-mediated resistance in *E. coli*

To assess how C8-linked tail substituents affect AlbAS-induced resistance in *E. coli*, we tested a panel of PBD derivatives differing in aromatic and aliphatic ring length and composition (**Fig. 1f**). As AlbA orthologues are conserved across *Klebsiella* and the Gram-negative ESKAPE pathogens (**Suppl. Fig 1**), the resistance conferred by *K. oxytoca* AlbAS in a heterologous *E. coli* host is taken as a proxy for AlbA function in these bacteria. Minimum inhibitory concentrations (MICs) were determined in BL21(DE3) carrying an inducible *K. oxytoca* AlbAS plasmid (**Fig. 1g**, **Suppl. Fig. 2**). Without AlbAS induction (control conditions) most compounds were strongly growth-inhibitory (MIC < 2 µg mL⁻¹). Only KMR-04-160 and KMR-04-163 displayed MICs > 64 µg mL⁻¹, likely because of poor membrane permeability^17^. Upon AlbAS induction, three compounds exhibited a ≥32-fold increase in their MIC (KMR-04-161, KMR-04-184, KMR-14-14), and a fourth (KMR-28-27) a 16-fold increase, indicating AlbAS-mediated resistance. As several MICs lie at the limits of the tested range, the corresponding fold-shifts are bounds rather than point estimates (**Suppl. Fig. 2**).

MIC shifts are only an indirect measure of AlbAS binding. Thus, we quantified affinities by intrinsic tryptophan fluorescence quenching (**Fig. 1h**). Although crystallographic data for KMR-14-14 (**Fig. 1e**) and other PBD conjugates (*vide infra*) reveal two ligand molecules bound per AlbAS molecule, our titrations were well described by a single binding transition within the range explored, bounded above ∼1 µM by strong intrinsic ligand fluorescence and inner-filter effects (**Suppl. Fig. 3, 4**). The reported affinity values (*K*_D,app_, **Fig. 1h**) are therefore apparent macroscopic for overall binding to AlbAS rather than for individual site (**Suppl. Table 1**). They the series and show that half of the fourteen conjugates bound AlbAS with a *K*_D,app_ at or below 50 nM, four of them at or below 10 nM. Albicidin affinity for AlbAS has been reported alternatively as 64 nM or more recently as 5.6 ± 0.2 nM (open and full black circles, respectively, in **Fig. 1h**)^20,37^.

Across the panel, AlbAS-mediated resistance required two partly separable features. Affinity was set mainly by the ring directly linked to the anilide amide, the four *N*-methylpyrrole conjugates being the tightest binders (*K*_D,app_ ≤ 10 nM). Tight binding alone, however, was not sufficient for neutralization. Of those, the two two-ring tails, KMR-28-32 and KMR-28-31 at second and fourth, shifted their MIC only ∼2-fold, an order of magnitude less than the two longer-tailed members of the class. The most efficiently neutralized compounds instead combined measurable affinity with a longer tail, in three of four cases terminating in a morpholine: KMR-04-184 (64-fold), KMR-04-161 (≥32-fold), KMR-28-27 (16-fold) and KMR-14-14 (≥32-fold). The morpholine correlation is not absolute, as KMR-04-186 carries one but shifted only 2-fold, consistent with its much weaker affinity. Conversely, KMR-04-154 and KMR-04-177 (*K*_D,app_ > 150 nM) showed no AlbAS-dependent MIC change. KMR-04-160 and KMR-04-163 offered no window (MIC > 64 µg mL⁻¹ uninduced), so protection could not be assessed. KMR-04-163 is the sixth tightest binder of the series and is captured in complex with AlbAS crystallographically (*vide infra*), indicating that its inactivity has a cellular rather than a recognition basis.

### The PBD core preferentially occupies a conserved inner pocket while the C8 tail is accommodated flexibly

To generalize the molecular basis of PBD recognition we determined X-ray crystal structures of AlbAS bound to KMR-04-154, KMR-04-161, KMR-04-163, KMR-04-165, KMR-04-177 and KMR-28-27. Crystals belonged to the monoclinic space group *C*2 or, for KMR-04-161 and KMR-04-165, to the tetragonal *P*4_3_2_1_2 space group, and diffracted between 1.6 and 3.0 Å. Data collection and refinement statistics are in **Suppl. Table 2**.

We previously observed two KMR-14-14 molecules arranged “tail-to-tail” with their ester termini facing each other within the AlbAS core while their PBD “heads” protrude out of the channel (**Fig. 1e**)^19^. None of the compounds here adopts this organization. The predominant binding mode (KMR-04-154, KMR-04-161, KMR-04-163, KMR-04-177) has one PBD “head” anchored in the inner CTD channel with its tail projecting toward the exit, while a second PBD head protrudes through the central β-cyanoalanine aperture (**Fig. 2a**).

**Figure 2.**
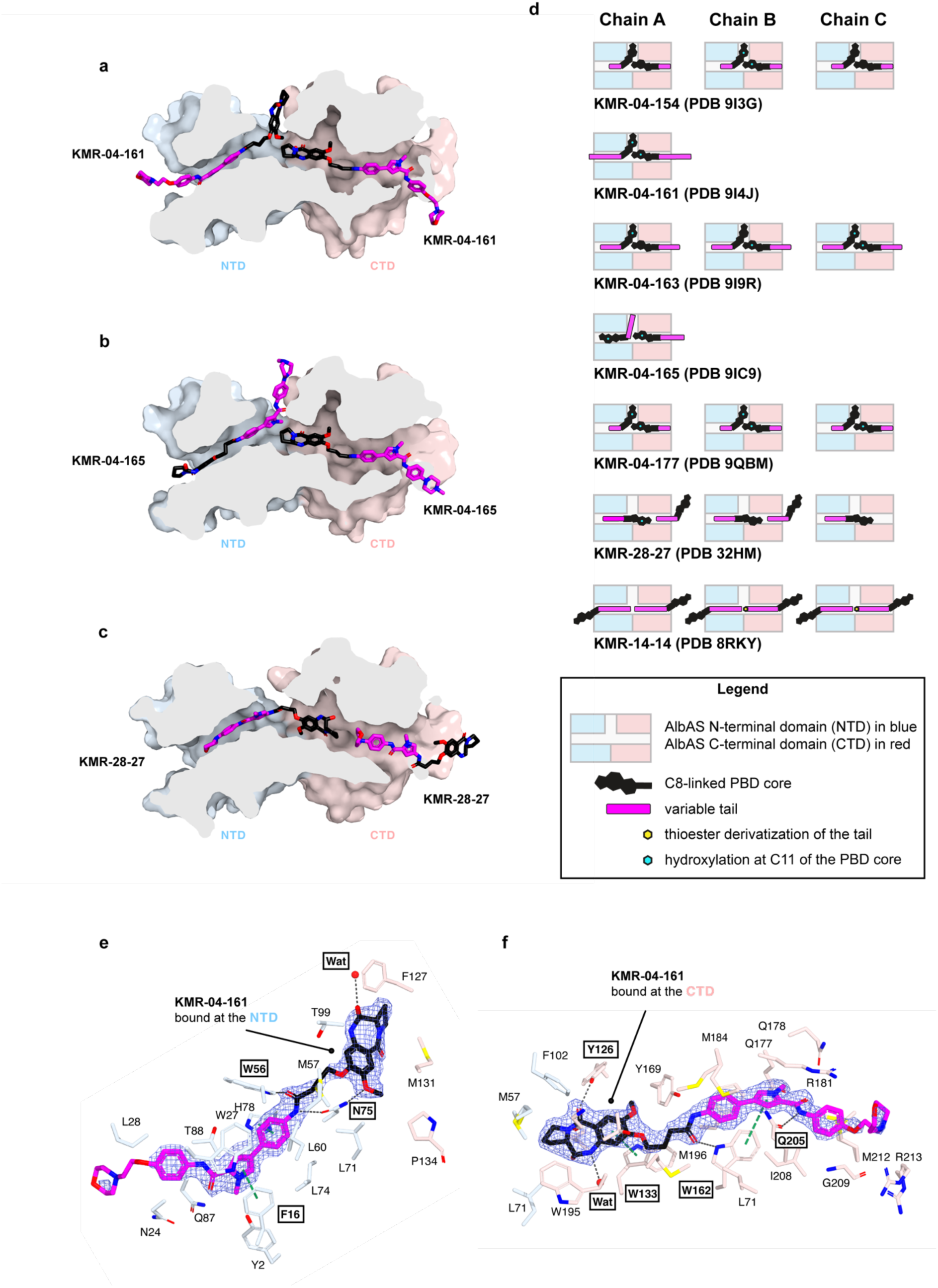
Crystal structures of AlbAS in complex with C8-linked PBD conjugates reveal both conserved and variable recognition features. (**a**–**c**) Surface representations of AlbAS, shown as cross-sections through the central binding channel to expose the bound ligands, with the N-terminal domain (NTD) colored light blue and the C-terminal domain (CTD) light pink. The two independently bound PBD molecules are drawn as sticks in black and magenta and are labeled individually; heteroatoms are colored by element (nitrogen, blue; oxygen, red). (**a**) AlbAS–KMR-04-161 (PDB 9I4J), representative of the predominant binding mode, in which one PBD head is anchored in the inner CTD channel with its tail projecting toward the channel exit while the second molecule presents its head at the central β-cyanoalanine aperture. (**b**) AlbAS–KMR-04-165 (PDB 9IC9), in which CTD binding is preserved but the NTD-bound ligand is inverted, directing its tail rather than its head into the aperture. (**c**) AlbAS–KMR-28-27 (PDB 32HM), in which the CTD-bound PBD head occupies approximately the canonical position, but its tail is redirected toward the NTD channel exit. (**d**) Schematic summary of the observed binding modes, drawn separately for each protein chain in the asymmetric unit. (**e**, **f**) Close-up views of various conserved protein–ligand contacts taking the AlbAS-KMR-04-161 structure as an example. Electron density maps (2*mF*o – *DF*c) for the ligands is represented as a blue mesh at the 1σ level. (E) and (F) are shown in different orientations to improve clarity. Protein side chains are shown as sticks and colored by domain. Hydrogen bonds are drawn as black dashed lines and π–π stacking interactions as green dashed lines. Residues involved in these interactions are highlighted. Ordered water molecules are shown as red spheres (Wat). In both ligands the PBD imine group is present in its hydrated carbinolamine form.

For KMR-04-165, CTD binding is conserved, but the NTD pose is inverted, directing the tail into the aperture (**Fig. 2b**). KMR-28-27 differs again. One PBD head occupies roughly the same CTD location but its tail is redirected toward the NTD channel exit (**Fig. 2c**). Electron density indicates lower occupancy and is visible mostly for the tail. A summary of these binding modes is provided in **Fig. 2d**.

Despite this positional diversity, several core host–ligand interactions are broadly conserved. In KMR-04-161, taken as representative, the NTD head group sits within the solvent-exposed β-cyanoalanine aperture and is primarily stabilized by N75 (AlbAS numbering) via dual hydrogen bonds to the spacer amide and the C8 ether oxygen. W56 can similarly hydrogen-bond to the spacer amide, while F16 mediates auxiliary π–π stacking with the tail’s aromatic scaffold (**Fig. 2e**). At the CTD, the PBD core buries deeply to stack with W133 and hydrogen bond through its lactam carbonyl to Y126, while the C8-linked tail stacks its aromatic rings against W162, accepts a hydrogen bond from the W162 indole nitrogen at its spacer amide, and forms additional hydrogen bonds with Q205 (**Fig. 2f**). Interactions for all other ligands are in **Suppl. Fig. 5.** For several bound ligands the PBD imine is present as the hydrated carbinolamine, consistent with its known reversible hydration in aqueous media (**Suppl. Fig. 6**)^13,14^.

These structures reveal a plastic recognition mode in which the PBD core almost invariably anchors within the CTD channel through the same contacts, while the C8 tail determines the overall binding mode and channel occupancy, providing a framework for hypothesis-driven tail modification.

### A reciprocal N-terminal arm occludes the NTD end of the central tunnel in the AlbA dimer

As no experimental structure of full-length AlbA exists, we expressed and purified *K. oxytoca* AlbA for downstream structural analysis. SEC-MALS confirmed it as a dimer in solution (**Suppl. Fig. 7**). AlbA crystals could be obtained but diffracted only to 10 Å (**Suppl. Fig. 8**). We therefore turned to single-particle cryo-electron microscopy (cryo-EM) and reconstructed dimeric ∼80 kDa AlbA at 3.6 Å resolution. Data collection, processing statistics and workflow are summarized in **Suppl. Table 3** and **Suppl. Fig. 9**.

In the reconstructed AlbA map the two LBDs associate through their NTDs about a two-fold axis. Each protomer projects its N-terminal segment across the dyad and into the partner subunit, capping the NTD aperture of the partner tunnel and leaving only the CTD ends accessible (**Fig. 3a**, **b, c**). The DBDs, the dimerization CC, and the first eleven residues of each LBD could not be resolved, and 2D class averages showed density for these regions either as a diffuse smear or not at all (**Suppl. Fig. 10**), indicating pronounced positional heterogeneity relative to the LBDs. Such mobility may also explain the poor X-ray diffraction of AlbA crystals.

**Figure 3.**
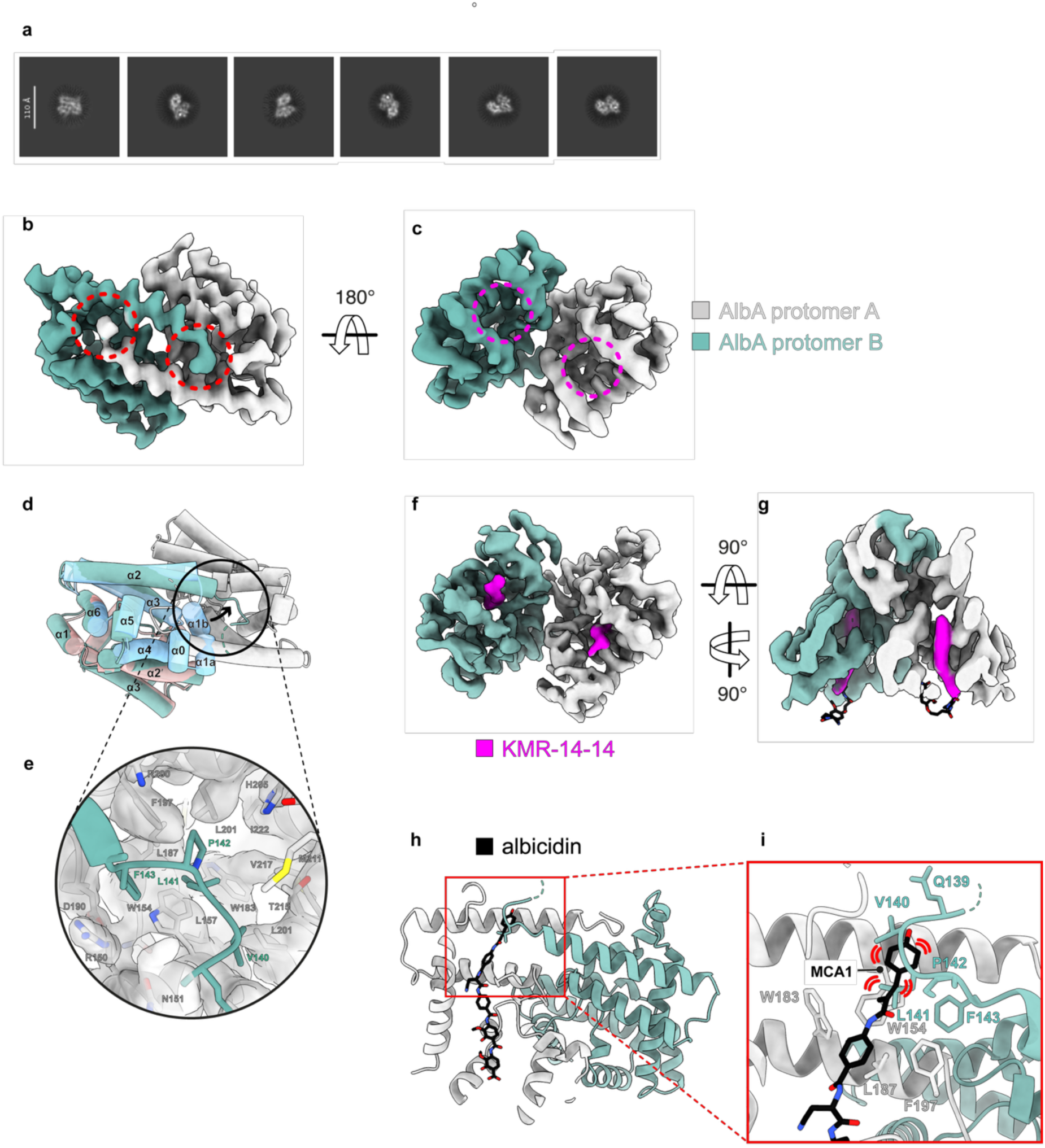
Cryo-EM structure of the AlbA dimer and occlusion of the NTD ligand site. (**a**) Representative 2D class averages of AlbA in multiple orientations, showing well-defined density for the dimeric LBD core. Density for the N-terminal DBDs and the CCs is not resolved, consistent with conformational flexibility in this region (see also **Suppl.** Fig. 10). Scale bar, 110 Å. **(b, c)** Cryo-EM map of AlbA viewed along two orientations related by a 180° rotation, with protomer A in gray and protomer B in teal. Dashed red circles in (a) mark the closed off NTD apertures while dashed magenta circles in (b) highlight the available CTD access to the channel. **(d)** Superposition of monomeric AlbAS (light blue and light red, NTD and CTD) on one protomer of the AlbA dimer (teal), with the partner protomer in gray. Helices are labeled following the AlbAS nomenclature. The arrow indicates the displacement of the N-terminal segment, which in the dimer leaves the position occupied by α1b in the monomer and crosses the dyad into the partner subunit. The circle marks the region enlarged in (d). **(e)** Close-up of the N-terminal arm of protomer B (teal) inserted into the partner protomer (gray). Side chains lining the pocket are shown as sticks. L141 packs against W154, L187, F197, L201 and V217 of the partner, and F143 makes a cation-π with R150. **(f, g)** Cryo-EM map of the AlbA:KMR-14-14 complex in two orthogonal views, colored as in (a, b), with density for KMR-14-14 in magenta. Ligand density is present only at the CTD site of each protomer. **(h)** Superposition of the AlbAS:albicidin complex (PDB 6ET8) on one protomer of the AlbA dimer, with albicidin in black. The box marks the region enlarged in (h). **(i)** Close-up of the albicidin MCA1 unit at the dimer interface. Red arcs mark steric overlap between the MCA1 unit and residues of the partner N-terminal arm.

The LBD interface in the AlbA dimer buries a surface area of approximately 2,550 Å² (1,275 Å² per protomer) contributed mainly by the first half of α2 and its N-terminal extension, together with the C-terminal portion of α3 and the α3–α4 turn (**Suppl. Fig. 11**).

However, the complex formation significance score (CSS) computed with PDBePISA^38^ is only 0.1, indicating that the observed LBD interface alone is unlikely to sustain dimerization. This is therefore maintained predominantly by the CC within the N-terminal domain, which in isolated AlbA is conformationally independent of the LBDs.

Relative to monomeric AlbAS, where α1b and α2 are roughly orthogonal, α2 is extended N-terminally in the dimer. The preceding segment lacks regular secondary structure and carries the chain across the dyad into the partner subunit, occluding the NTD entry to the central tunnel (**Fig. 3d**). Specifically, L141 of each protomer inserts into a conserved hydrophobic pocket formed by W154, L187, F197, L201 and V217 of the partner subunit (**Fig. 3e**), reciprocally blocking the NTD aperture. The conserved F143 completes this arrangement through a cation–π interaction with R150 of the second monomer.

This arrangement suggested that only one PBD conjugate would bind per LBD, at the CTD site, since the NTD site is inaccessible. To test this, we obtained reconstructions for the AlbA:KMR-14-14 and AlbA:KMR-28-27 complexes at 3.5 Å and 4.3 Å, respectively (**Suppl. Table 3** and **Suppl. Fig. 12**, **13**). As predicted, ligand density was seen only at the CTD site, extending ∼20 Å into the tunnel, while the LBD dimer remained unaltered (**Fig. 3f, g**). Tryptophan quenching against AlbA for a subset of four ligands showed all bound more weakly than to AlbAS, by 1.7-, 3.0-, 9.1- and 42-fold for KMR-04-177, KMR-04-161, KMR-14-14 and KMR-28-27, respectively (**Suppl. Fig. 14**, **Suppl. Table 4**). The large loss of affinity for KMR-28-27 is well rationalized by the structural data. Unlike the other conjugates, whose most conserved AlbAS pose already directs the tail toward the CTD exit, KMR-28-27 binds AlbAS with its tail projecting into the NTD channel (**Fig. 2c**), a pose not possible in the dimer forcing it into a disfavored CTD-only pose (**Suppl. Fig. 13 inset**). The shift therefore reflects the cost of changing binding mode rather than the loss of a second site common to all ligands.

Critically, superposition of the available AlbAS:albicidin structures (PDB 6ET8 and 6H96) on a dimer-engaged LBD protomer shows the antibiotic’s MCA1 moiety clashing with the stretch around V140 and L141 (**Fig. 3h, i**), indicating that binding of the rigid albicidin molecule is incompatible with the observed arrangement. Thus, we posit that the interaction with albicidin displaces the region centered around L141 which we define as the ligand-sensing ‘switch’, thereby triggering a rearrangement which can be transmitted to the N-terminal DBDs. Supporting this, the MCA1 unit (**Fig. 1b**) is key for transcription activation and that transcription activity is mainly modulated by modifications at the MCA1 end of albicidin^22,39^.

### The ‘switch’ region controls AlbA conformational transitions

AI-driven structure prediction (ColabFold-AlphaFold2^40,41^, AlphaFold3^42^) provided complete AlbA models of high confidence for most of the individual domains but not for their relative orientations. Although predictions failed to capture the LBD dimer observed experimentally, they concurred on CC-mediated dimerization and proposed models ranging from having proximal LBDs (LBD-close) to LBDs splayed apart (LBD-open) and from having DBDs tucked in between the CCs and the LBDs (DBD-in) to being solvent exposed on the opposite side of the LBDs (DBD-out) (**Fig. 4a, b, c, d**). Chai-1^43^ models restrained to the experimental cryo-EM dimer also returned a mixture of DBD-in and DBD-out structures (**Fig. 4e, f, g, h**). Their consistently low pLDDT scores identify as nodal points both the junction between the DBD and the CC and that between the CC and the LBD, the latter encompassing the ‘switch’.

**Figure 4.**
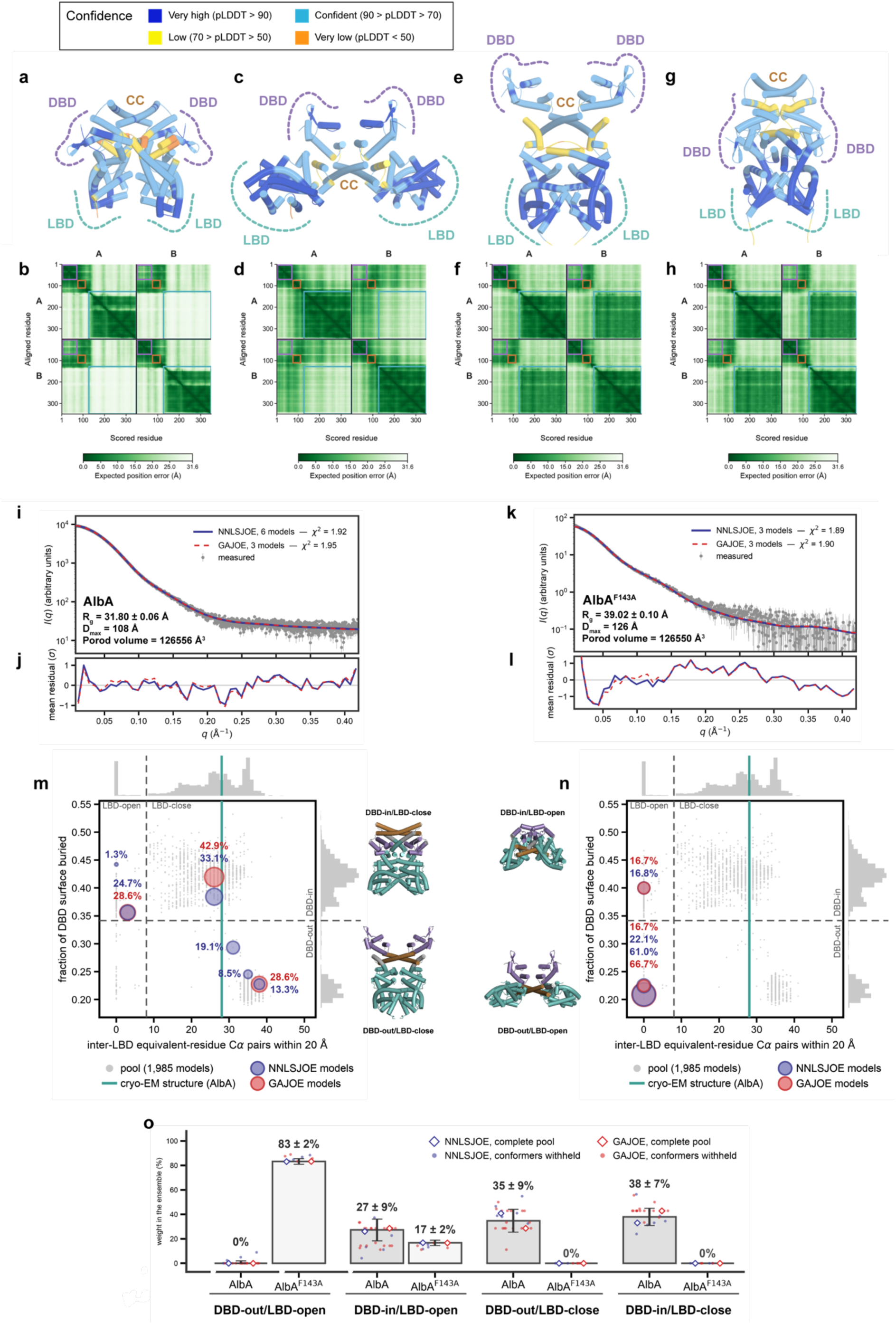
Conformational heterogeneity of the AlbA dimer and its redistribution by the AlbA^F143A^ ‘switch’ variant. (**a**–**h**) Predicted structures of the AlbA dimer (chains A and B). Each cartoon (**a**, **c**, **e**, **g**) is paired with its predicted aligned error matrix (**b**, **d**, **f**, **h**). Color bars give pLDDT and expected position error in Å. Dashed outlines mark the DBD, CC and LBD regions. (a, b) from AlphaFold3, (c, d) from ColabFold-AlphaFold2, (e–h) from Chai-1 restrained to the cryo-EM LBD dimer. Domains are predicted confidently but their relative orientations are not, the largest expected errors falling in the chain-to-chain blocks and at the DBD–CC and CC–LBD junctions, including the ‘switch’. (**i**) SAXS curve of AlbA. Measured intensities with uncertainties in gray, ensemble fits blue (NNLSJOE) and red dashed (GAJOE). Conformer count and χ^2^ in the key. *R*g, *D*max and Porod volume are on the panel. (**j**) Residuals for the fits in (i), averaged in *q* bands and scaled by the measurement uncertainty. (**k**, **l**) Like (i, j) for AlbA^F143A^. (**m**, **n**) Conformational maps of the ensembles for AlbA (k) and AlbA^F143A^ (n), on identical axes and from the same 1,985-model pool. Axes are inter-LBD equivalent-residue Cα pairs within 20 Å against the fraction of DBD surface buried, dashed lines dividing the four classes, which are named along the top and right edges. Gray, the pool, with marginal histograms. The vertical line marks the wild-type cryo-EM structure on both panels. Discs mark selected conformers, blue NNLSJOE and red GAJOE, area proportional to fitted weight, shown as percentage. Cartoons show a representative of each occupied class (DBD violet, CC brown, LBD teal, connecting regions gray). (**o**) Ensemble composition by conformational class. Bars are the mean of the two algorithms’ fits. Error bars are the standard deviation over all fits of that protein (26 for AlbA, 10 for AlbA^F143A^), comprising Monte-Carlo replicates of the intensities, independent GAJOE seeds, both algorithms and the jackknife withholding selected conformers in turn. Open diamonds, the complete fits. Small points, individual withholding fits. Values above the bars, mean and standard deviation as percentages.

To probe conformational heterogeneity further we measured SEC-SAXS data on AlbA. Analyses with the ATSAS package^44^ confirmed it as a dimer (MW = 82,279 ± 6,210 Da, theoretical = 80,165.5 Da) with *R*_g_ = 31.80 ± 0.06 Å and *D*_max_ = 108.1 Å (**Suppl. Table 5**). Using Boltz-2^45^ we then generated a pool of 1,985 unique structures sampling DBD/LBD orientations within the dimer. No single conformer accounted for the measured scattering.

The best model gave *χ*^2^ = 2.63, was less extended than the particle in solution (*D*_max_ = 102 Å), and its residuals oscillated systematically (**Suppl. Fig. 15**). Poor fit was concentrated at real-space distances of 30–57 Å, the scale of the separation between domains within the dimer, placing the discrepancy in their relative arrangement and leading us to consider mixtures of conformers. Non-negative least-squares selection with NNLSJOE^46,47^ returned six conformers at *χ*^2^ = 1.92, and the genetic algorithm GAJOE, run independently, reached *χ*^2^ = 1.95 with three models (**Fig. 4i, j**) reproducing the experimental dimensions. Multiple independent runs with both methods showed that the solution is robust and dominated by LBD-close conformers, whose inter-LBD arrangement is consistent with that of the cryo-EM structure, and in which the DBDs are predominantly tucked in (DBD-in) (**Fig. 4k**).

As F143 in the ‘switch’ region ranks amongst the most buried interface residues in the cryo-EM structure (217 Å^2^, 8.5 % of the interface area), we reasoned that an alanine replacement might destabilize the LBD-close arrangement. A phenylalanine at this position also totally conserved amongst orthologues (**Suppl. Fig. 1**). SAXS data on AlbA^F143A^ confirmed a more elongated shape than the wild type (*R*_g_ = 39.02 ± 0.10 Å, *D*_max_ = 126 Å), with the classifier changing from compact to flat (**Fig. 4l, m**). The Porod volume remained essentially identical, indicating a change in shape rather than in oligomeric state. Ensemble fitting on the same pool resolved that elongation into a specific rearrangement. Both algorithms place every selected conformer at zero inter-LBD contacts, that is, entirely within the LBD-open classes, and the majority with the DBDs solvent exposed (DBD-out) (**Fig. 4n**).

Quantifying both ensembles over all independent fits (different seeds and jackknife resampling), LBD-open rises from 27 ± 9 % in the wild type to 100 % in the variant and DBD-out from 35 ± 10 % to 83 ± 2 %, summing the classes in (**Fig. 4o**) that share each domain state. The F143A replacement in the ‘switch’ region therefore shifts the inter-LBD arrangement from predominantly closed, as seen in the cryo-EM structure, to entirely separated, and this translates into an increased exposure of the DBDs.

### AlbA recognizes a conserved promoter dyad whose affinity increases upon LBD opening

To refine our knowledge of the DNA sequence preference of AlbA we performed unbiased systematic evolution of ligands by exponential enrichment (SELEX) based on the electrophoretic mobility shift assay (EMSA) (**Suppl. Fig. 16**). Sequence analysis after seven rounds of enrichment revealed the 9-bp ACNCTNACG motif (33-fold enrichment over shuffled controls) as the primary recognition sequence (**Fig. 5a**). This motif corresponds to the sequence spanning the 3’ end of the −35 element and the adjacent 19-bp spacer of the *K. oxytoca albA* promoter (*palbA*) region (**Fig. 5b**). An essentially invariant motif lies upstream of putative *albA* genes in *K. pneumoniae*, *P. aeruginosa* and *A. baumannii*, indicating conservation across Gram-negative species (**Fig. 5b, c**).

**Figure 5.**
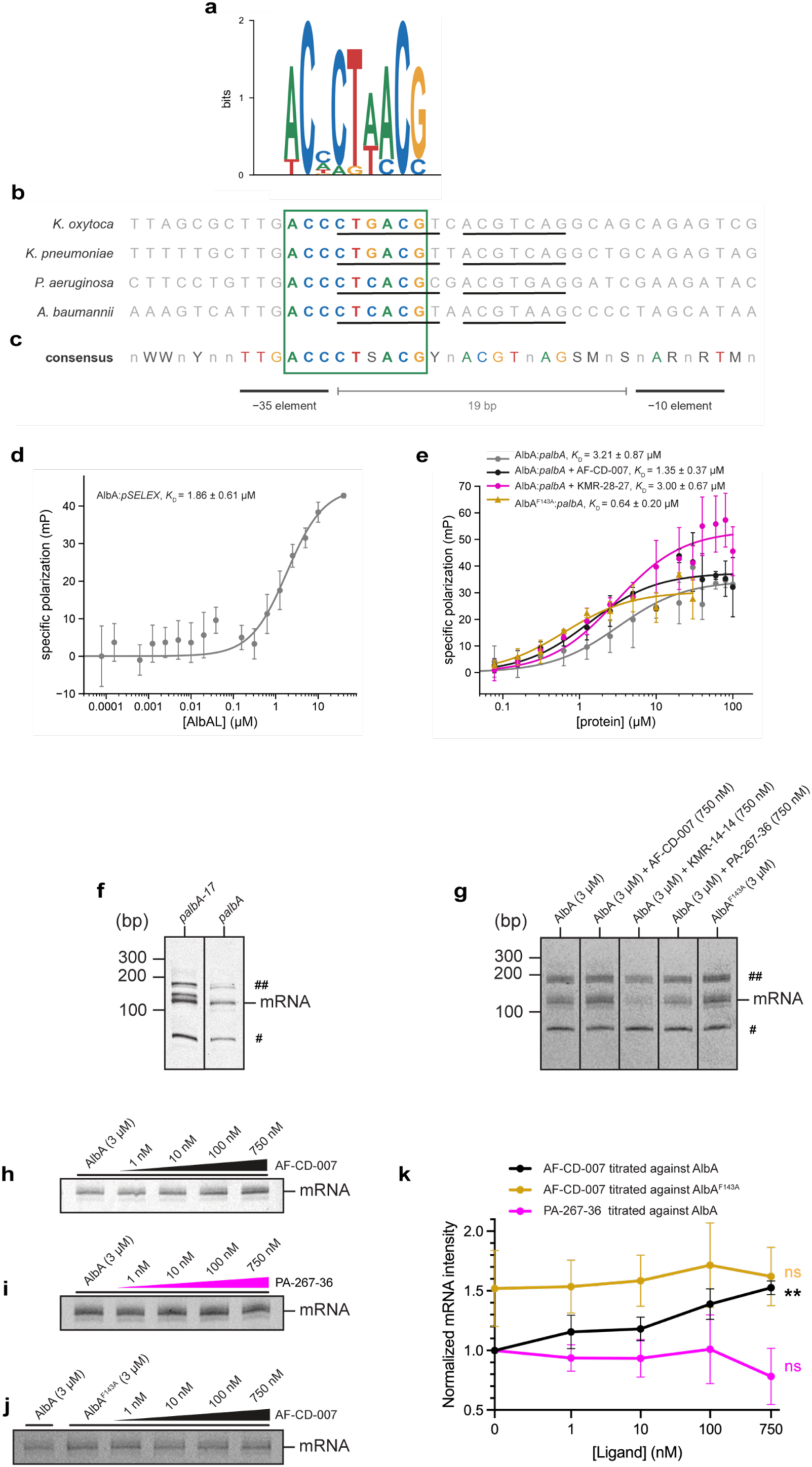
‘Switch’ opening enhances AlbA-driven transcription from its own promoter *in vitro* and is triggered by albicidin but not by PBDs. (**a**) Sequence logo of the 9-bp motif after seven rounds of SELEX. (**b**) The motif (green box) in the *albA* promoter region from *K. oxytoca* and three ESKAPE group pathogens, with motif positions colored by base and flanking sequence gray. Perfect or near-perfect palindromic sequences are underlined. (**c**) Consensus in which the motif spans the 3′ end of the −35 element and the adjacent spacer, the −35 and −10 elements being separated by 19 bp instead of the optimal 17 bp. (**d**) Binding of AlbA to a duplex carrying the SELEX-derived motif (*pSELEX*) measured by FP. (**e**) Binding by FP of AlbA to its own promoter (*palbA*) alone (gray), with the albicidin derivative AF-CD-007 (black) or with KMR-28-27 (magenta), and of the AlbA^F143A^ variant to *palbA* without ligand (gold). In (d) and (e) specific polarization is plotted against total protein on a log scale. Points are mean ± standard deviation of three independent experiments and solid lines are fits to a single-site model with a nonspecific term. Fitted *K*D values are reported with their standard error. (**f**) Molecular beacon transcription assay with RNAP alone on a variant in which the spacer is shortened to 17 bp (*palbA-17*) and on the native *palbA* promoter, showing that the native spacer is suboptimal for RNAP. # = abortive transcript, ## = runoff transcript. (**g**) Transcription from *palbA* with AlbA, AlbA plus AF-CD-007, KMR-14-14, AlbA or the KMR-14-14 dilactam PA-267-36, and with AlbA^F143A^. (**h–j**) Titrations of AF-CD-007 against AlbA (h), PA-267-36 against AlbA (i) and AF-CD-007 against AlbA^F143A^ (j). (**k**) Quantification of (h–j), with band intensities normalized to AlbA without ligand. Points are mean ± standard deviation of three experiments. AF-CD-007 against AlbA increases transcription in a statistically significant concentration-dependent manner (one-sample *t* test on the fitted slope, two-tailed P = 0.0032). PA-267-36 against AlbA and AF-CD-007 against AlbA^F143A^ show non-significant deviations from the null slope (P = 0.4233 and 0.5514, respectively). Concentration values are on a symlog scale.

We next measured binding affinities by fluorescence polarization (FP). Using a 42-bp DNA fragment labeled at the 3’ of one strand and containing a 30-bp SELEX-derived sequence (*pSELEX*, **Suppl. Table 6**), we found that AlbA bound with a *K*_D_ of 1.9 ± 0.6 μM (**Fig. 5d**). We obtained *K*_D_ = 3.2 ± 0.9 μM (**Fig. 5e**, gray curve) for its own *palbA* promoter, marginally lower than the 7.8 ± 1.1 μM reported previously from EMSA measurements^22^. We then tested the effect of ligands: in the presence of the albicidin derivative AF-CD-007 (**Suppl. Fig. 17**) we obtained *K*_D_ = 1.7 ± 0.4 μM (**Fig. 5e**, black curve), a 1.9-fold improvement over the wild type in line with the 1.6-fold improvement reported for albicidin^22^ while addition of KMR-28-27 did not result in a change in affinity (*K*_D_ = 3.0 ± 0.7 μM, **Fig. 5e**, magenta curve). For the AlbA^F143A^ ‘switch’ variant, which populates the DBD-out/LBD-open state we obtained a further increase in affinity (*K*_D_ = 0.6 ± 0.2 μM, **Fig. 5e**, gold curve).

Overall, these findings support a mechanism in which destabilization of the LBD-close state is transmitted to the DBDs, resulting in an improved affinity for its own promoter. This is rationalized by a shift toward a higher population of DBD-out/LBD-open conformers competent for DNA binding.

### Unlike albicidin, a high-affinity PBD does not enhance AlbA-mediated transcription *in vitro*

We next reconstituted an *in vitro* AlbA-dependent transcription system using purified components and a 178 bp PCR fragment containing the native *palbA* promoter. This eliminated confounding cellular effects such as efflux or permeability differences, allowing ligand-dependent activation to be measured directly.

MerR transcription factors activate transcription by distorting promoters in which the −35 and −10 elements are separated by 19/20 bp instead of the optimal 17 ± 1 bp distance thus hindering recognition by the RNA polymerase (RNAP)^34,35,48–50^. A control experiment in the absence of AlbA confirmed that our assay recapitulated the expected behavior. Basal transcription was higher from a promoter in which the 19 bp spacer between the −35 and −10 elements had been shortened to 17 bp (*palbA-17*) compared to *palbA* (**Fig. 5f**).

We next tested the effect of ligands on transcription. While the albicidin derivative AF-CD-007 increased AlbA-mediated *palbA* transcription (lanes 1,2 **Fig. 5g**), capturing the known ligand-responsive behavior⁴⁷, addition of KMR-14-14 had an apparent inhibitory effect (lanes 1,3 **Fig. 5g**). As PBDs are minor groove binders and might affect transcription independently of AlbA, we also tested the KMR-14-14 dilactam (PA-267-36) (**Suppl. Fig. 18**) which lacks the imine responsible for covalent DNA modification. Unlike KMR-14-14, the dilactam did not alter control levels of transcription (lanes 1,4 **Fig. 5g**). Compared to AlbA, the AlbA^F143A^ variant showed an increase in mRNA levels (lanes 1,5 **Fig. 5g**).

Titration experiments quantified these effects. AF-CD-007 exhibited a clear concentration-dependent effect (two-tailed P = 0.0032) reaching approximately 1.5-fold at 750 nM (**Fig. 5h**, **k**), whereas no such trend was detected for PA-267-36 (P = 0.4233, **Fig. 5i**, **k**). When titration of AF-CD-007 was performed against AlbA^F143A^, concentration dependence was lost (P = 0.5514, **Fig. 5j**, **k**) with the variant affording a ∼60% increase in mRNA compared to wild type AlbA.

These results support a model in which nanomolar-affinity PBD binders occupy the CTD-side of the hydrophobic tunnel without displacing the ‘switch’, thereby preventing the albicidin-induced opening of the LBDs that promotes transcriptional activation.

### AlbA undergoes a major conformational change when bound to its promoter DNA and the RNA polymerase (RNAP)

To understand how AlbA facilitates promoter recognition by the RNA polymerase (RNAP) we reconstituted the ternary RNAP:*palbA*:AlbA complex and solved its structure by single particle cryo-EM (**Suppl. Table 3**). The overall 2.4 Å resolution drops at the periphery where AlbA is bound to its promoter (4-5 Å local resolution for DBDs and CC, >6 Å for LBDs).

Focused refinement on AlbA yielded a much better reconstruction (3 Å, **Suppl. Fig. 19**).

Promoter-bound AlbA exhibits a ‘butterfly-like’ DBD-out/LBD-open arrangement in which its LBDs diverge from the CC junction by ∼150°, lifting them away from the duplex, while the DBDs engage with it 42 Å apart over 20-bp, reaching into the −35 element but stopping one base pair before the −10 element (**Fig. 6a, b**, **c**). Density for KMR-28-27, which we added to the RNAP:*palbA*:AlbA complex post-reconstitution, is also seen at its primary binding location identified in the crystallographic AlbAS complex (**Suppl. Fig. 20**). Some additional density in the minor groove between the DBDs (**Suppl. Fig. 21**) might correspond to a low-occupancy DNA-bound PBD, given the slow kinetics of covalent modification.

**Figure 6.**
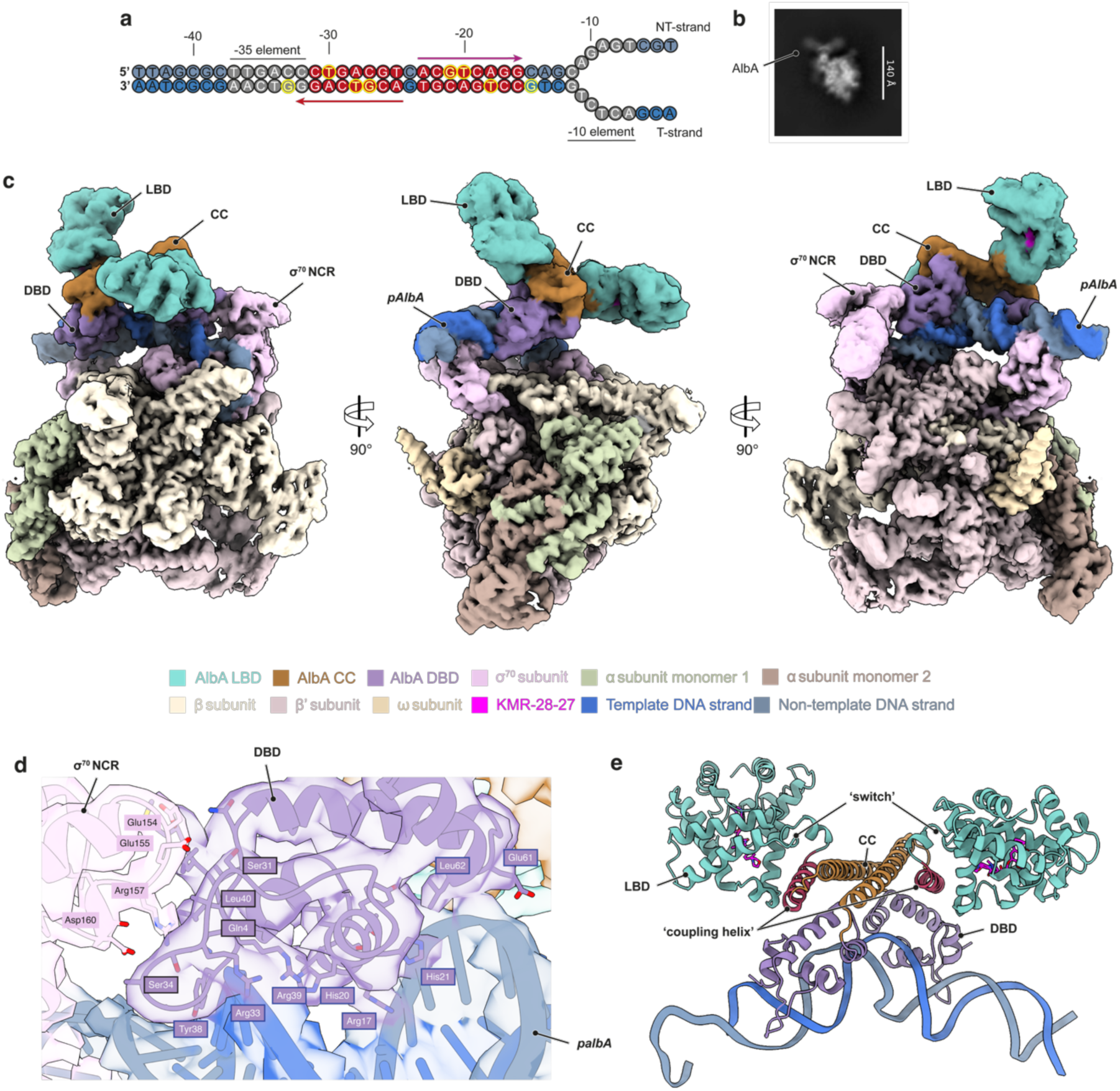
Cryo-EM structure of the RNAP:*palbA*:AlbA initiation complex in a promoter-engaged, ‘butterfly-like’ conformation. (**a**) Sequence of the *palbA* promoter duplex used for complex reconstitution, numbered relative to the transcription start site, with the −35 and −10 elements underlined. On red background the palindromic recognition motif, which spans the 3′ end of the −35 element and the adjacent spacer. Arrows mark the two dyad-related half-sites read by each DBD. Highlighted in yellow is the base read by AlbA in each of the eight base pairs critical for sequence discrimination, two of which, at −33 and −15, lie just outside the palindrome. NT-strand and T-strand denote the non-template and template strands, respectively. (**b**) Representative 2D class average of the complex, with the AlbA density protruding from the core RNAP particle. Scale bar, 140 Å. (**c**) Cryo-EM map of the complex in three orthogonal views related by 90° rotations. AlbA adopts a ‘butterfly-like’ arrangement, its LBDs (teal) diverging from the CC (brown) while its DBDs (purple) engage the promoter alongside σ⁷⁰ (pink). Labels mark the LBD, CC and DBD, the σ⁷⁰ non-conserved region (NCR) and the *palbA* promoter. The remaining colors follow the key beneath the panel. Magenta, density for KMR-28-27 within the LBD. (**d**) Close-up of the AlbA DBD (purple, with fitted model and density) engaging the promoter major groove and contacting the σ⁷⁰ NCR (pink). Residues at the AlbA:σ⁷⁰ interface (Gln4, Ser31, Ser34 and Leu40 on AlbA, with Glu154, Glu155, Arg157 and Asp160 on σ⁷⁰) and at the AlbA:DNA interface (Arg17, His20, His21, Arg33, Tyr38, Arg39, Glu61 and Leu62) are labeled. (**e**) Ribbon representation of the AlbA dimer bound to bent promoter DNA, colored as in (c), with the ‘switch’ and the ‘coupling helix’ labeled and KMR-28-27 in magenta.

Dimerization is largely driven by the antiparallel CC which crosses the DNA axis at about 60° such that each DBD inserts its α2N recognition helix (15-22), into the major groove of one half-site, with its N-terminal arm (4-6), wing (33-39) and residues (61-62) gripping the flanking phosphate backbone (**Fig. 6d**). Although the two protomers are related by a near-perfect 2-fold axis, their interactions are not completely symmetric, with Y38 reading a guanine on the template strand at positions −33 and −15. Sequence discrimination rests mainly on eight base pairs, with the guanines at −27 and −21 particularly important as their N7 and O6 atoms make a bidentate interaction with R17. Our SELEX motif (**Fig. 5a**) follows the chemistry of the interaction well since the two invariant positions correspond to the two guanines read at −27 and −33, while neither of the least conserved positions is read through its base, −32 contacting only the sugar-phosphate backbone and −29 lying outside the direct interface. Like other MerR-family regulators^31,32,36^, AlbA induces a substantial bending and underwinding of the promoter DNA which displays a kink of ∼59°. The 19-bp spacer is compressed to 53 Å, virtually identical to the 52.8 Å expected of a canonical 17-bp spacer, and the helical twist falls to 562° against 646° for ideal B-form DNA, an under-twisting of 84°. This distortion is not spread evenly along the spacer but converges on the central C:G pair at −24, the only position to depart from Watson-Crick geometry, where base-pair opening reaches 18.8° against 1.1 ± 2.6° for the other pairs. Together, bending and underwinding realign the −35 and −10 elements recognized by σ_4_ and σ_2_, respectively.

As in the RNAP-bound EcmrR complex^36^, we observed an interaction between the DBD nearer the -10 element and σ^70^ NCR. While the short stretch (E154-D160) on σ^70^ is a conserved interaction patch, the DBDs present structurally equivalent but chemically distinct interfaces compared to EcmrR (A30, Y31, N33, D35, F40 and T42 corresponding to AlbA S31, A32, S34, A35, L40 and N42). AlbA additionally recruits its first four amino acids for σ^70^ binding, resulting in an overall larger buried surface compared to EcmrR.

The transition from the LBD-close to the LBD-open state observed here required the remodeling of the ‘switch’ region, which, as shown in the isolated AlbA:PBD complexes, does not occur in response to PBD binding. Promoter binding appears therefore to remodel the ‘switch’ itself, providing a return path from the DNA-reading surface to the LBDs. Unlike in isolated AlbA dimers, the helical segments at the LBD N-terminus (α0, α1a and α1b) adopt here the same arrangement seen in monomeric AlbAS, with the additional extension of the α0 helix, which fuses N-terminally to α6N. The α6N-α0 helix (114-131), which immediately precedes the ‘switch’ is ideally poised to couple LBD and DBD movements as it contacts all structural domains. We refer to this helix as ‘coupling helix’ (**Fig. 6e**).

Specifically, it mediates cross-chain interactions with α3N (43-58) of the DBD with L116, V120 and L123 packing against A54 and Q57, while E117-H50 and E124-Q57 form carboxylate-imidazole and carboxylate-amide pairs. In addition to the CC, the α3N helix, and the (α6N-α0) ‘coupling helix’, contribute the dimerization interface (**Suppl. Fig. 22**). The identification of H50N and L120Q (corresponding to V120 in *K. oxytoca*) AlbA mutations in *K. pneumoniae* PBD-resistant isolates lends support to the key role of this hotspot^17,19^.

## Discussion

Albicidin and the cystobactamids are potent natural antibiotics targeting gyrase and resistance mechanisms have already been reported in *Klebsiella* strains including the human pathogen *K. pneumoniae* of the ESKAPE group^19,23^. Recently, we showed that resistance to C8-linked PBDs, a chemically distinct class, converges on the same determinant, the MerR-family transcription factor AlbA^19^. As AlbA is also present in other critical ESKAPE pathogens such as *P. aeruginosa* and *A. baumannii*, its relevance is likely general, and understanding how it orchestrates a resistance response offers guiding principles for the design of next-generation antibiotics able to evade it and of targeted inhibitors as resistance breakers.

This study allows us to put forward general principles for AlbA-mediated transcriptional regulation in response to its ligand effectors which differ from the common mechanism of MerR regulators. MerR-family members have been classified into metalloregulators, redox sensors, and drug-sensor subgroups^33^. AlbA belongs to the drug-sensor subgroups^33^ which also includes *Bacillus subtilis* BmrR^32^, BltR^34^ and Mta^34^, *Pseudomonas aeruginosa* BrlR^51^, *Escherichia coli* EcmrR^36^, and *Streptomyces* spp. TipA^34^. MerR-family members are typically repressors that trap RNAP at their operators in an inactive pre-initiation complex until effector binding at the LBDs is relayed allosterically to the DBDs, distorting the suboptimal 19/20-bp promoter spacer so as to realign the −35 and −10 elements for optimal RNAP engagement and formation of the competent initiation complex^34^. For AlbA, we propose a different mechanism which rests on a conformational equilibrium between differentially active conformations (**Fig. 7**). AlbA is predominantly in a partly autoinhibited state (LBD-close) in which the ‘switch’ region of each protomer crosses over, reaching the NTD end of the partner’s tunnel while the DBDs and CC remain mobile. This, possibly together with a minor active LBD-open/DBD-out state, affords a basal level of ligand-independent transcription. An equilibrium between ordered-disordered states and drug-induced transitions between these has also been reported in the related TipA system^52–54^. Transcription enhancement is then achieved in a ligand-selective manner by destabilizers like albicidin, but not PBDs, that act directly on the autoinhibited ‘switch’ conformation shifting the population to a larger fraction of competent states.

**Figure 7.**
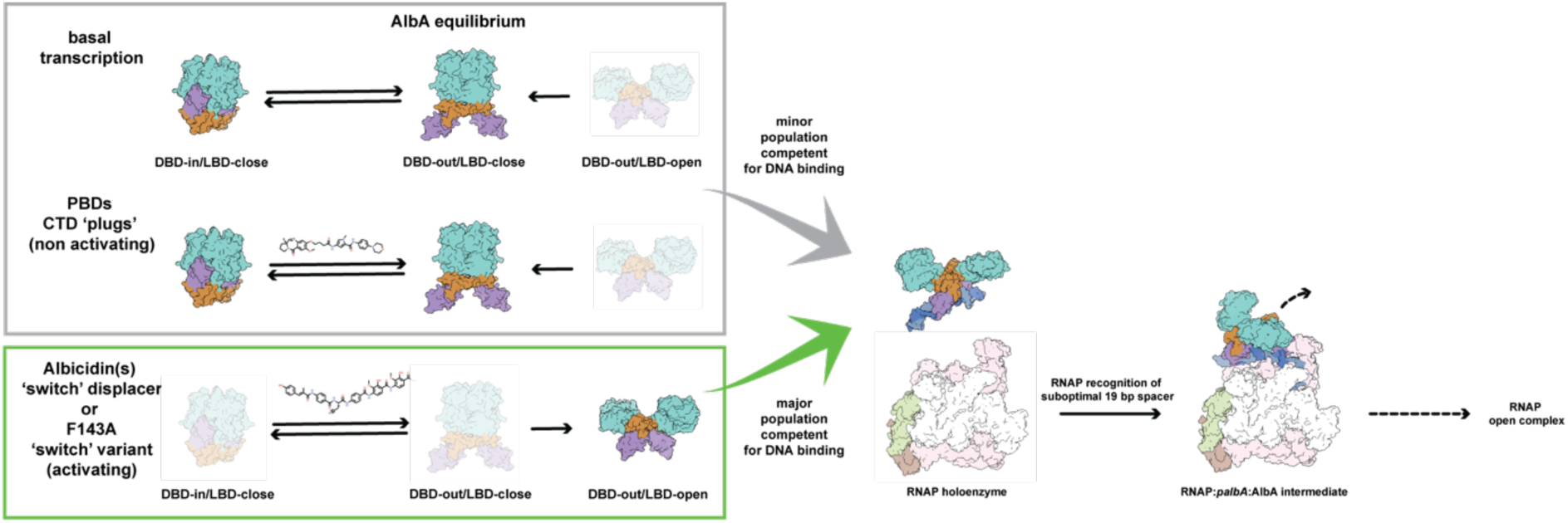
AlbA differs from conventional MerR-family regulators in modulating transcription by reweighting a pre-existing conformational equilibrium. AlbA interconverts between multiple states, dominated by LBD-close ones in which the DBDs are only partly accessible for DNA binding. In each panel the species drawn in pale color is the disfavored member of that equilibrium. The two ligand classes act differently on the equilibrium. PBD ‘plugs’ occupy the CTD end of the AlbA LBD tunnel without displacing the ‘switch’, leaving the protein in the ‘switch’-closed, wild-type-like state and the equilibrium unchanged. Albicidin spans the LBD tunnel and is incompatible with the ‘switch’-closed arrangement, displacing it and shifting the population toward the DBD-out/LBD-open conformer. The same behavior is recapitulated by the F143A ‘switch’ variant in the absence of albicidin. Upon binding to the *palbA* promoter, AlbA distorts the otherwise suboptimal 19 bp spacer between the −35 and −10 elements, leading to the RNAP:*palbA*:AlbA intermediate in which AlbA is seen in the DBD-out/LBD-open state. In a subsequent step, AlbA must leave to produce the RNAP open complex. AlbA is colored with the DBDs in violet, the coiled coil and ‘switch’ in brown and the LBDs in teal. RNAP subunits are in pastel shades and the two strands of the *palbA* duplex in dark/light blue.

The operational logic of AlbA follows that of the TipA system, which responds to thiostrepton and other antibiotics of its class with the translation of the long self-upregulating TipAL transcription factor and the short TipAS, its LBD alone. Interestingly, early studies indicated that TipAL is not a repressor and activates transcription whether thiostrepton is present or not, with thiostrepton further enhancing transcription^55^. This mirrors what we observe in our AlbA *in vitro* assay. In the cellular context this behavior, which is atypical within the MerR-family, can be rationalized by the presence of the short AlbAS/TipAS proteins, which depending on their ratio compared to the longer form, can act as negative regulators preventing uncontrolled amplification of downstream transcripts.

With respect to PBD binding, we have generalized here our earlier results (ref) showing that AlbAS binds two C8-linked PBD molecules in its central hydrophobic tunnel, with variable poses and nanomolar affinities making it an effective sequestering agent. The observation that PBDs bind AlbAS with comparable affinity to albicidin yet do not trigger activation suggests a strategy for overcoming AlbA-mediated resistance. Because the MCA1 aromatic moiety of albicidin is essential for transcriptional activation but also critical for DNA intercalation and antibacterial potency, modifying this region to reduce AlbA binding would likely impact activity against the DNA gyrase as well^39,56^. As non-enhancers, PBD derivatives could be developed as adjuvants to be co-administered with oligoarylamide antibiotics to counteract AlbA-mediated resistance. Our data show the carbinolamine form of PBDs binds within the AlbAS channel, suggesting the reactive imine responsible for cytotoxicity can be modified without disrupting AlbA binding. This opens the way to PBD dilactams or analogs with lower cytotoxicity as AlbA-targeted adjuvants that suppress resistance without antibacterial activity of their own. Moreover, compounds such as cystobactamids and coralmycins^57^, which share the linear, aromatic-rich scaffolds of albicidin, may also interact with AlbA, potentially expanding the scope of this resistance mechanism.

Overall, our results provide a mechanistic basis for AlbA-mediated resistance and reveal that PBDs can act as potent, non-activating AlbA binders, offering a blueprint for the rational design of AlbA inhibitors that could restore the efficacy of oligoarylamide derivatives and support the development of next-generation antibiotics.

## Methods

An extended version of the methods employed is available in the accompanying Supplementary Information file.

### Chemical synthesis

Synthetic procedures for KMR-04-154, KMR-04-160, KMR-04-161, KMR-04-163, KMR-04-165, KMR-04-177, KMR-04-184, KMR-04-186, KMR-14-14, KMR-28-27, KMR-28-31, KMR-28-32 and KMR-28-38 were described previously^15,19^. The of AF-CD-007 was also previously reported^58^. The synthesis and characterization of the PA-267-36, the KMR-14-14 dilactam is described in the Supplementary Information material.

### Protein expression and purification

Expression and purification of *K. oxytoca* His_6_-AlbAS (UniProt Q8KRS7, residues 1-221) has been described previously^19^. For AlbA expression, a pET-28a(+)-TEV plasmid encoding codon-optimized (for *E. coli* expression) *K. oxytoca* His_6_-AlbA (UniProt A0A9P0U153, residues 1-348) was obtained from Genscript (USA). The AlbA^F143A^ variant was generated by site-directed mutagenesis using the Q5 Site-Directed Mutagenesis Kit (NEB). Primers are reported in **Suppl. Table 6**. All constructs were verified by plasmid sequencing. *E. coli* RNA polymerase (RNAP) was encoded on the PVS10 plasmid obtained from the Addgene repository (Addgene plasmid #104398)^59^. *E. coli* σ^70^ factor was encoded on the PIA586 plasmid, obtained from the Addgene repository (Addgene plasmid #104399)^59^. Details of the purification protocols for the various proteins can be found in the Supplementary Information file. The RNAP(σ^70^):DNA:AlbA complex was reconstituted by incubating the individual components in a 1(2):1.5:5 ratio. The sample was then purified on a Superose 6 10/300 Increase (Cytiva) in S6 buffer (10 mM HEPES, 100 mM KCl, 5 mM MgCl_2_, 1 mM DTT, pH 7.5) and selected fractions were concentrated to about 5 mg/ml.

### Minimum Inhibitory Concentration (MIC) assay

MICs for each antibiotic were assessed by the broth microdilution method, following a modified protocol from Surani *et al*^19^. AlbAS expression was induced with 200 μM IPTG at OD_600_ = 0.1 and protein synthesis was induced at 20°C for 1 hour. After that, the culture was diluted to OD_600_ = 0.08-0.13 (approximated McFarland standard of 0.5) and used to prepare the microdilution plate according to the protocol, except for the presence of 200 μM IPTG in each well and the growth at 28°C to minimize protein misfolding. Experiments were carried out in triplicate.

### Equilibrium dissociation constant (*K*_D_) determination by fluorescence quenching

Ligand binding to AlbAS and to AlbA was followed by quenching of intrinsic protein fluorescence. A fixed protein concentration (50 nM) was titrated with ligand over approximately 0.1 nM to 10-50 μM, three replicate emission spectra were recorded per point, and intensities were taken at the emission peak determined from the lowest-ligand point (**Suppl. Table 1**). Because several dissociation constants approach or fall below the total binding-site concentration, the data were fitted to the Morrison quadratic equation for tight binding, which explicitly accounts for ligand depletion. The titration data were well described by a single binding transition, and the fitted parameter is therefore reported as an apparent (macroscopic) dissociation constant (*K*_D,app_) for ligand binding to AlbAS rather than a site-resolved constant. We acknowledge the use of Claude Science for the analysis of fluorescence quenching data.

### **X-** ray crystallography

AlbAS at 30 mg/mL was mixed with each ligand from a 100% DMSO stock to a nominal 3-5-fold molar excess and 1-2% final DMSO, incubated overnight at 4 °C and clarified by centrifugation. Crystals were grown by sitting-drop vapor diffusion, in 1.0-2.0 M ammonium sulfate for the KMR-04-154, KMR-04-163, KMR-04-177 and KMR-28-27 complexes and in 0.15 M ammonium sulfate, 0.1 M MES pH 6.5, 25% PEG 4000 for the KMR-04-161 and KMR-04-165 complexes. Crystals were cryoprotected with 2 M sodium malonate or 25% (v/v) ethylene glycol and cryo-cooled in liquid nitrogen. Data were collected remotely at the ESRF (Grenoble) and at Diamond Light Source (Didcot) and processed with Autoproc^60^ and Dials^61^. Structures were solved by molecular replacement in Phaser^62^ using the AlbAS:KMR-14-14 complex (PDB 8RKY) as search model, built in Coot^63^ and refined with Refmac5^64^ or Phenix.refine^65^, with ligand restraints from Grade2^66^. Data collection and refinement statistics are given in **Suppl. Table 2**.

### AI-driven model generation of the AlbA dimer

Dimer models were generated with AlphaFold2 (v1.5.2)^40^ and AlphaFold3^42^ from the *K. oxytoca* AlbA sequence (UniProt A0A9P0U153, residues 1-348) at default settings, and with Chai-1 (v0.6.1)^43^ under intermolecular distance restraints taken from the experimental cryo-EM LBD dimer. A pool of 2,000 dimeric models sampling alternative relative domain orientations was then generated with Boltz-2 (v2.2.1)^45^, comprising 1,000 template-free models, 500 restrained to the experimental apo cryo-EM structure (LBDs only) and 500 restrained to the Chai-1 model. All predictions used the same locally generated ColabFold (v1.6.2)^41^ MSA, 10 recycling steps and 200 diffusion sampling steps, with independent random seeds and otherwise identical inference settings. Models were screened for coiled-coil integrity and those whose two helices had dissociated were rejected, leaving a final pool of 1,985. Restraint lists, hardware and full inference settings are given in the Supplementary Information.

### Small angle X-ray scattering (SAXS)

SEC-SAXS data for AlbA (100 μL at 10 mg/mL in 50 mM Tris-HCl pH 7.5, 150 mM NaCl, 1 mM DTT) were collected at beamline P12 (DESY, Hamburg) on a Pilatus 6M detector at 0.124 nm, with elution from a Superdex 200 Increase 10/300 column at 0.5 mL/min. Batch data for AlbA^F143A^ (1.25 mg/mL in the same buffer) were collected at BM29 (ESRF, Grenoble) on a Pilatus3 2M detector at 0.099 nm. Both experiments were performed at 20 °C and all processing used the ATSAS^44^ suite, with CHROMIXS for the SEC-SAXS data and PRIMUS for buffer subtraction of the batch data, AUTORG for *R*_g_ and *I*(0), GNOM for the pair distance distribution, DATVC for molecular weight and DATCLASS for shape classification. Theoretical scattering was computed with CRYSOL for single models and with FFMAKER for ensembles, the latter applying a common hydration-shell contrast to every model. The solvent electron density was fixed at 0.334 e Å^-3^ and the shell contrast set to 0.060 e Å^-3^ for AlbA and 0.030 e Å^-3^ for AlbA^F143A^, each chosen as the minimum of a contrast scan over ensemble fit quality. Mixtures were selected from the model pool independently by non-negative least squares (NNLSJOE) and by a genetic algorithm (GAJOE). Ensemble composition was assessed over all independent fits of each protein, 26 for AlbA and 10 for AlbA^F143A^, which combine Monte Carlo replicates, repeated runs from independent seeds, both selection algorithms and a jackknife in which each selected conformer was withheld in turn. Each model was classified on two coordinate-derived axes, the fraction of DBD surface buried in the dimer, computed over residues 5-70 of both subunits and with values above 0.341 defining DBD-in, and an inter-LBD contact count. Boundaries were chosen from the bimodal pool distributions. We acknowledge the use of Claude Science for the SAXS classification analysis.

### Cryo-EM

Grids for AlbA were prepared from freshly purified protein at 2 mg/mL applied to glow-discharged UltrAuFoil grids and plunge-frozen in liquid ethane with an EM GP2 (Leica). For the AlbA:KMR-14-14 and AlbA:KMR-28-27 complexes the protein was first incubated with a 10-fold excess of ligand (0.6% final DMSO) for 3 h at 4 °C. For the RNAP(σ^70^):DNA:AlbA complex, the purified complex at 5 mg/mL was supplemented with additional AlbA, KMR-28-27 and CHAPSO and vitrified on Quantifoil C-Au grids with a Vitrobot (ThermoFisher). The AlbA and AlbA:KMR-14-14 datasets were collected on a Krios G3i operating at 300 kV with a K3 camera and BioQuantum energy filter at 0.54 Å/px in super-resolution mode, giving 16,987 and 24,100 micrographs at a total dose of 70 e^-^/Å^2^. The AlbA:KMR-28-27 dataset was collected on a Glacios operating at 200 kV with a Falcon 4i camera at 1.2 Å/px, giving 5,239 micrographs at 60 e^-^/Å^2^. The RNAP(σ^70^):DNA:AlbA(KMR-28-27) dataset was screened on a Glacios and collected at eBIC on a Titan Krios at 300 kV with a K3 camera and BioQuantum filter at 0.825 Å/px, giving 24,521 micrographs at 40 e^-^/Å^2^. All datasets were processed in CryoSPARC^67^ following near-identical pipelines of patch motion correction, patch CTF estimation, template picking, 2D classification, *ab initio* reconstruction, heterogeneous refinement and reference-based motion correction, as schematized in **Suppl. Fig. 9, 12, 13** and 18. For the RNAP complex, 3D classification was used to isolate the classes carrying AlbA density, giving a consensus reconstruction at 2.43 Å, and subtraction of the RNAP core followed by masked local refinement gave a focused AlbA reconstruction at 3.03 Å (gold-standard FSC, 0.143 criterion). Starting models were PDB 8RKY for apo AlbA and the AlbA:PBD complexes and PDB 6XL5 for the RNAP core, with AlphaFold2^40^ predictions for the AlbA DBD and CC. Remaining regions were built manually in Coot^63^ and models were refined with a combination of Servalcat^68^, Phenix.refine^65^ and Isolde^69^, with ligand restraints from Grade2^66^.

### SELEX

An 82 bp double-stranded library carrying a 42 bp random region flanked by 20 bp primer regions with BamHI and EcoRI sites was prepared from a synthetic oligonucleotide (Eurogentec) by Klenow extension and amplified with Vent polymerase. The library (500 ng) was incubated with AlbA at protein to DNA ratios of 5:1, 20:1 and 40:1 under progressively more stringent conditions across rounds, up to 500 mM NaCl and 500 ng/μL competitor ssDNA, and complexes were resolved on 6% polyacrylamide gels in 0.5×TBE. Shifted bands were excised, DNA was eluted overnight in diffusion buffer, purified and amplified for the next round. After seven rounds the selected library was cloned into pUC19, 95 white colonies were sequenced by direct colony sequencing (Eurofins Genomics) and motif analysis was performed with the MEME Suite^70^.

### Fluorescence polarization (FP)

3′-TAMRA-labeled oligonucleotides (Merck) were annealed with their unlabeled complements, incubated for 1 h at 4 °C with increasing concentrations of AlbA and read on a TECAN Spark in black 384-well plates. Polarization values were baseline-corrected against the DNA-only signal and each titration was fitted to a one-site binding model incorporating a linear nonspecific term, so that reported *K*_D_ values describe the saturable component only. Individual wells whose robust standardized residual exceeded three were excluded, 10 of 249 measurements, with the residual scale estimated from a robust fit to the complete dataset so that the threshold was not itself influenced by the outlying points. The three *palbA* titrations acquired under matched conditions were adequately described by a single nonspecific slope (F-test, P = 0.97) and were therefore fitted globally with that slope shared, whereas AlbA^F143A^ and the *pSELEX* duplex were fitted individually. Differences between conditions were assessed by F-tests on nested models, and confidence intervals on *K*_D_ were obtained by profile likelihood. Regression was performed in SciPy using the trust region reflective algorithm with all parameters constrained to be non-negative. We acknowledge the use of Claude Science for the analysis of FP data.

### *In vitro* transcription assay

The 178 bp and 176 bp templates carrying the *palbA* and *palbA*-17 promoters were amplified from the corresponding plasmids with Vent polymerase and gel-purified, *palbA*-17 having been generated from *palbA* by site-directed mutagenesis. Reactions contained RNAP holoenzyme, template DNA and AlbA in TA buffer (40 mM Tris-HCl, 150 mM NaCl, 2 mM MgCl_2_, 1 mM DTT, pH 7.5), with ligands pre-incubated with AlbA before addition. Transcription was initiated with 500 μM rNTPs, allowed to proceed for 2 h at 37 °C and stopped with RNA loading buffer. Products were resolved on 10% urea-PAGE, stained with ethidium bromide and imaged on a Uvitec NineAlliance station.

## Data availability

X-ray crystallography structures are freely accessible on the Protein Data Bank with codes 9I3G, 9IC9, 9I4J, 9I9R, 9QBM, 32HM. Cryo-EM maps are freely accessible at the EMDB repository with codes EMD-57289, EMD-58346, EMD-57335, EMD-57336, EMD-57337.

Cryo-EM structures are freely accessible on the Protein Data Bank with codes 29QP, 31ER, 29RP.

## Acknowledgments

We are grateful to staff scientists at the ESRF (beamlines ID30B and BM29), DLS (I04), and DESY (P12) synchrotrons for their support during measurements. Scientists at the UK electron Bio-Imaging Centre (eBIC), the King’s College London Centre for Ultrastructural Imaging (KCL-CUI), the London Consortium for Cryo-EM (LonCEM), and the University of Padova Cryo-EM facility within the Department of Biomedical Sciences (DSB-CEM) are also acknowledged for their help. LonCEM is supported by the Wellcome Trust grant 206175/Z/17/Z and its partner institutes. DSB-CEM is supported by the ’Dipartimenti di Eccellenza (2023-2027)’ grant by the Italian Ministry of University and Research (MUR).

Work in the group of RDS was supported by the Deutsche Forschungsgemeinschaft (DFG, German Research Foundation) as part of the DFG-funded graduate school RTG 2473 (Bioactive Peptides, project number 392923329) to L.K. and R.D.S. We thank Tam Bui of the KCL Molecular Characterisation Facility for her help with the SEC-MALS experiment. We are also grateful to Julien Bergeron and Andrew Goodale for useful discussions in the early stages of the project. M.D.P. is a PhD student in the group of R.A.S. funded by MUR.

## Author contributions

J.M.S., K.M.R, and R.A.S. conceived the study. M.D.P. and R.A.S. designed the experiments. M.D.P. and A.G. carried out the experiments. P.A., K.M.R., L.K., R.D.S. contributed essential reagents. M.D.P., A.G., M.L. G.M., R.A.S. analyzed the data and interpreted the results. M.D.P. and R.A.S. wrote the paper with contributions from all authors.

---

**Suppl. Fig. 1.**
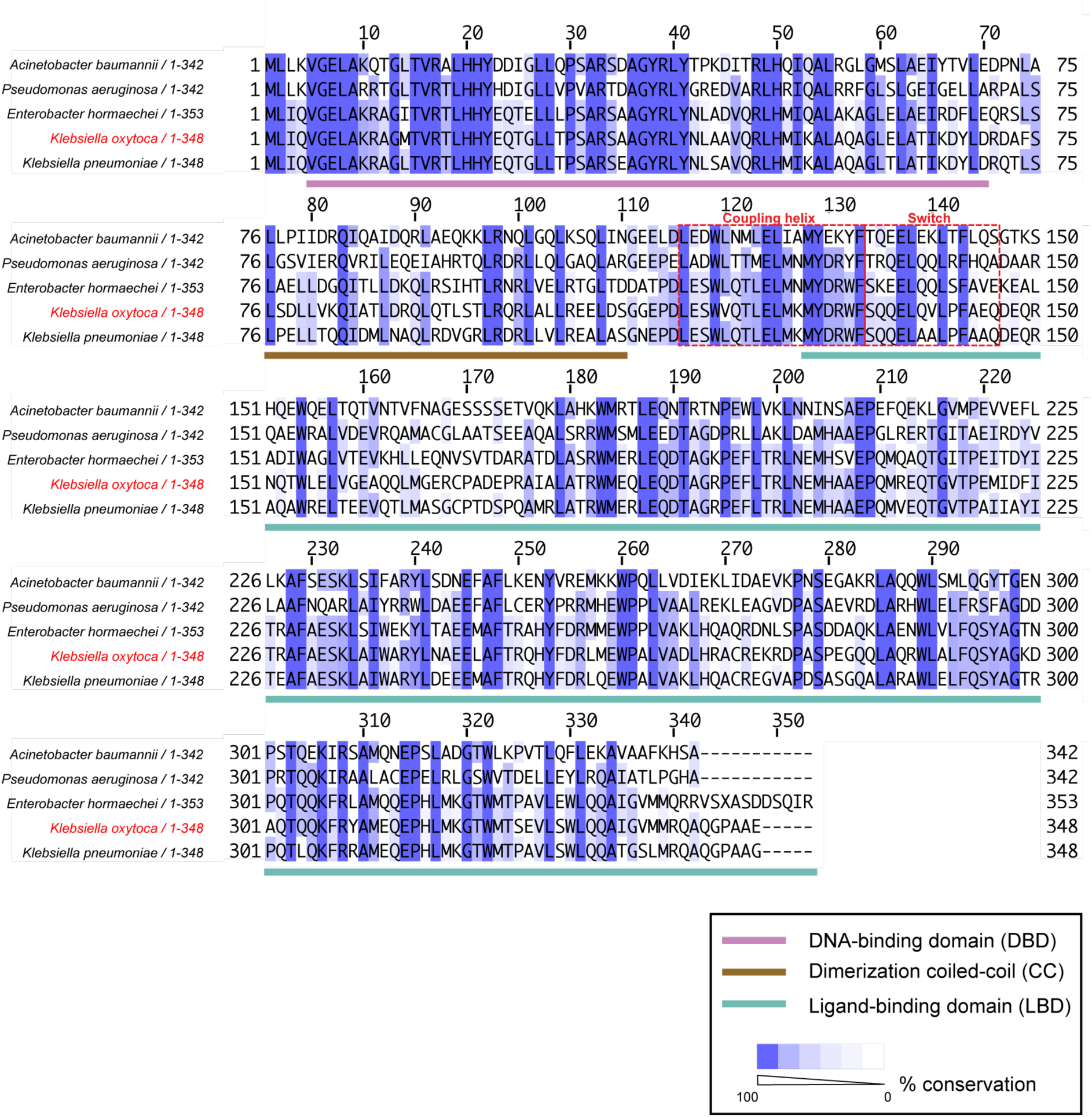
AlbA orthologues are present across the Gram-negative ESKAPE pathogens. Multiple sequence alignment of full-length AlbA from *Acinetobacter baumannii* and *Pseudomonas aeruginosa* (342 residues), from *Klebsiella oxytoca* and *Klebsiella pneumoniae* (348 residues) and from *Enterobacter hormaechei* (353 residues). The *K. oxytoca* sequence studied here is labeled in red. Columns are shaded by conservation as indicated in the key. Colored bars beneath the alignment mark the domain boundaries in *K. oxytoca* numbering, the DNA-binding domain (DBD), the dimerization coiled coil (CC) and the ligand-binding domain (LBD). All proteins share this MerR-family architecture. Red dashed boxes mark the ’coupling helix’ and the ’switch’ region, which we show to be important for ligand-dependent transcriptional activation (see main text).

**Suppl. Fig. 2.**
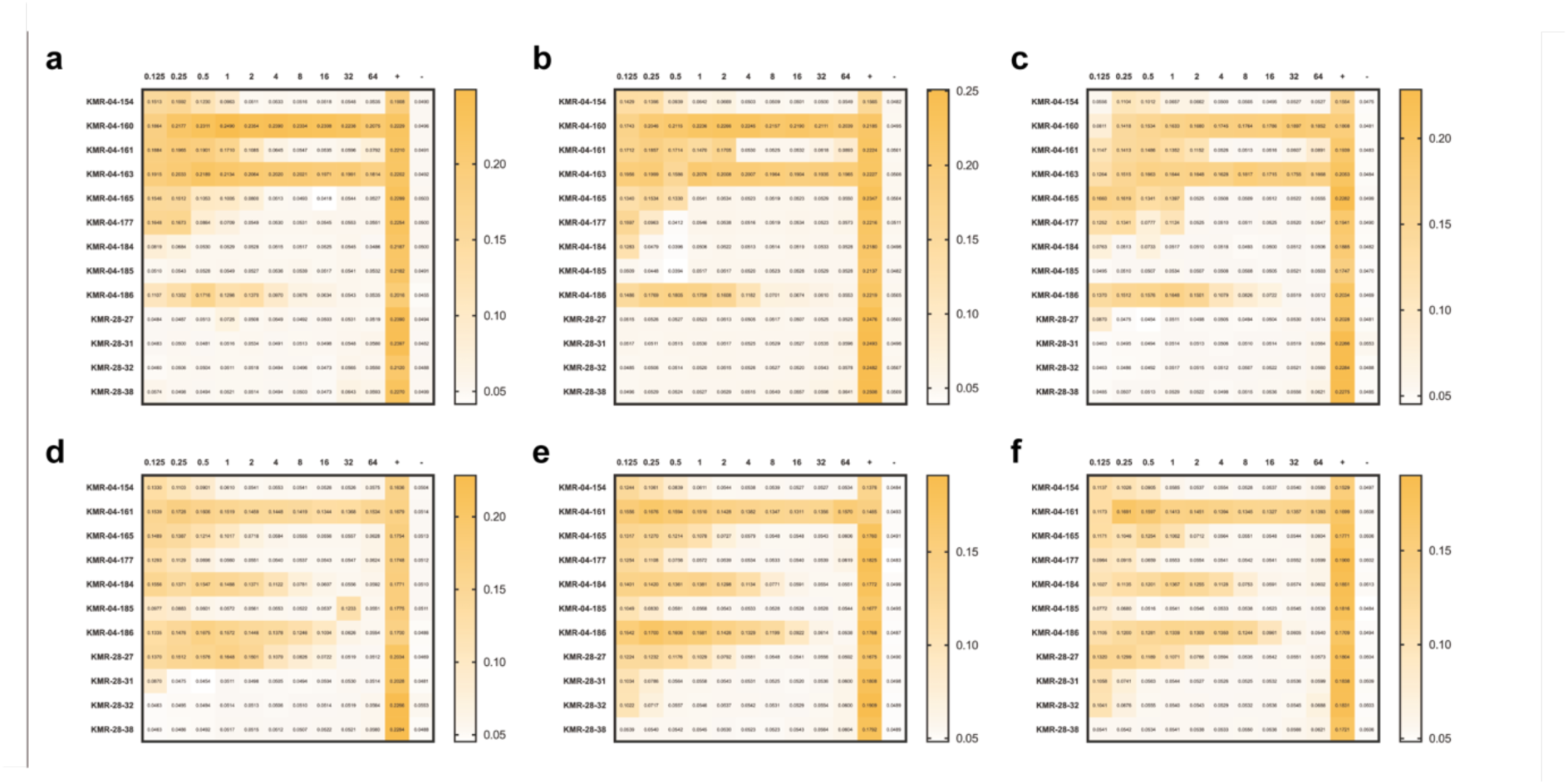
Minimum inhibitory concentration (MIC) determination for C8-linked PBD conjugates against *E. coli* in the absence and presence of AlbAS-induced resistance. MIC values were determined for the C8-linked PBD conjugates described in Fig. 1f of the main text using a broth microdilution assay with *E. coli* BL21(DE3) carrying an inducible plasmid expressing *K. oxytoca* AlbAS. Compound concentrations ranged from 0.125 to 64 µg/mL; MIC values recorded at either limit of this range are bounds rather than point estimates, and fold-shifts derived from them are correspondingly bounded. Cell growth was quantified by absorbance and is represented as a color gradient from white (no growth) to yellow. **(a–c)** Triplicate MIC experiments performed under non-inducing conditions. Under these conditions, most compounds displayed strong growth-inhibitory activity (MIC < 2 µg/mL), except for KMR-04-160 and KMR-04-163, which exhibited MICs > 64 µg/mL. **(d–f)** Triplicate MIC experiments performed under AlbAS-inducing conditions, restricted to compounds that exhibited a MIC below 64 µg/mL in panels a–c. The positive control column (+) contains bacterial culture without any PBD compound and serves as a reference for uninhibited growth. The negative control column (–) contains neither PBD compound nor bacteria and serves as a blank reference for background absorbance. MIC values reported in Fig. 1g of the main text represent the lowest compound concentration at which the optical density at 600 nm (OD_600_) is below 0.06 AU under the assay conditions.

**Suppl. Fig. 3.**
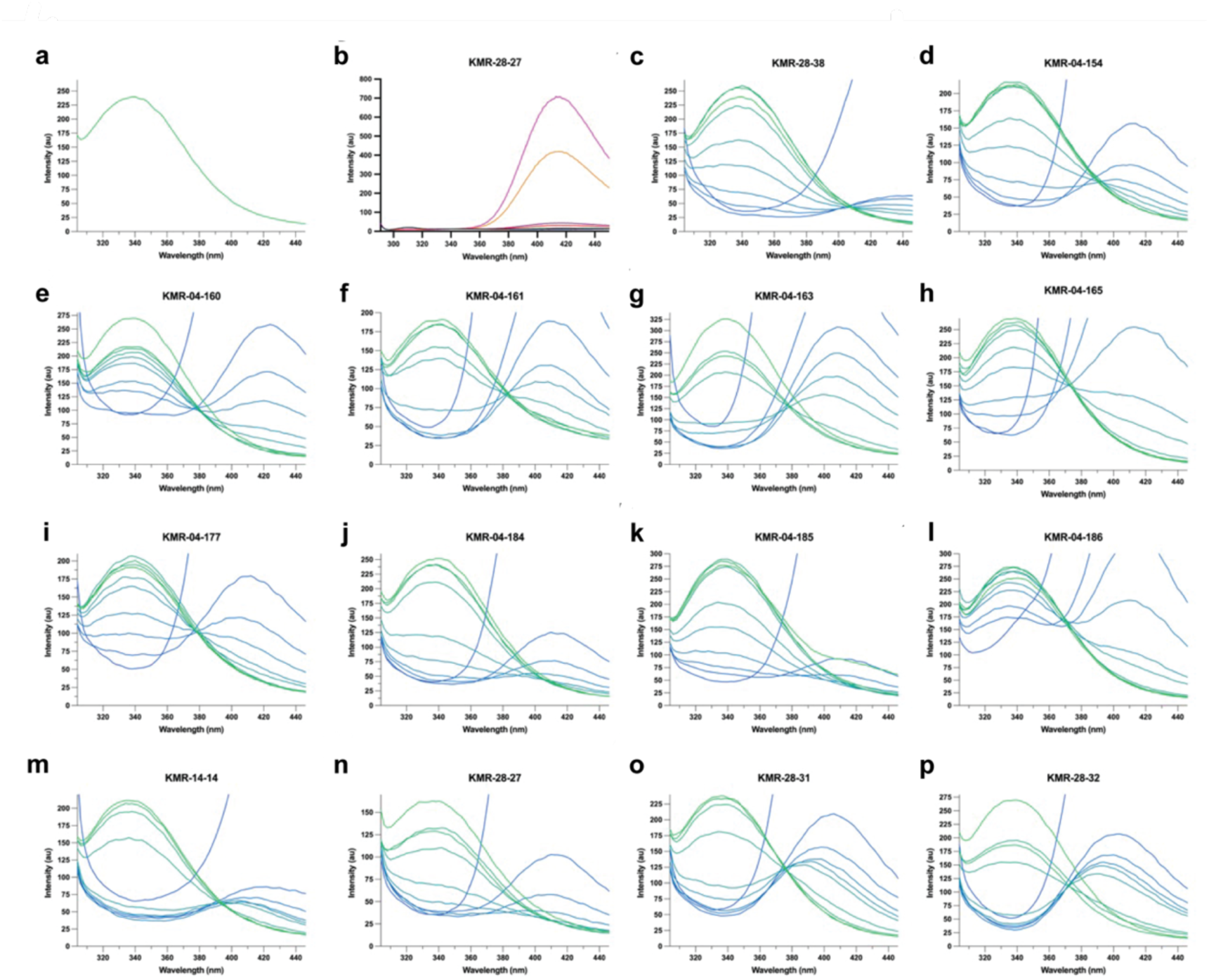
Fluorescence emission spectra of AlbAS upon titration with C8-linked PBD conjugates. **(a)** Intrinsic fluorescence emission spectrum of AlbAS (50 nM) recorded between 305 and 450 nm following excitation at 290 nm, reflecting the tryptophan-dominated emission of the unliganded protein with a peak around 340 nm. **(b)** Fluorescence emission spectra of KMR-28-27 at concentrations ranging from 0.001 to 20 µM, recorded under identical conditions. As exemplified by this compound, PBDs display a prominent intrinsic emission peak around 418 nm, confirming that the compounds themselves contribute significantly to the fluorescence signal at elevated concentrations. **(c–p)** Fluorescence emission spectra of AlbAS (50 nM) recorded upon stepwise titration with the indicated PBD ligands. Spectra progress from green (lowest ligand concentration) to blue (highest ligand concentration). Ligand binding causes quenching of the tryptophan emission peak at ∼340 nm, which was used to derive equilibrium dissociation constants. At higher ligand concentrations, the intrinsic PBD emission around 420 nm becomes increasingly apparent, overlapping with and partially masking the quenching signal and thereby limiting the useful concentration range for quantitative analysis. All experiments were performed in triplicate; representative curves from a single experiment are shown.

**Suppl. Fig. 4.**
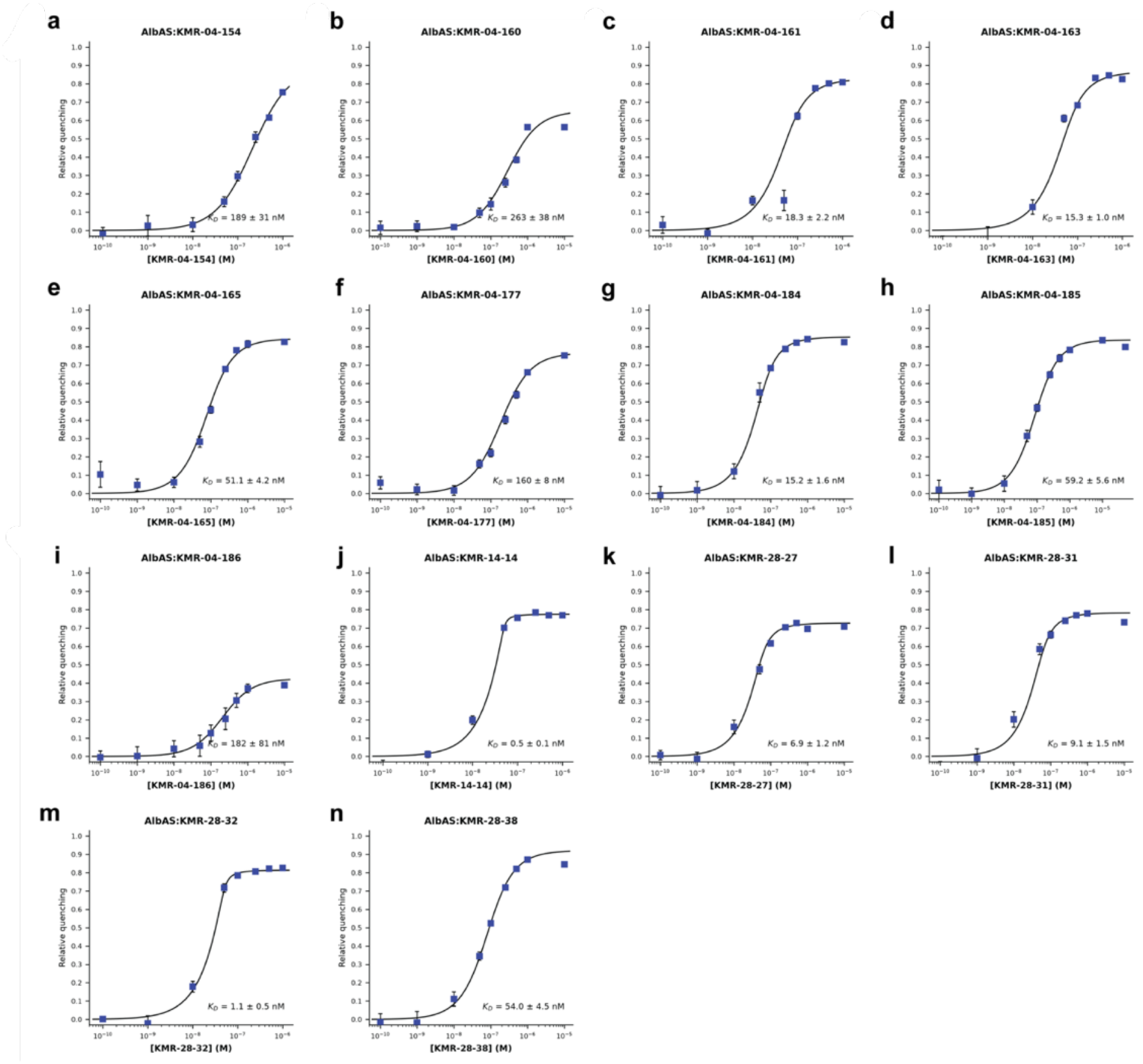
Apparent equilibrium dissociation constants for C8-linked PBD conjugates binding to AlbAS. (**a**–**n**) Binding curves for each indicated AlbAS:PBD complex, derived from the concentration-dependent quenching of AlbAS intrinsic fluorescence upon titration with the respective ligand (see **Suppl.** Fig. 3). The y-axis reports relative quenching, defined as the fractional decrease in tryptophan emission intensity measured at the wavelength of maximal emission (∼334–340 nm, determined independently for each titration series) relative to the fitted unbound-protein fluorescence (F_0_) and the fitted total quenching amplitude (ΔF). The x-axis shows total ligand concentration on a logarithmic scale. Data points (blue squares) represent the mean ± standard deviation of three independent experiments. Solid lines represent non-linear least-squares fits performed in Python (SciPy) to the quadratic (Morrison) equation for tight-binding, single-site equilibria, which explicitly accounts for depletion of free ligand by the bound protein. The ligand concentration range used for fitting was restricted to exclude high-concentration points at which intrinsic PBD fluorescence interfered with the quenching signal.

**Suppl. Fig. 5.**
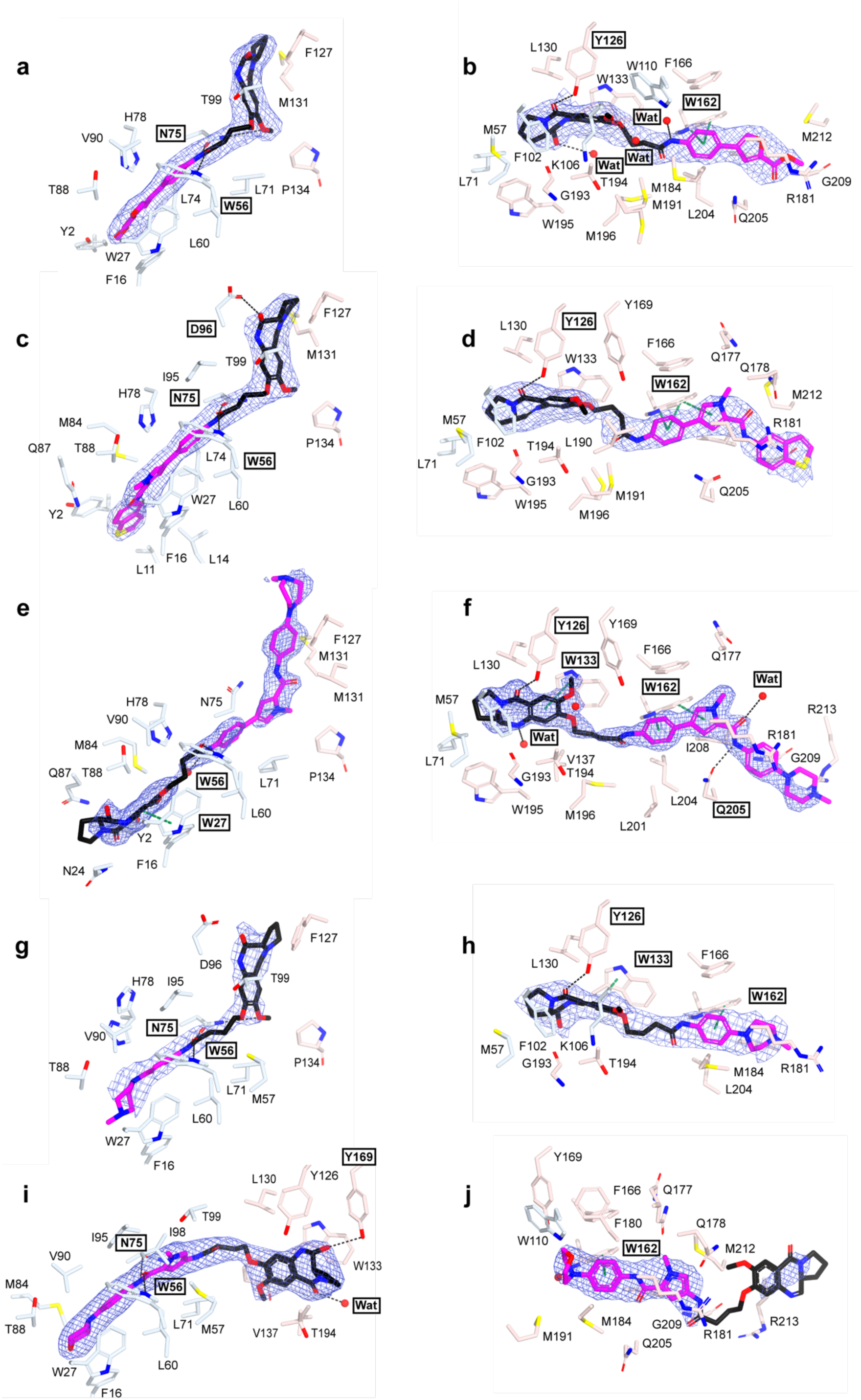
Protein–ligand contacts across five AlbAS–PBD complexes. Close-up views of the two ligand-binding sites of AlbAS for each of five compounds. Electron density maps (2*mF*_o_ – *DF*_c_) for the ligands are represented as a blue mesh at the 1σ level. Panels are paired by structure, with the N-terminal domain (NTD) site on the left and the C-terminal domain (CTD) site on the right: (**a**, **b**) KMR-04-154 (PDB 9I3G, 2.38 Å); (**c**, **d**) KMR-04-163 (9I9R, 2.58 Å); (**e**, **f**) KMR-04-165 (9IC9, 1.60 Å); (**g**, **h**) KMR-04-177 (9QBM, 2.95 Å); (**i**, **j**) KMR-28-27 (32HM, 2.56 Å). All ten panels share one orientation, derived from the ligand-binding channel of the superposed structures, and one scale; the NTD ligand therefore appears diagonal because the channel is bent at the domain interface. The C8-linked PBD core is colored black and the variable tail magenta. Protein side chains are shown as sticks and colored by domain, NTD in blue and CTD in red. Hydrogen bonds are drawn as black dashed lines and π–π stacking interactions as green dashed lines. Residues involved in these interactions are highlighted. Ordered water molecules bridging ligand and protein are shown as red spheres (Wat). The PBD imine group is present in its hydrated carbinolamine form in eight of the ten sites; it is not modified in the CTD sites of KMR-04-163 (d) and KMR-28-27 (j). Interactions for KMR-04-161 (PDB 9I4J, 1.74 Å) are given in Figure 2e**, f** of the main text.

**Suppl. Fig. 6.**
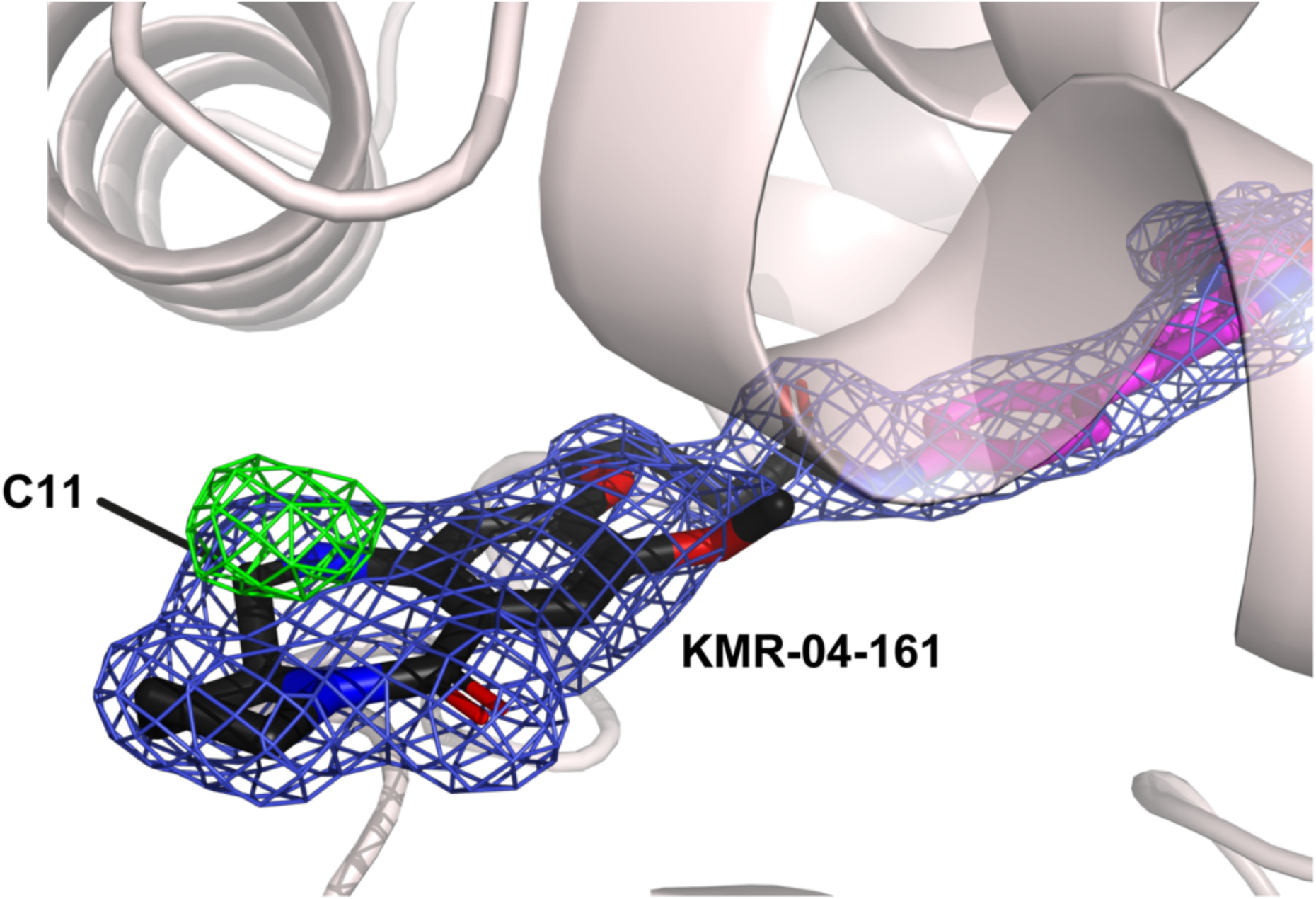
Many PBD ligands are bound in their hydrated carbinolamine form. The KMR-04-161 ligand shown here as an example was refined in its imine form. Sticks for the PBD core carbons are in black and the R-group tail carbons in magenta; nitrogen blue, oxygen red. In blue and green are shown the 2*mF*_o_-*DF*_c_ (1σ level) and the *mF*_o_-*DF*_c_ (+3σ level), respectively. The unmodeled positive difference peak above the C11 atom is consistent with its hydroxylation and is seen in several AlbAS-bound PDBs ligands studied here.

**Suppl. Fig. 7.**
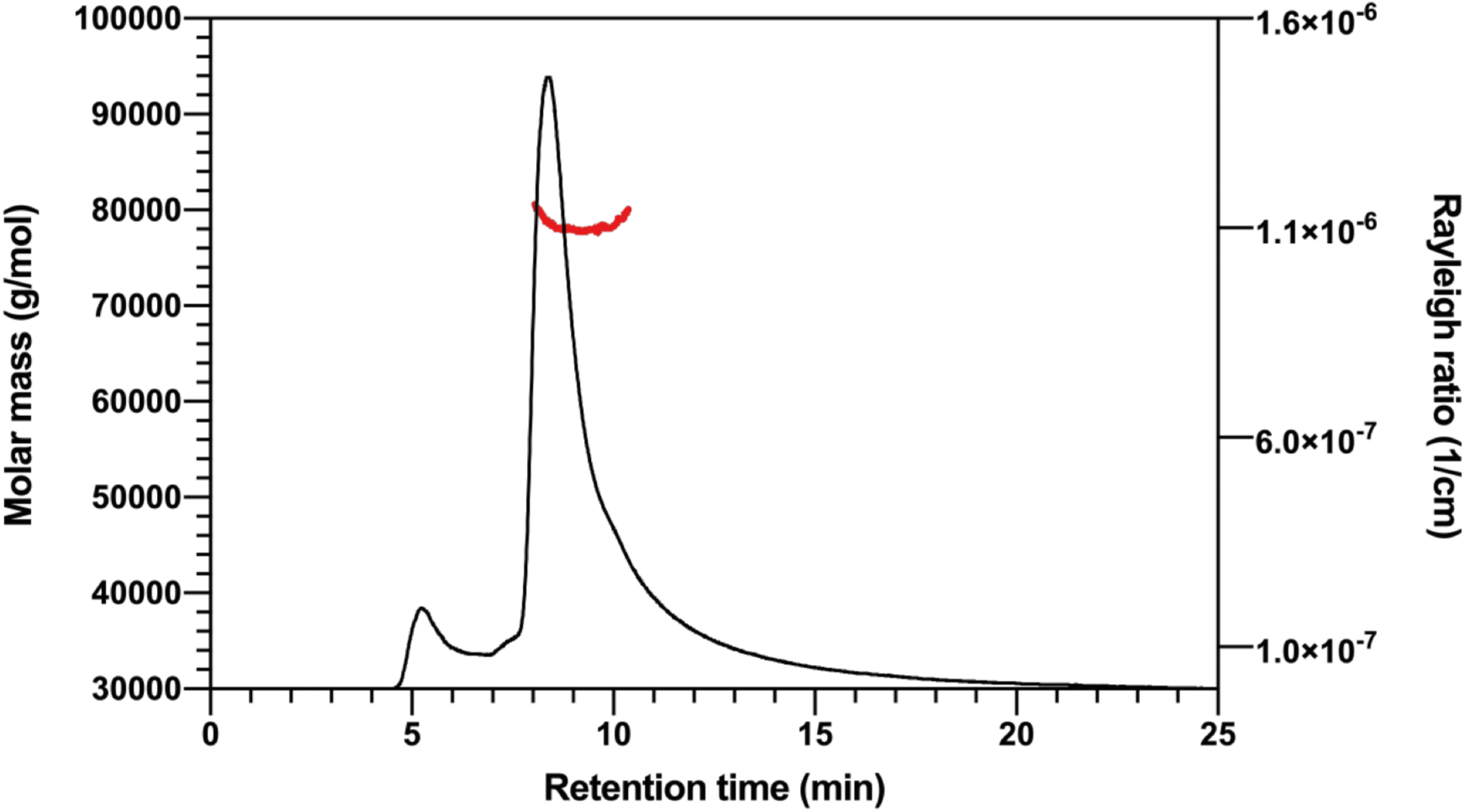
AlbA is a dimer in solution. SEC-MALS elution profile of AlbA confirming that the protein is a dimer in solution. The calculated MW of the AlbA monomer (UniProt A0A9P0U153_KLEOX) is 40082.76 Da.

**Suppl. Fig. 8.**
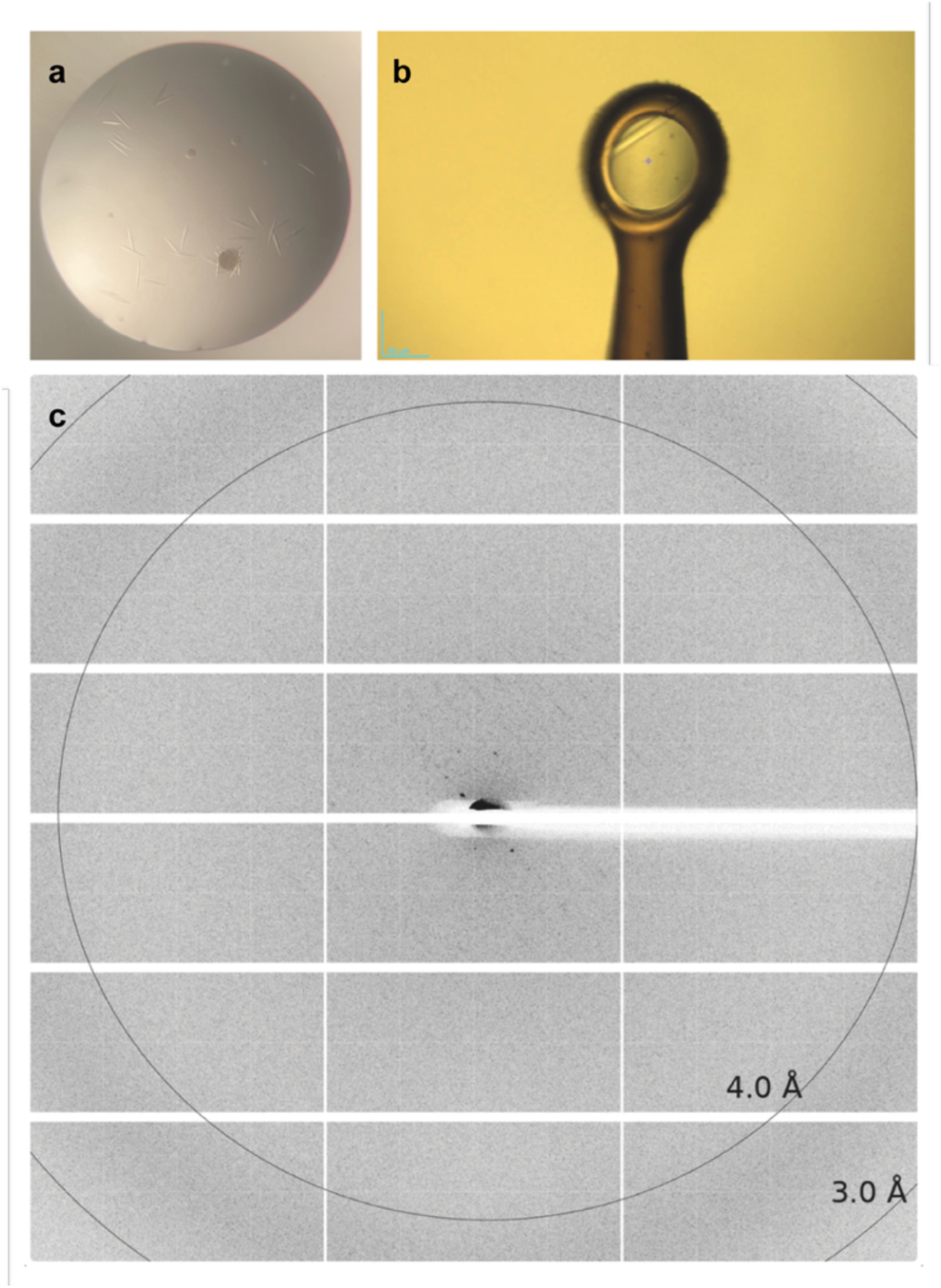
AlbA crystals diffracted X-rays poorly. **(a)** Crystals of AlbA from *Klebsiella oxytoca* were grown using the vapor diffusion technique at 20°C in 0.2 M KSCN, 20% PEG3350, 0.1 M Bis-tris propane pH 7.5. (**b**) For data collection, single crystals were cryoprotected with the reservoir solution enriched with with 25% (v/v) ethylene glycol and mounted on cryoloops. (**c**) A few diffraction spots at about 10 Å resolution are visible. X-ray diffraction data were measured at the ESRF (Grenoble, France) beamline ID23-2.

**Suppl. Fig. 9.**
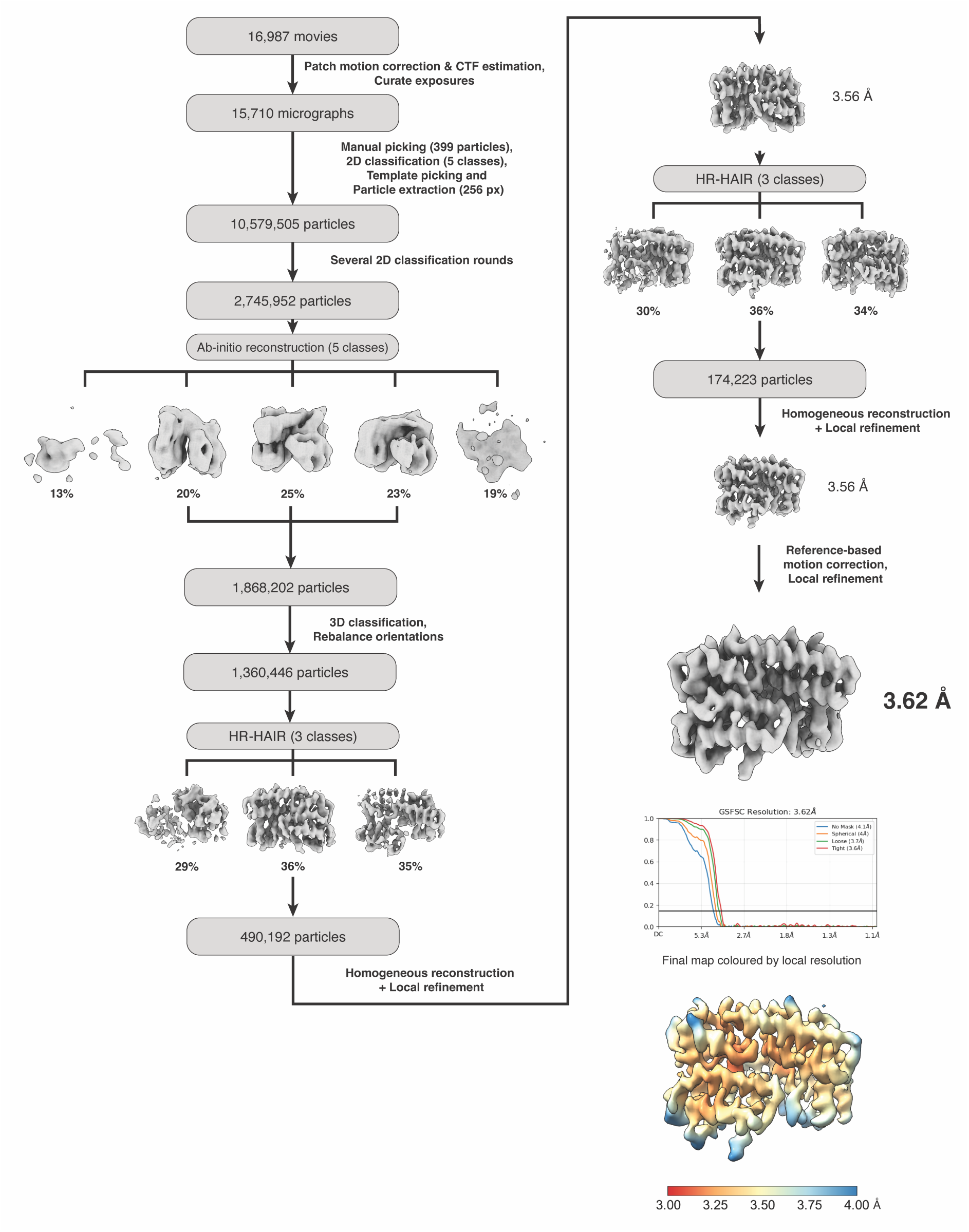
Cryo-EM data-processing workflow for AlbA. 16,987 movies were subjected to patch motion correction and patch CTF estimation, and exposures were curated to yield 15,710 micrographs. An initial set of 399 manually picked particles was classified into 5 2D classes to generate templates, and template-based picking with extraction in a 256-pixel box gave 10,579,505 particles, reduced to 2,745,952 by several rounds of 2D classification. *Ab initio* reconstruction into 5 classes, shown as density maps, yielded three classes with interpretable protein features, which were combined to give 1,868,202 particles. 3D classification with orientation rebalancing gave 1,360,446 particles, which were subjected to high-resolution heterogeneous ab-initio reconstruction (HR-HAIR) into 3 classes, and the best class of 490,192 particles was taken forward to homogeneous reconstruction and local refinement at 3.56 Å. A second round of HR-HAIR into 3 classes, followed by homogeneous reconstruction and local refinement of the best class of 174,885 particles, gave a map at 3.56 Å. Reference-based motion correction and a final local refinement produced the final reconstruction at 3.62 Å. The lower right panels show gold-standard FSC curves for the final refinement, calculated with no mask and with spherical, loose and tight masks, where the horizontal line marks the 0.143 threshold. The final map is also shown colored by local resolution over the range 3.00–4.00 Å.

**Suppl. Fig. 10.**
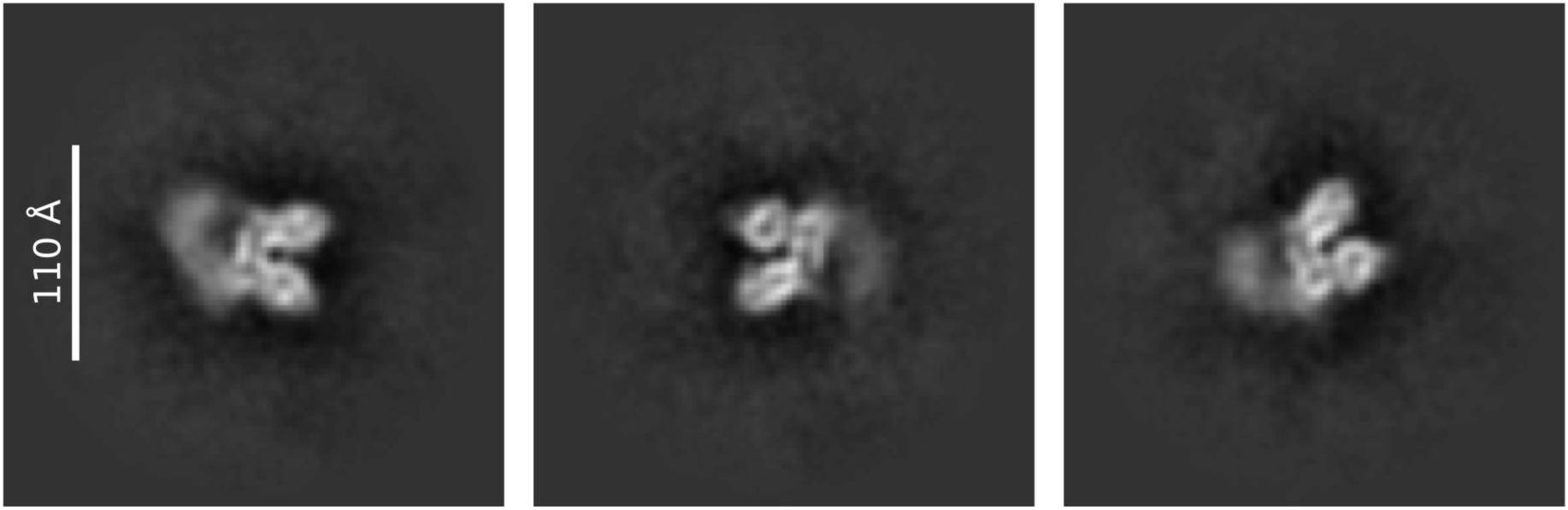
The N-terminal DNA-binding domains (DBDs) and dimerization coiled coils (CCs) are mobile relative to the ligand-binding domains (LBDs). Representative 2D class averages of AlbA particles. Class in which density attributable to the DBDs and CCs is present but smeared, consistent with a range of positions relative to the ordered LBD dimer. The smeared density indicates that the CC and DBD module are present in the particles but conformationally heterogeneous. Scale bar, 110 Å.

**Suppl. Fig. 11.**
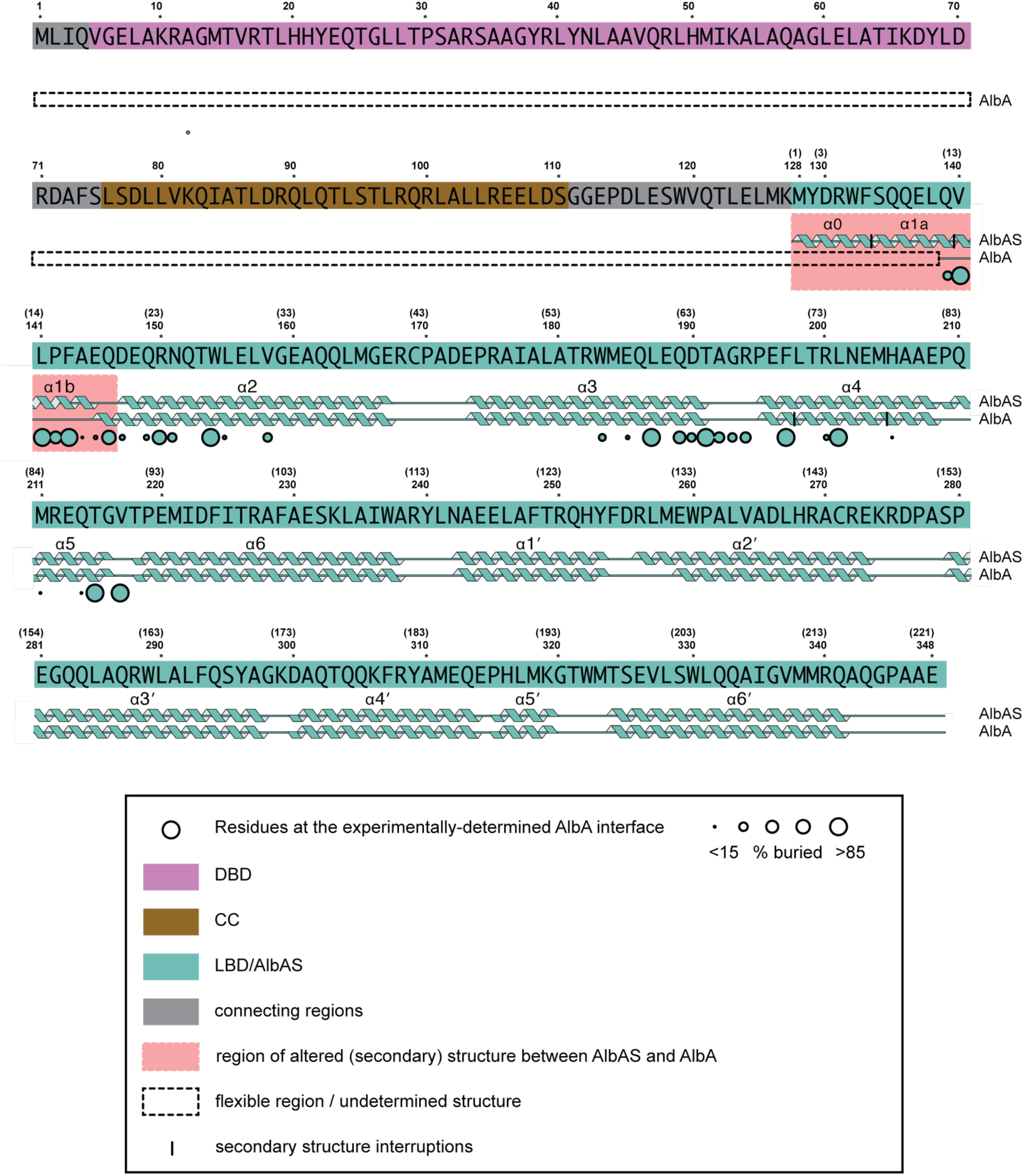
*K. oxytoca* AlbAS/AlbA secondary structure and AlbA interaction contact map. The AlbA sequence is shown in blocks with AlbAS numbering in parentheses, the two schemes differing by an offset of 127 amino acids. The different domains are color-coded as indicated in the legend. Secondary structure, from crystallography for AlbAS and from cryo-EM for AlbA, is drawn beneath each block, with dashed outlines and vertical bars as defined in the legend. LBD residues involved in the observed interface are highlighted by circles whose radius is proportional to the percentage of buried surface. The regions highlighted in red are those in which the secondary structure differs between the two forms, with α0 and α1a disordered and α1b non-helical in AlbA, and they coincide with the cluster of interface residues that forms the reciprocal ‘switch’ arm.

**Suppl. Fig. 12.**
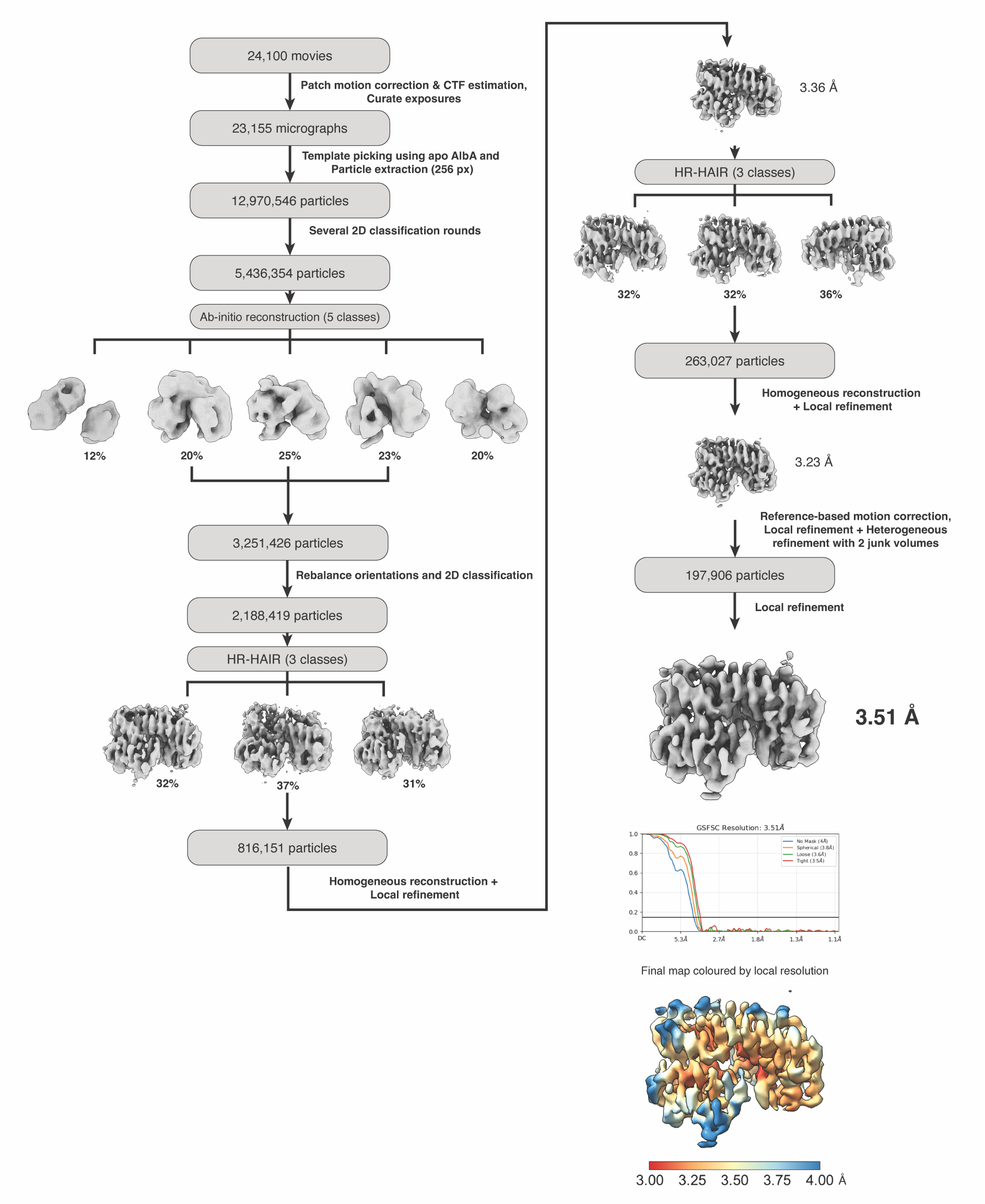
Cryo-EM data-processing workflow for AlbA:KMR-14-14. 24,100 movies were subjected to patch motion correction and patch CTF estimation, and exposures were curated to yield 23,155 micrographs. Particles were picked using templates from the AlbA dataset, extracted (12,970,546 particles) and reduced to 5,436,354 particles by several rounds of 2D classification. *Ab initio* reconstruction into five classes, shown as density maps, yielded three classes with interpretable protein features, which were combined to give 3,251,426 particles. Orientation rebalancing and 2D classification gave 2,188,419 particles, which were subjected to high-resolution heterogeneous ab-initio reconstruction (HR-HAIR) into 3 classes, and the best class of 816,151 particles was taken forward to homogeneous reconstruction and local refinement at 3.36 Å. A second round of HR-HAIR into 3 classes, followed by homogeneous reconstruction and local refinement of the best class of 263,027 particles, gave a map at 3.23 Å. Reference-based motion correction and a final local refinement produced the final reconstruction at 3.51 Å. The lower right panels show gold-standard FSC curves for the final refinement, calculated with no mask and with spherical, loose and tight masks, where the horizontal line marks the 0.143 threshold. The final map is also shown colored by local resolution over the range 3.00-4.00 Å.

**Suppl. Fig. 13.**
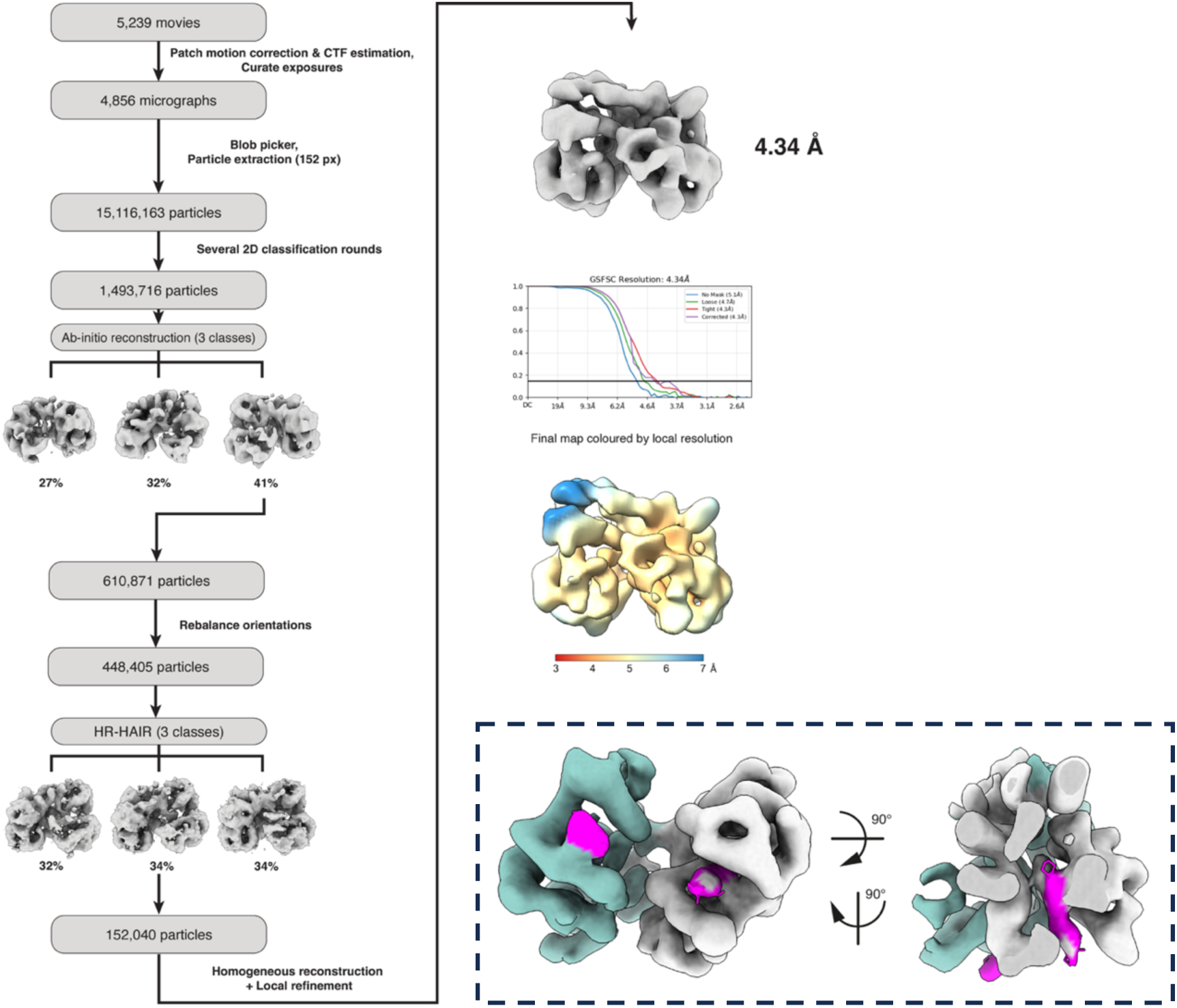
Cryo-EM data-processing workflow for AlbA:KMR-28-27. 5,239 movies were subjected to patch motion correction and patch CTF estimation, and exposures were curated to yield 4,856 micrographs. Particles were blob picked, extracted (15,116,163 particles) and reduced to 1,493,716 particles by several rounds of 2D classification. *Ab-initio* reconstruction into 3 classes, shown as density maps, yielded one class with interpretable protein features, which were combined to give 610,871 particles. Orientation rebalancing gave 448,405 particles, which were subjected to high-resolution heterogeneous ab-initio reconstruction (HR-HAIR) into 3 classes, and the best class of 152,040 particles was taken forward to homogeneous reconstruction and local refinement at 4.34 Å. The lower right panels show gold-standard FSC curves for the final refinement, calculated with no mask and with spherical, loose and tight masks, where the horizontal line marks the 0.143 threshold. The final map is also shown colored by local resolution over the range 3.00–7.00 Å. (**inset**) cryo-EM map of the AlbA:KMR-28-27 complex in two orthogonal views, colored as in Fig. 3f, g, with density for KMR-28-27 in magenta. Ligand density is present at the CTD site of each protomer.

**Suppl. Fig. 14.**
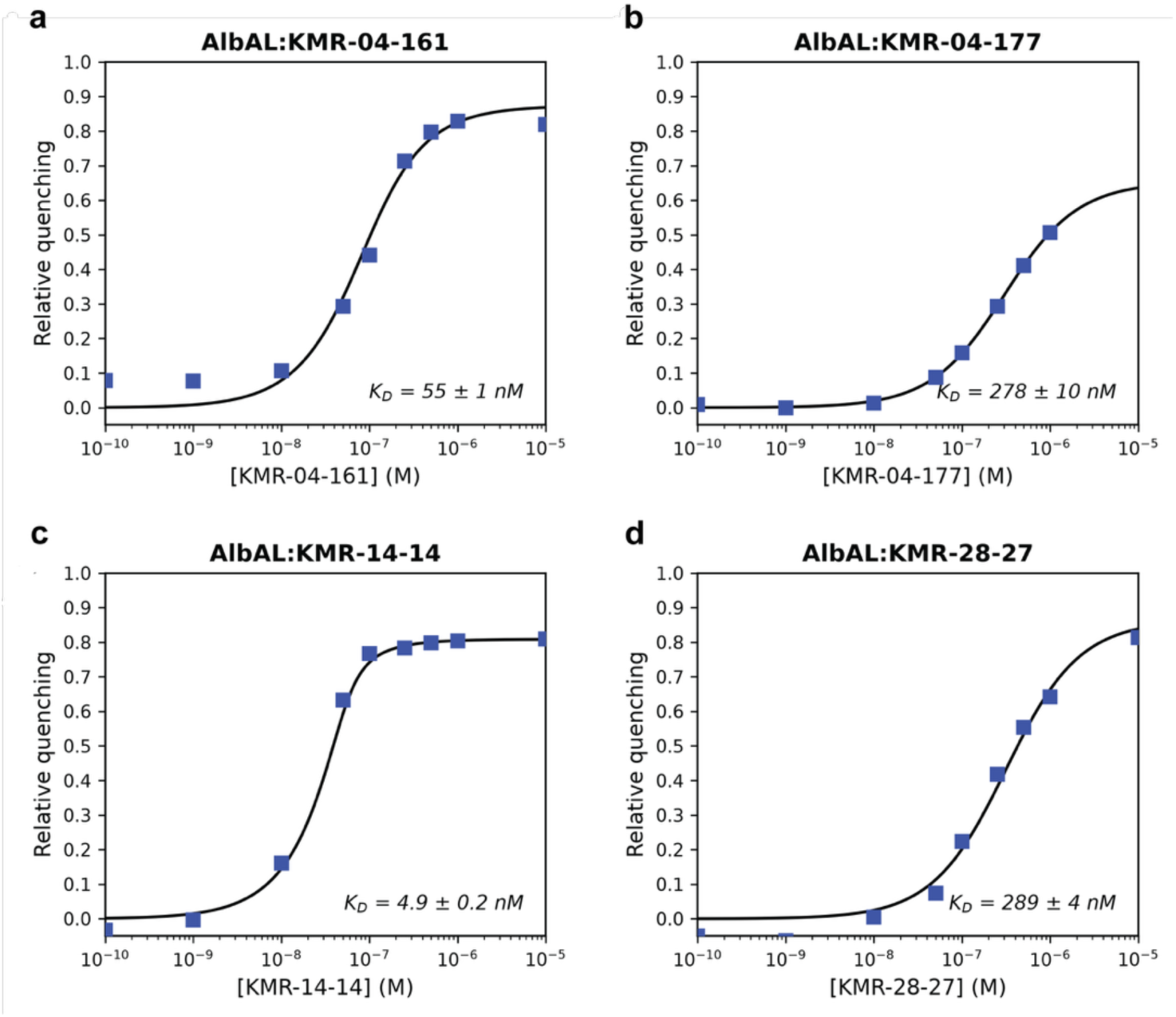
Single-site equilibrium dissociation constants for a subset of C8-linked PBD conjugates binding to full-length AlbA. (**a**–**d**) Binding curves for each indicated AlbA:PBD complex, derived from the concentration-dependent quenching of AlbA intrinsic fluorescence upon titration with the respective ligand. Axes, symbols, error bars and the high-concentration exclusion are as in **Suppl.** Fig. 4, with the emission wavelength determined independently for each titration series. Because only the CTD site is accessible in the AlbA dimer, the Morrison fit returns a single-site *K*_D_ rather than the apparent constant reported for AlbAS (**Suppl. Table 4**).

**Suppl. Fig. 15.**
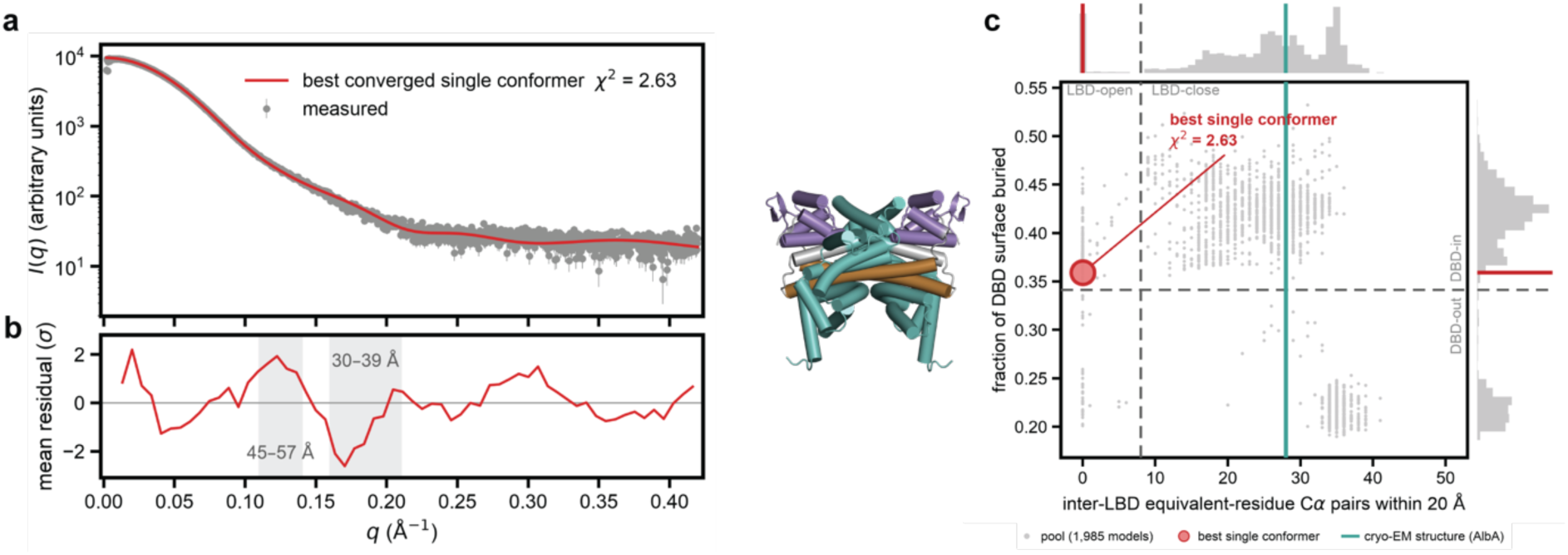
No single predicted conformer accounts for the measured AlbA scattering. (**a**) SEC-SAXS intensity (gray, every point with its uncertainty) over *q* = 0.0020-0.4198 Å, with the calculated curve of the best-fitting single conformer of the pool (red, χ^2^ = 2.63). This is the best-fitting structure for which every refined parameter converged inside its allowed range. Theoretical curves were computed with CRYSOL allowing the hydration-shell contrast and displaced volume of each structure to refine individually. For this conformer the contrast refined to 0.036 e Å^−3^ and the volume to 106,804 Å^3^, 7.2% above its unadjusted value. Two other conformers reach a nominally lower χ^2^, 2.36 and 2.46, but both sit exactly on the 7.5% volume ceiling the program imposes, so their optima lie on a parameter boundary rather than at a minimum. (**b**) Mean residual in units of the experimental uncertainty, averaged over 60 equal-width *q* bands of about 25 points each. The residual oscillates rather than scattering randomly. Shaded bands mark the two regions of largest misfit, *q* = 0.11-0.14 Å^−1^ where the band χ^2^ is 3.8σ and *q* = 0.16-0.21 Å^−1^ where it is 4.0σ. These correspond to real-space distances of 45-57 Å and 30-39 Å respectively, both at the scale of the separation between domains within the dimer. The conformer is also more compact than the particle in solution, with *R*_g_ = 31.3 Å against the measured 31.8 and *D*_max_ = 102 Å against 108. (**c**) The same conformer on the conformational map, with its cartoon representation. The axes are inter-LBD equivalent-residue Cα pairs within 20 Å and the fraction of DBD surface buried in the dimer. The pool of 1,985 models is in gray, with histograms of its distribution on each axis. Dashed lines divide the four conformational classes, which are named. The teal line marks the inter-LBD contact count of the experimental cryo-EM structure. In the cartoon the domains are colored as elsewhere, with the DBD purple, the CC brown, the LBD teal and the connecting segments gray.

**Suppl. Fig. 16.**
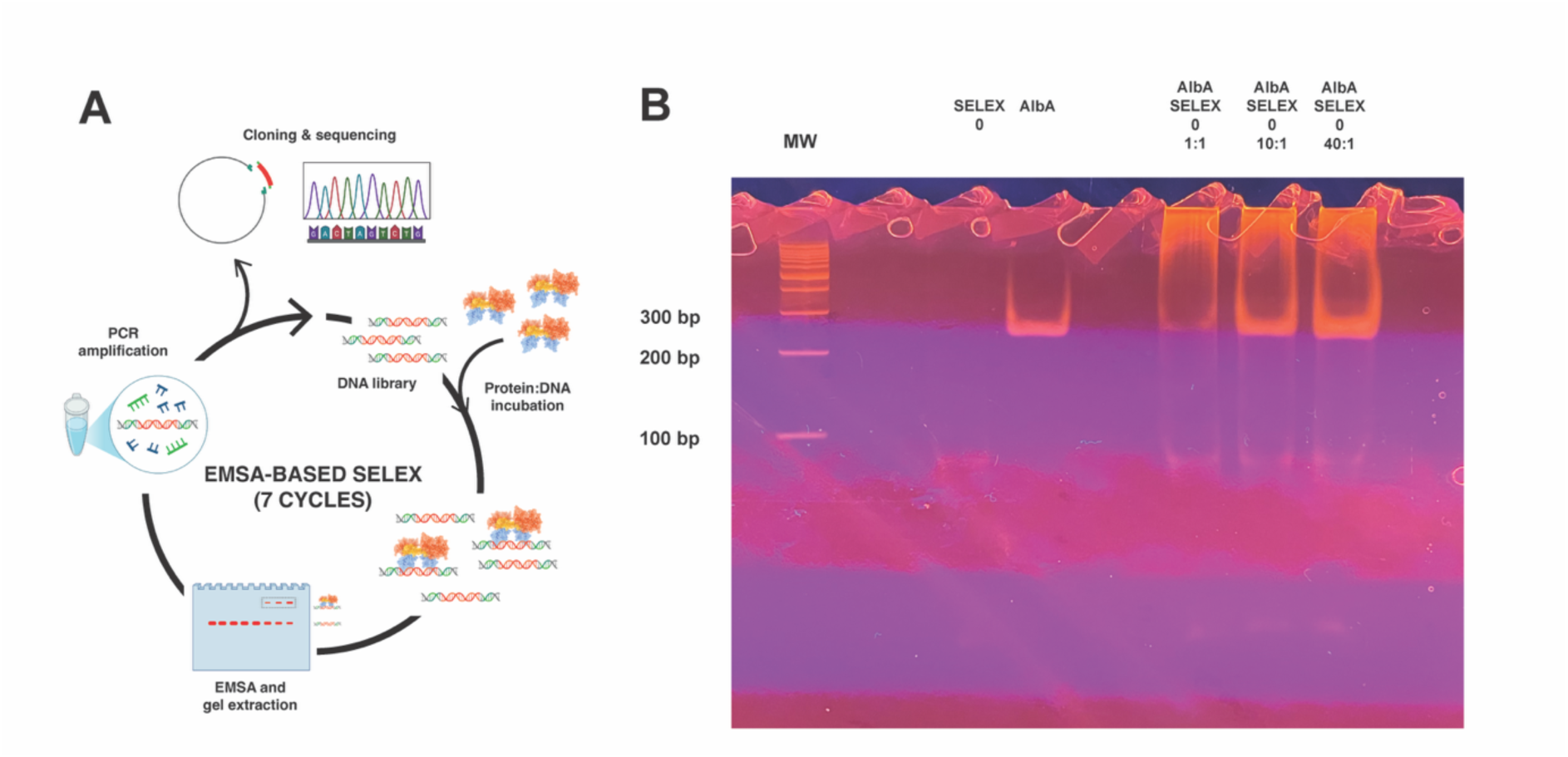
EMSA-based SELEX identifies AlbA-bound DNA sequences. (**a**) Schematic of the iterative EMSA-based SELEX workflow. A random-sequence dsDNA library is incubated with AlbA, protein-bound complexes are resolved and excised from a native polyacrylamide gel (EMSA), and the recovered DNA is PCR-amplified to seed the next of seven selection cycles; the final enriched pool is cloned and sequenced. (**b**) Representative EMSA showing increasing retention of the selected (round 4) DNA library in the wells with increasing AlbA:DNA molar ratio (1:1, 10:1, 40:1). MW, DNA molecular-weight ladder.

**Suppl. Fig. 17.**
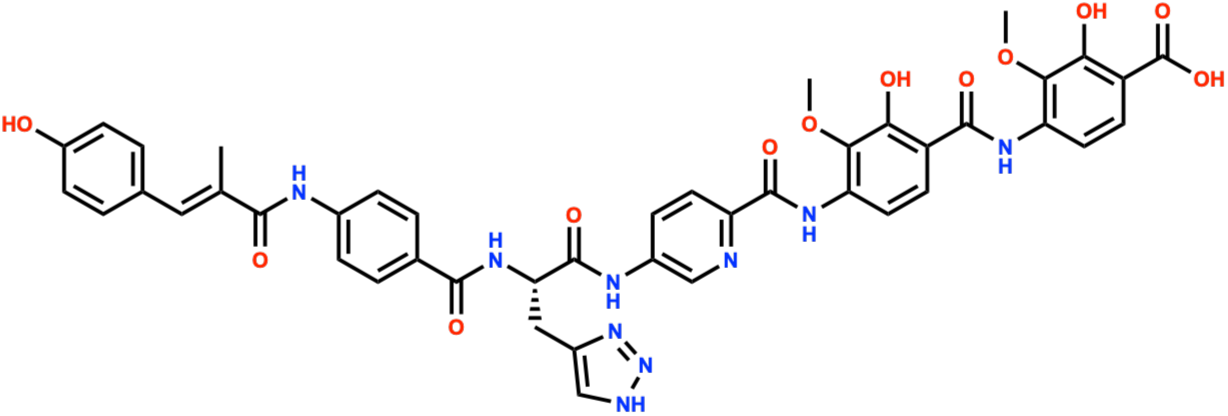
Chemical structure of the albicidin-derivative AF-CD-007.

**Suppl. Fig. 18.**
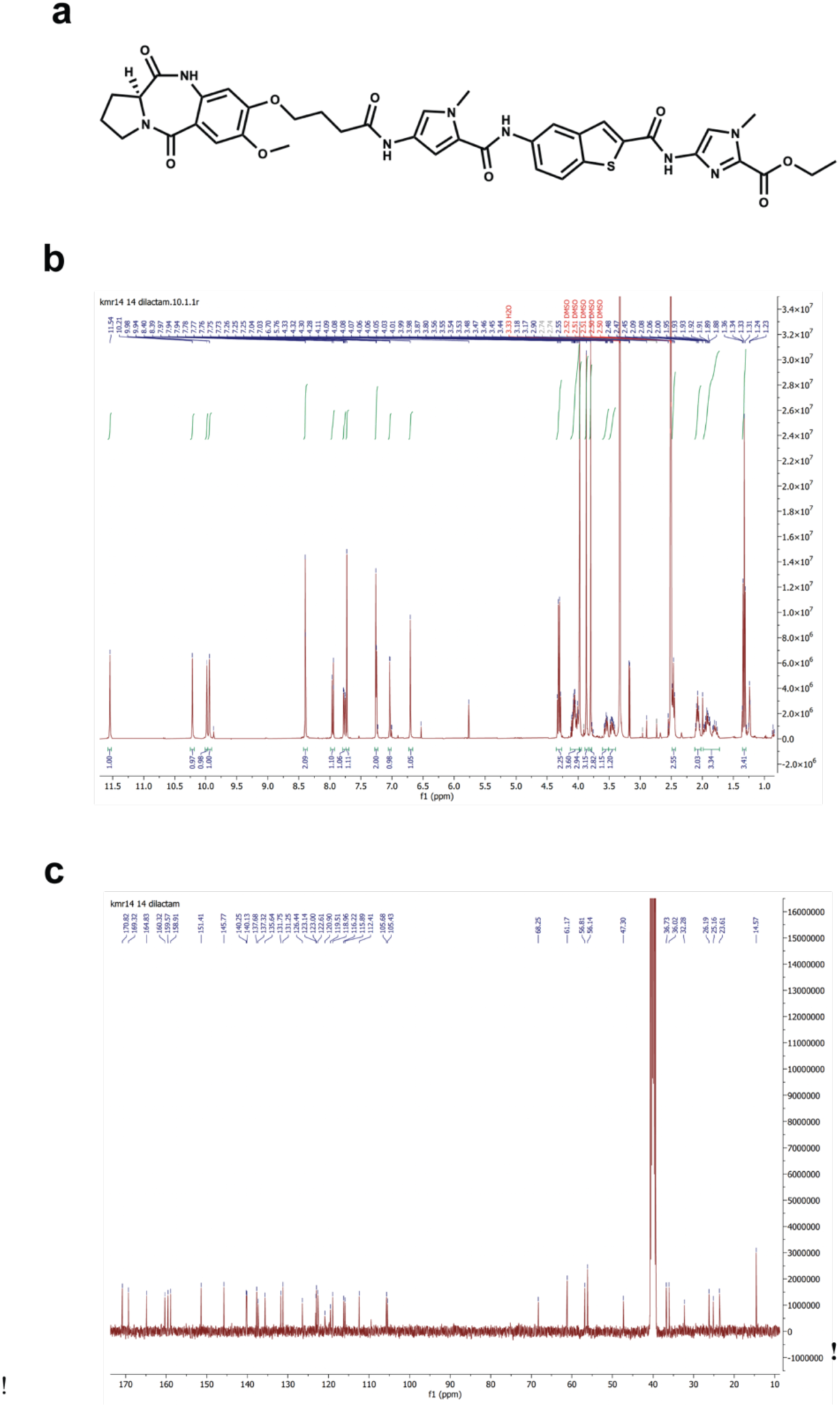
PA-267-36, the KMR-14-14 dilactam. **(a)** Chemical structure of PA-267-36. The synthetic procedure for this compound is reported in the extended Methods section at the end of this document. (**b**) ¹H NMR spectrum recorded at 400 MHz in DMSO-*d*₆. Numbers above the trace give picked chemical shifts in ppm and numbers below it give relative integrals. **(c)** Proton-decoupled ¹³C NMR spectrum recorded at 101 MHz in DMSO-*d*₆, with picked chemical shifts in ppm. Full assignments, coupling constants and high-resolution mass data are given in the Supplementary Methods.

**Suppl. Fig. 19.**
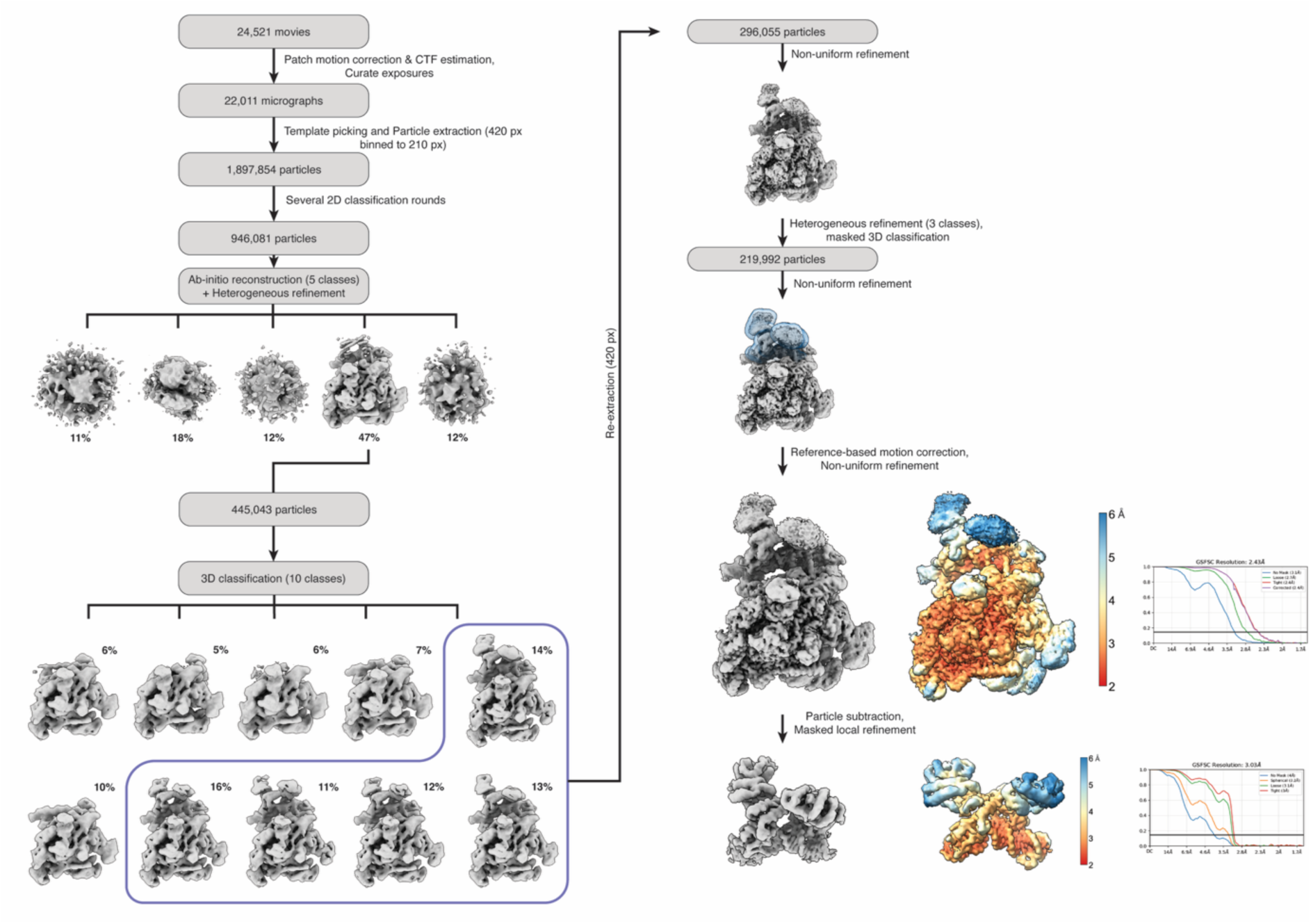
Cryo-EM data-processing workflow for the RNAP:*palbA*:AlbA(KMR-28-27) complex. 24,521 movies were subjected to patch motion correction and CTF estimation, and exposures were curated to give 22,011 micrographs. After manual picking of a subset of particles, template-based picking and particle extraction (420-pixel box, binned to 210 pixels) yielded 1,897,854 particles, reduced to 946,081 particles after several rounds of 2D classification. Ab-initio reconstruction into 5 classes and heterogeneous refinement identified an interpretable class (445,043 particles), which was subjected to 3D classification into 10 classes; the five classes containing AlbA density (outlined) were re-extracted at full (420-pixel) box size to give 296,055 particles. These were carried through non-uniform refinement, heterogeneous refinement, masked 3D classification (219,992 particles), a further round of non-uniform refinement, and reference-based motion correction followed by non-uniform refinement, yielding the consensus reconstruction at 2.43 Å (gold-standard FSC, 0.143 criterion). Particle subtraction and masked local refinement focused on the AlbA-containing region then produced the focused reconstruction at 3.03 Å. Local-resolution-colored maps (2–6 Å) and gold-standard FSC curves are shown for both reconstructions.

**Suppl. Fig. 20.**
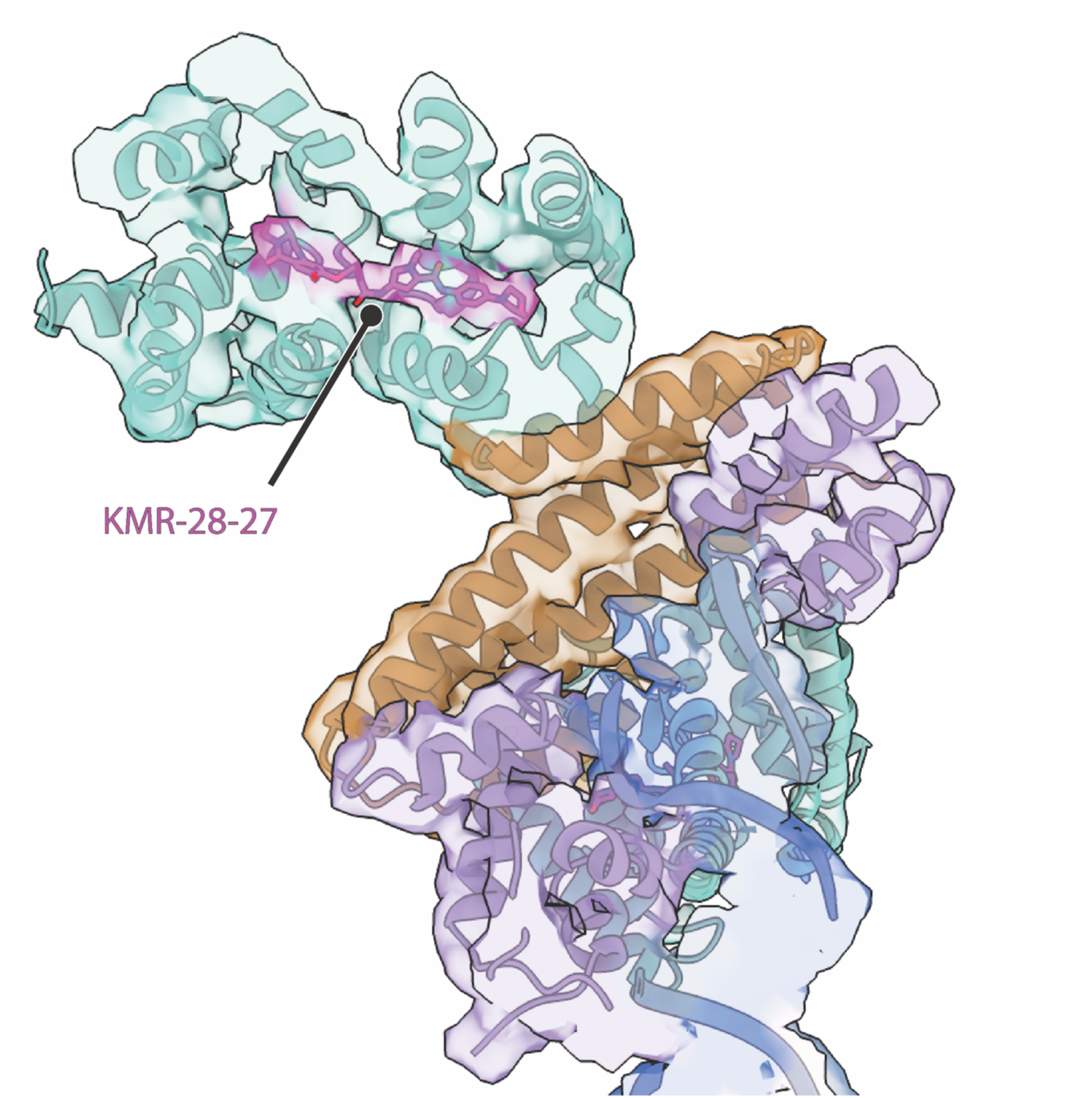
KMR-28-27 occupies the same site in the promoter-bound complex as in the crystallographic AlbAS structure. The focused-refinement reconstruction (Suppl. Fig. 17) fitted with the AlbA dimer model (LBD, teal; CC, brown; DBD, violet) bound to promoter DNA (blue), shown as ribbons within the semi-transparent map. Density for KMR-28-27 (magenta sticks) is resolved within the LBD tunnel at the same position identified in the crystallographic AlbAS:KMR-28-27 complex (Fig. 2).

**Suppl. Fig. 21.**
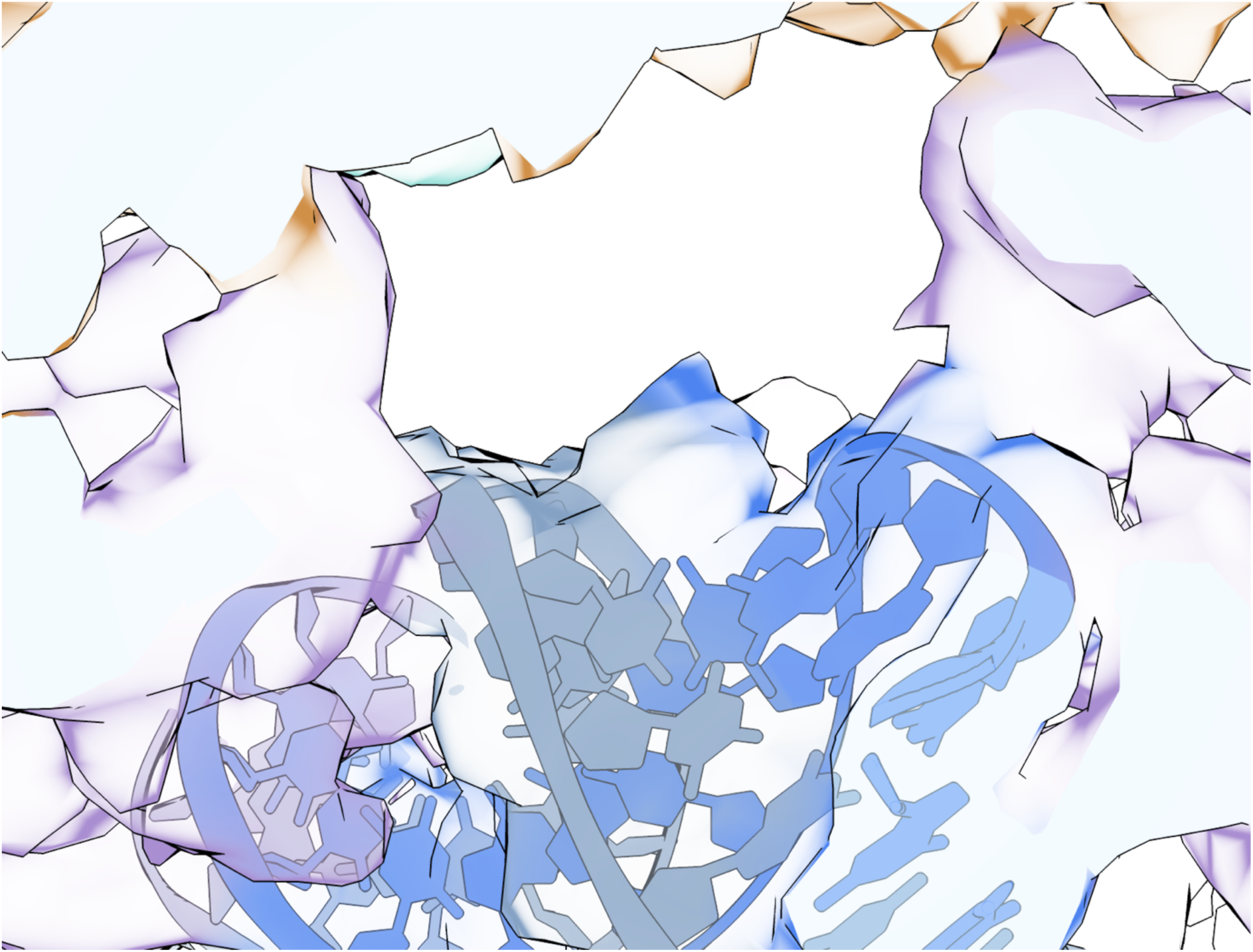
Unassigned density in the DNA minor groove between the two AlbA DBDs. Close-up of the promoter DNA (blue) engaged by the two AlbA DNA-binding domains (DBD, violet) in the promoter-bound reconstruction. An additional patch of density not accounted for by the AlbA–DNA model is present in the minor groove between the two DBDs (red circle). This might correspond to a low occupancy DNA-bound PBD molecule, consistent with the slow kinetics of covalent adduct formation.

**Suppl. Fig. 22.**
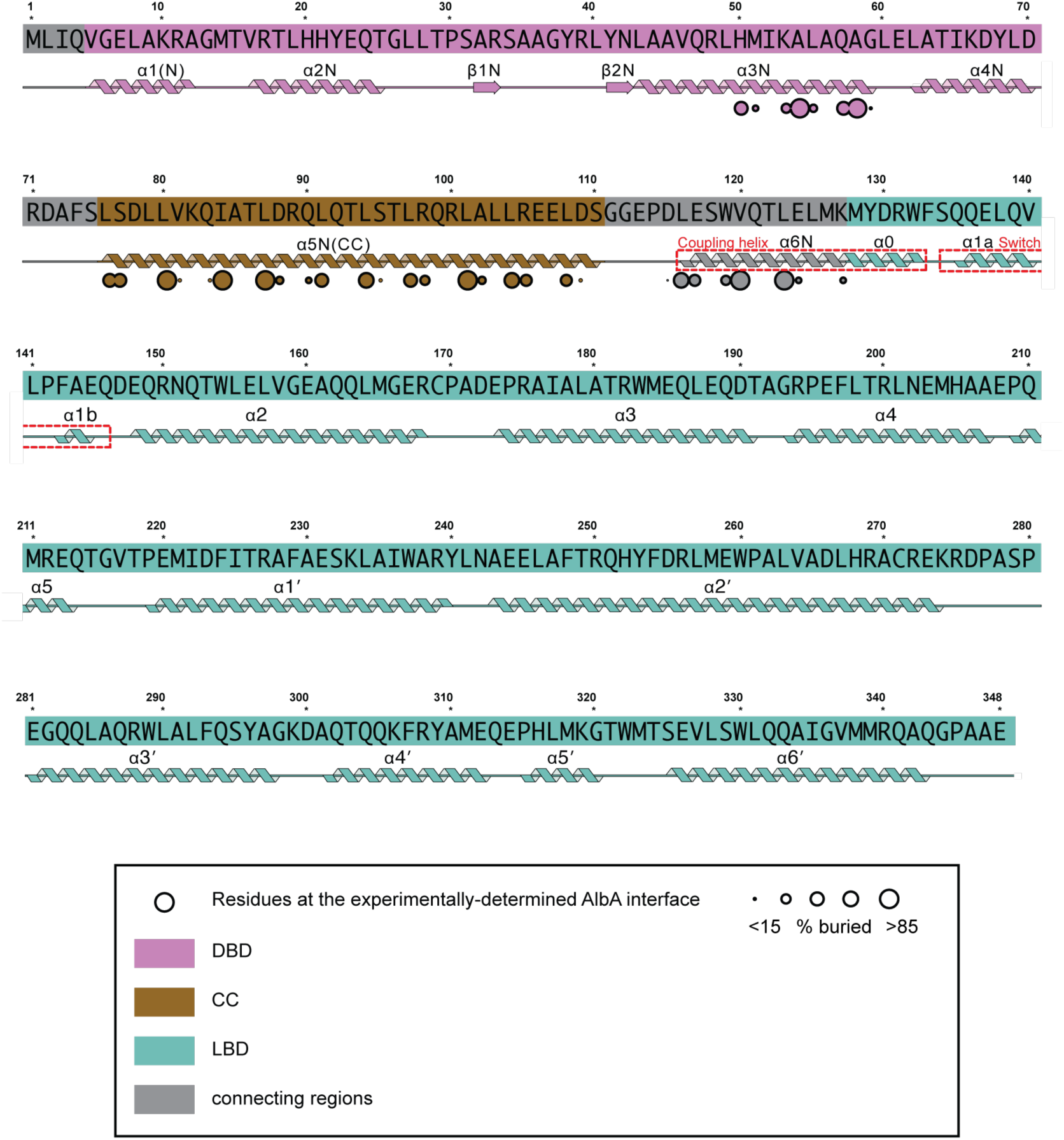
*K. oxytoca* AlbA secondary structure and dimerization contact map in the RNAP:*palbA*:AlbA complex. The AlbA sequence is shown in blocks numbered as in the full-length protein. The different domains are color-coded as indicated in the legend. Secondary structure determined for the complex is drawn beneath each block, with helices as coils and the two short β-bridges of the DBD wing (β1N and β2N) as arrows. Red dashed boxes mark the ‘coupling helix’, which runs continuously from α6N into α0, and the ‘switch’ region, comprising α1a and α1b. Residues involved in the observed dimer interface are highlighted by circles whose radius is proportional to the percentage of buried surface. Burial is confined to α3N of the DBD, to α5N(CC) along its whole length and to the connecting α6N region that follows it. Unlike isolated AlbA, no LBD residues are buried in this complex.

**Suppl. Table 1.** Apparent equilibrium dissociation constants for C8-linked PBD conjugates binding to AlbAS, determined by intrinsic tryptophan fluorescence quenching. Peak λ is the emission wavelength at which quenching was quantified, determined independently for each titration series from the mean spectrum of the lowest-concentration points. *K*_D,app_ is the apparent (macroscopic) dissociation constant obtained by fitting the Morrison quadratic tight-binding equation with the total binding-site concentration fixed at *P*_t_ = 50 nM; it describes overall ligand binding to AlbAS and is not resolved between the two crystallographically observed sites. SE is the asymptotic standard error taken from the square root of the corresponding diagonal element of the covariance matrix, and the 95% CI is the half-width of the interval computed as SE × *t*_0.975,ν_ with ν = (number of points fit − 3); the number of points retained per fit varies with the high-concentration exclusion rule described in the methods. *R*² is the coefficient of determination. Ligands are ordered by increasing *K*_D,app_. Values at or below *P*_t_ (KMR-14-14 through KMR-04-161) lie in the tight-binding regime and are to be read as upper bounds rather than precise constants; for KMR-28-32 and KMR-04-186 the asymptotic interval extends below zero and is therefore unreliable, and these two values should be treated as one-sided. *R*^2^ is the coefficient of determination.

| Ligand | Peak $\lambda$ (nm) | $K_{D,app}$ (nM) | SE (nM) | 95% CI (nM) | $R^2$ |
| --- | --- | --- | --- | --- | --- |
| KMR-14-14 | 334 | 0.54 | 0.13 | 0.34 | 0.99 |
| KMR-28-32 | 338 | 1.07 | 0.49 | 1.27 | 1.00 |
| KMR-28-27 | 340 | 6.91 | 1.20 | 2.93 | 0.99 |
| KMR-28-31 | 334 | 9.15 | 1.53 | 3.73 | 0.98 |
| KMR-04-184 | 340 | 15.2 | 1.62 | 3.96 | 1.00 |
| KMR-04-163 | 338 | 15.3 | 0.98 | 2.52 | 0.98 |
| KMR-04-161 | 338 | 18.3 | 2.23 | 5.74 | 0.90 |
| KMR-04-165 | 336 | 51.1 | 4.22 | 10.3 | 0.98 |
| KMR-28-38 | 338 | 54.0 | 4.52 | 11.1 | 0.99 |
| KMR-04-185 | 338 | 59.2 | 5.58 | 13.2 | 1.00 |
| KMR-04-177 | 340 | 160 | 8.19 | 20.0 | 0.99 |
| KMR-04-186 | 338 | 182 | 80.9 | 198 | 0.99 |
| KMR-04-154 | 340 | 189 | 30.5 | 78.5 | 1.00 |
| KMR-04-160 | 340 | 263 | 38.1 | 93.3 | 0.97 |

**Suppl. Table 2.**
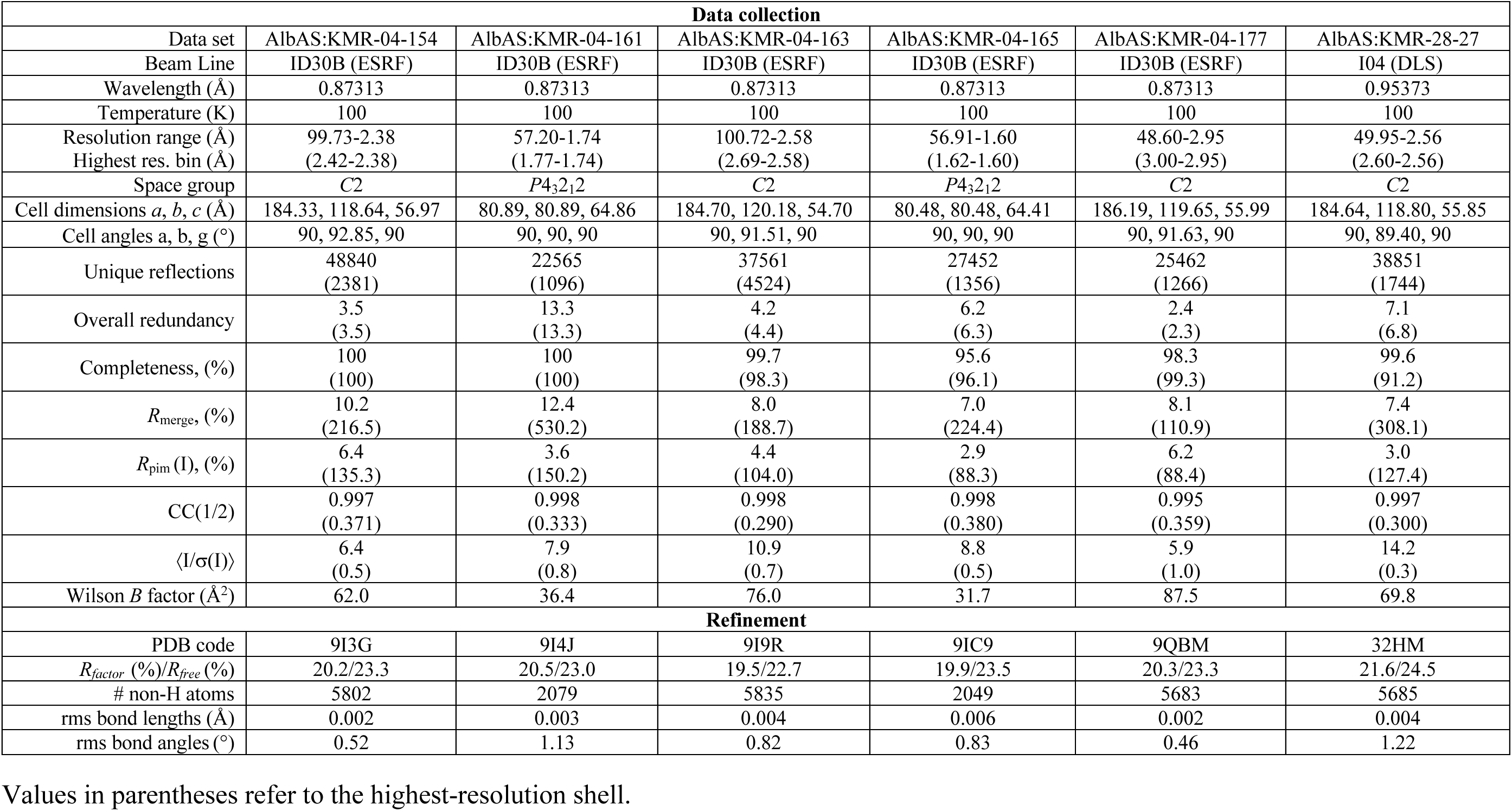
Data collection and refinement statistics.

**Suppl. Table 3.** Cryo-EM.

| Data set | AlbA<br>(EMD-57289)<br>(PDB 29QP) | AlbA:KMR-14-14<br>(EMD-58346)<br>(PDB 31ER) | AlbA:KMR-28-27<br>(EMD-59703)<br>(PDB 33ZZ) |
| --- | --- | --- | --- |
| **Data collection and processing** |  |  |  |
| Facility / Microscope | LonCEM / Krios | LonCEM / Krios | UniPD / Glacios |
| Magnification (kX) | 165 | 165 | 120 |
| Voltage (kV) | 300 | 300 | 200 |
| Electron exposure (e/Å<sup>2</sup>) | 70 | 70 | 60 |
| Defocus range (µm) | -0.6, -1,8 | -0.6, -1,8 | -0.8, -2.0 |
| Pixel size (Å) | 0.52 | 0.52 | 1.22 |
| Symmetry imposed | C1 | C1 | C1 |
| Initial particle images | 1360446 | 2188419 | 1498941 |
| Final particle images | 174223 | 197906 | 152040 |
| Map resolution (Å)<br>FSC threshold 0.143 | 3.62 | 3.51 | 4.34 |
| Map resolution range (Å) | 2.7-4.0 | 2.5-4.2 | 3.0-5.5 |
| **Refinement** |  |  |  |
| Initial model (PDB code) | 8RKY | 8RKY | 8RKY |
| Final model composition |  |  |  |
| Non-hydrogen atoms | 3342 | 3454 | 3432 |
| Protein | 408 | 408 | 408 |
| Nucleotides | 0 | 0 | 0 |
| Ligands | 0 | 2 | 2 |
| *B* factors (Å<sup>2</sup>) |  |  |  |
| Protein | 176.40 | 99.56 | 358.34 |
| Nucleotides |  |  |  |
| Ligands |  | 98.20 | 325.30 |
| R.m.s. deviations |  |  |  |
| Bond lengths (Å) | 0.004 (0) | 0.004 (0) | 0.003 (0) |
| Bond angles (°) | 0.607 (0) | 0.808 (2) | 0.602 (0) |
| **Validation** |  |  |  |
| MolProbity score | 1.65 | 1.62 | 1.58 |
| Clashscore | 5.94 | 3.74 | 3.75 |
| Poor rotamers (%) | 2.65 | 2.65 | 2.65 |
| Ramachandran plot |  |  |  |
| Favored (%) | 98.02 | 97.28 | 97.52 |
| Allowed (%) | 1.98 | 2.72 | 2.48 |
| Disallowed (%) | 0.00 | 0.00 | 0.00 |

**Suppl. Table 3. Cryo-EM**
| Data set | RNAP-DNA-<br>-AlbA(KMR-28-27)<br>composite map<br>(EMD-57337)<br>(PDB 29RP) | RNAP-DNA-<br>-AlbA(KMR-28-27)<br>consensus map<br>(EMD-57335) | RNAP-DNA-<br>-AlbA(KMR-28-27)<br>focused map<br>(EMD-57336) |
| --- | --- | --- | --- |
| Facility / Microscope | eBIC / Krios | eBIC / Krios | eBIC / Krios |
| Magnification (kX) | 105 kX | 105 kX | 105 kX |
| Voltage (kV) | 300 kV | 300 kV | 300 kV |
| Electron exposure (e/Å<sup>2</sup>) | 40 | 40 | 40 |
| Defocus range (μm) | -0.6, -2.0 | -0.6, -2.0 | -0.6, -2.0 |
| Pixel size (Å) | 0.825 | 0.825 | 0.825 |
| Symmetry imposed | C1 | C1 | C1 |
| Initial particle images | 445043 | 445043 | 445043 |
| Final particle images | 219992 | 219992 | 219992 |
| Map resolution (Å)<br>FSC threshold 0.143 |  | 2.43 | 3.03 |
| Map resolution range (Å)* |  | 1.8-7.5 | 2.0-7.0 |
| Initial model (PDB code) | 6XL5 |  |  |
| Final model composition |  |  |  |
| Non-hydrogen atoms | 36228 |  |  |
| Protein | 4368 |  |  |
| Nucleotides | 73 |  |  |
| Ligands | 8 |  |  |
| *B* factors (Å<sup>2</sup>) |  |  |  |
| Protein | 121.77 |  |  |
| Nucleotides | 99.36 |  |  |
| Ligands | 180.63 |  |  |
| R.m.s. deviations |  |  |  |
| Bond lengths (Å) | 0.005 (5) |  |  |
| Bond angles (°) | 0.694 (12) |  |  |
| MolProbity score | 1.80 |  |  |
| Clashscore | 13.09 |  |  |
| Poor rotamers (%) | 0.46 |  |  |
| Ramachandran plot |  |  |  |
| Favored (%) | 96.98 |  |  |
| Allowed (%) | 2.99 |  |  |
| Disallowed (%) | 0.02 |  |  |

**Suppl. Table 4.** Equilibrium dissociation constants for a subset of C8-linked PBD conjugates binding to full-length AlbA, determined by intrinsic tryptophan fluorescence quenching, compared with the apparent constants for AlbAS.

| Ligand | AlbAS $K_{D,app}$ (nM) | AlbA $K_D$ (nM) | Fold weaker | $R^2$ |
| --- | --- | --- | --- | --- |
| KMR-14-14 | $0.54 \pm 0.13$ | $4.92 \pm 0.22$ | 9.1× | 1.00 |
| KMR-28-27 | $6.91 \pm 1.20$ | $289 \pm 4$ | 41.9× | 0.99 |
| KMR-04-161 | $18.3 \pm 2.2$ | $55.5 \pm 0.8$ | 3.0× | 0.98 |
| KMR-04-177 | $160 \pm 8$ | $278 \pm 10$ | 1.7× | 1.00 |

**Suppl. Table 5.** SAXS parameters for AlbA and the AlbA^F143A^ variant.

| Parameter | AlbA | AlbA<sup>F143A</sup> | ATSAS program |
| --- | --- | --- | --- |
| **Guinier analysis (reciprocal space)** |  |  |  |
| $R_g$ (Å) | $31.80 \pm 0.06$ | $39.02 \pm 0.86$ | AUTORG |
| $I(0)$ | $9.5e+03 \pm 4.8$ | $60.638 \pm 0.205$ | AUTORG |
| Guinier points used | 29–139 (111) | 13–45 (33) | AUTORG |
| Aggregation flag | — | 0.080 | AUTORG |
| **Distance distribution $P(r)$ (real space)** |  |  |  |
| Angular range (Å<sup>-1</sup>) | 0.0020–0.2515 | 0.0089–0.2045 | DATGNOM |
| Real space $R_g$ (Å) | $31.83 \pm 0.025$ | $39.90 \pm 0.099$ | DATGNOM |
| $D_{max}$ (Å) | 108.08 | 125.88 | DATGNOM |
| **Volume and shape** |  |  |  |
| Porod volume (Å<sup>3</sup>) | 126556 | 126550 | DATPOROD /<br>DATCLASS |
| Shape class | compact | flat | DATPOROD /<br>DATCLASS |
| **Molecular weight (Da)** |  |  |  |
| MW, Porod | 77,158 | 78,578 | DATMW |
| MW, $Q_p$ | 85,370 | 80,750 | DATMW |
| MW, $V_c$ | 79,189 | 76,904 | DATMW |
| MW, shape-aware | 87,602 | 99,781 | DATMW |
| Median (s.e. median) | $82,279 \pm 6,210$ | $79,664 \pm 13,330$ | |

**Suppl. Table 6.**
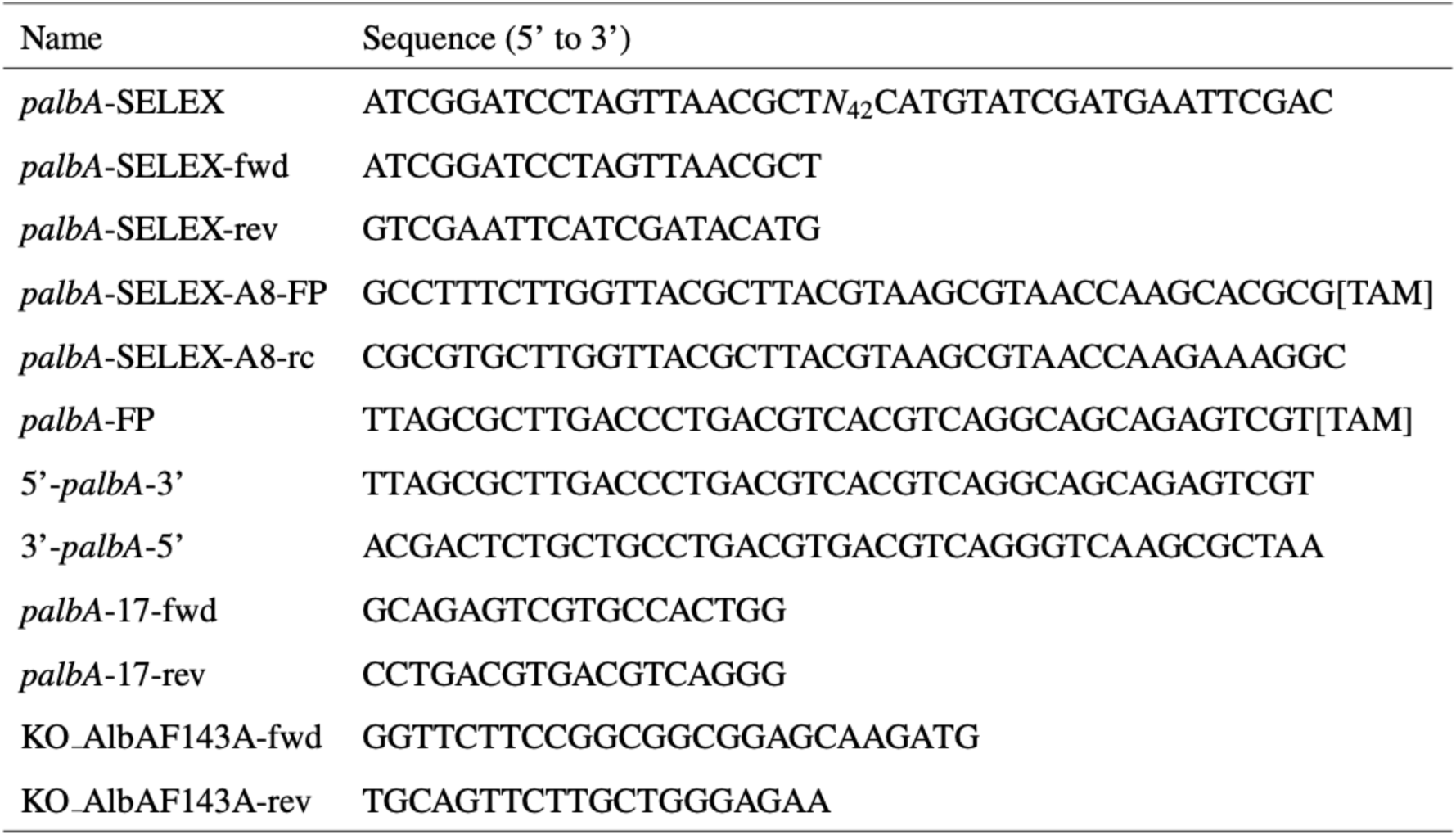
Primers and DNA oligonucleotides used in this study.

## Extended Methods

### Chemical synthesis

Synthetic procedures for KMR-04-154, KMR-04-160, KMR-04-161, KMR-04-163, KMR-04-165, KMR-04-177, KMR-04-184, KMR-04-186, KMR-14-14, KMR-28-27, KMR-28-31, KMR-28-32 and KMR-28-38 were described previously^1,2^. The of AF-CD-007 was also previously reported^3^. The Synthetic procedure for PA-267-36 (KMR-14-14 dilactam) is as follows:

Methyl 4-(4-formyl-2-methoxyphenoxy)butanoate (**1**)

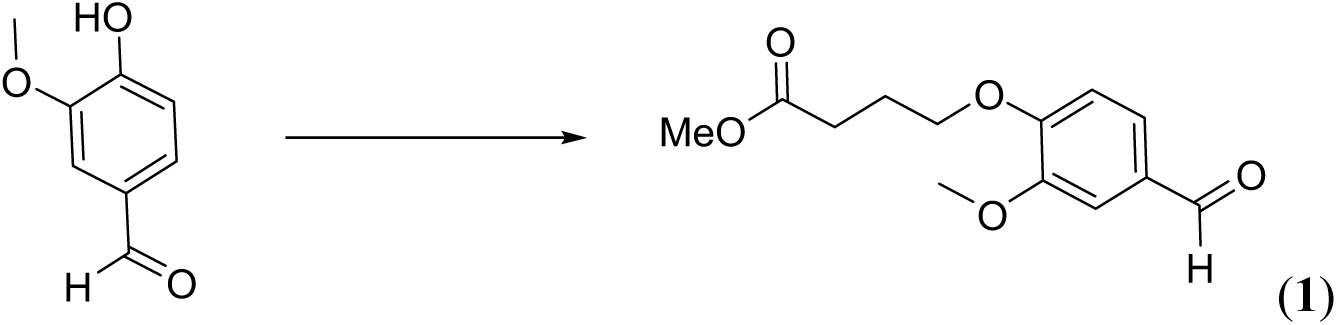

A mixture of vanillin (20.0 g, 131 mmol), methyl 4-bromobutanoate (17.5 mL, 139 mmol) and potassium carbonate (27.2 g, 197 mmol) in *N*,*N*-dimethylformamide (100 mL) was stirred at room temperature for 18 h. The reaction mixture was diluted with water (500 mL) and the title compound (30.2 g, 91%) was obtained by filtration as a white solid. The product was carried through to the next step without any further purification.

^1^H NMR (400 MHz, CDCl_3_) *δ* 9.84 (s, ^1^H), 7.46-7.37 (m, 2H), 6.98 (d, *J*=8.2 Hz, 1H), 4.16 (t, *J*=6.3 Hz, 2H), 3.91 (s, 3H), 3.69 (s, 3H), 2.56 (t, *J*=7.2 Hz, 2H), 2.20 (quin, *J*=6.7 Hz, 2H); ^13^C NMR (100 MHz, CDCl_3_) *δ* 190.9, 173.4, 153.8, 149.9, 130.1, 126.8, 111.6, 109.2, 67.8, 56.0, 51.7, 30.3, 24.2; [M+H]^+^ LCMS C_13_H_16_O_5_ calculated 253.11, found 253.1.

Methyl 4-(4-formyl-2-methoxy-5-nitrophenoxy)butanoate (**2**)

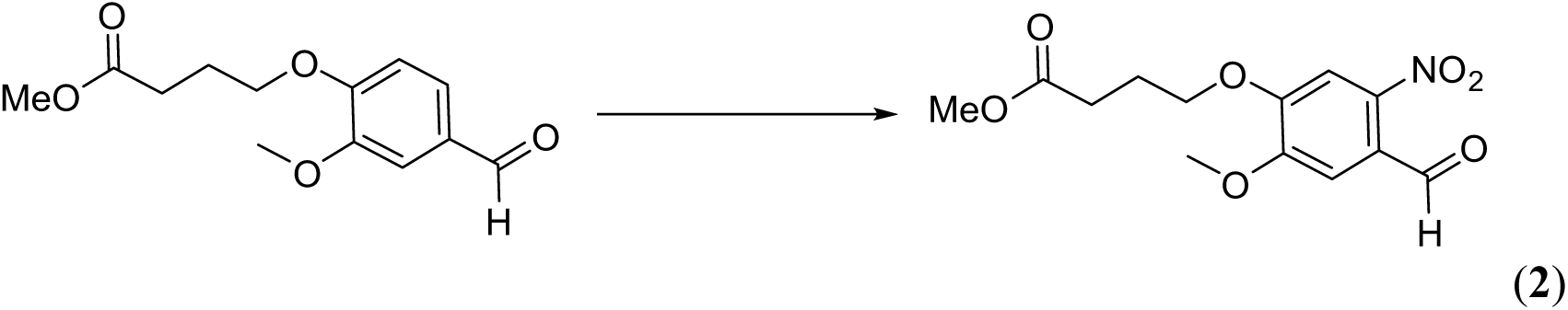

To a stirring solution of potassium nitrate (10.0 g, 98.9 mmol) in TFA (50 mL) at 0 °C was added dropwise a solution of methyl 4-(4-formyl-2-methoxyphenoxy)butanoate (**1**) (20.0 g, 79.2 mmol) in trifluoroacetic acid (50 mL). The reaction mixture was stirred at room temperature for 1 h. It was then concentrated *in vacuo* and diluted with ethyl acetate (400 mL). The organic layer was washed with brine (3 x 100 mL) and a saturated aqueous solution of sodium hydrogen carbonate (2 x 80 mL), dried over sodium sulfate, filtered and concentrated to give the title compound (23.5 g, 100%) as a yellow solid. The product was carried through to the next step without any further purification.

^1^H NMR (400 MHz, CDCl_3_) *δ* 10.42 (s, 1H), 7.60 (s, 1H), 7.39 (s, 1H), 4.21 (t, *J*=6.3 Hz, 2H), 3.98 (s, 3H), 3.70 (s, 3H), 2.61-2.53 (m, 2H), 2.22 (quin, *J*=6.6 Hz, 2H); ^13^C NMR (100 MHz, CDCl_3_) *δ* 187.8, 173.2, 153.5, 151.7, 143.8, 125.5, 109.9, 108.1, 68.6, 56.6, 51.8, 30.2, 24.1; [M+H]^+^ LCMS C_13_H_15_NO_7_ calculated 298.09, found 298.1.

5-Methoxy-4-(4-methoxy-4-oxobutoxy)-2-nitrobenzoic acid (**3**)

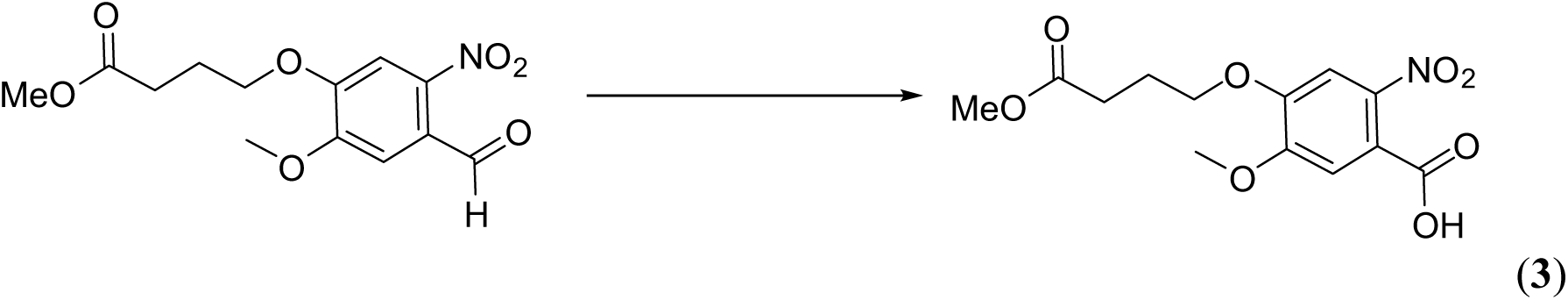

To a solution of methyl 4-(4-formyl-2-methoxy-5-nitrophenoxy)butanoate (**2**) (23.0 g, 77.4 mmol) in acetone (600 mL) was added a hot (70 °C) solution of potassium permanganate (46.0 g, 291 mmol) in water (400 mL). The reaction mixture was stirred at 70 °C for 3 h. The reaction mixture was cooled to room temperature and passed through celite. The cake of celite was washed with hot water (200 mL). A solution of sodium meta bisulfite in hydrochloric acid (1 M, 200 mL) was added to the filtrate which was extracted with dichloromethane (2 x 400 mL). The organic layer was dried over sodium sulfate, filtered and concentrated. The resulting residue was purified by column chromatography (silica), eluting with methanol-dichloromethane (from 0% to 50%), to give the title compound (17.0 g, 70%) as a pale yellow solid.

_1_H NMR (400 MHz, MeOD) *δ* 7.47 (s, 1H), 7.25 (s, 1H), 4.13 (t, *J*=6.2 Hz, 2H), 3.94 (s, 3H), 3.68 (s, 3H), 2.54 (t, *J*=7.2 Hz, 2H), 2.17-2.06 (m, 2H); ^13^C NMR (100 MHz, MeOD) *δ* 175.3, 168.6, 153.8, 151.3, 143.1, 122.8, 112.4, 109.2, 69.6, 57.0, 52.2, 31.2, 25.5; [M-H]^-^ LCMS C_13_H_15_NO_8_ calculated 312.07, found 312.1.

Methyl (5-methoxy-4-(4-methoxy-4-oxobutoxy)-2-nitrobenzoyl)-*L*-prolinate (**4**)

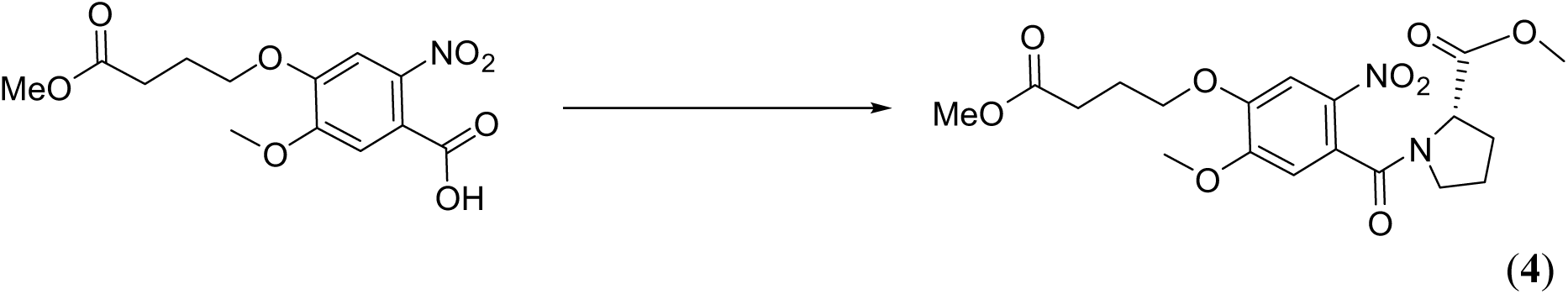

A mixture of 5-methoxy-4-(4-methoxy-4-oxobutoxy)-2-nitrobenzoic acid (**3**) (2.00 g, 6.37 mmol), oxalyl chloride (1.65 mL, 19.3 mmol) and anhydrous *N*,*N*-dimethylformamide (2 drops) in anhydrous dichloromethane (25 mL) was stirred at room temperature for 1 h.

Anhydrous toluene (5 mL) was added to the reaction mixture which was then concentrated *in vacuo*. The residue was dissolved in anhydrous dichloromethane (3 mL) was added dropwise to a solution of methyl *L-*prolinate (1.08 g, 8.4 mmol) and triethylamine (2.7 mL, 19.25 mmol) in anhydrous dichloromethane (25 mL) at -10 °C. The reaction mixture was stirred at room temperature for 2 h and then washed with hydrochloric acid (1 M, 15 mL) and a saturated aqueous solution of sodium chloride (15 mL), dried over sodium sulfate, filtered and concentrated. The resulting residue was purified by column chromatography (silica), eluting with methanol-dichloromethane (from 0% to 5%), to give the title compound (1.3 g, 48%) as a yellow oil. ^1^H NMR (400 MHz, CHLOROFORM-*d*) *δ*, 7.69 (s, 1H), 6.79 (s, 1H), 4.75-4.69 (m, 1H), 4.17-4.14 (t, *J* = 6.4, 2H) 4.13-4.00 (m, 1H), 3.93 (s, 3H), 3.66 (s, 3H), 3.53 (s, 3H), 3.33-3.29 (m, 1H), 3.21-3.17 (m, 1H), 2.58-2.52 (m, 3H), 2.38-2.30 (m, 1H), 2.24-2.18 (m, 3H), 2.12-1.89 (m, 4H). ^13^C NMR (101 MHz, CHLOROFORM-*d*) *δ*, 173.2, 172.6, 154.8, 148.5, 137.3, 1274, 110.2, 108.2, 68.1, 60.6, 58.5, 56.7, 52.4, 51.5, 48.3, 46.2, 31.0, 30.3, 29.6, 24.5, 24.0, 23.0; [M+H]^+^ LCMS C_19_H_24_NO_9_ calculated 425.15, found 425.1.

Methyl (*S*)-4-((7-methoxy-5,11-dioxo-2,3,5,10,11,11a-hexahydro-1*H*-benzo[*e*]pyrrolo[1,2-*a*][1,4]diazepin-8-yl)oxy)butanoate (**5**)

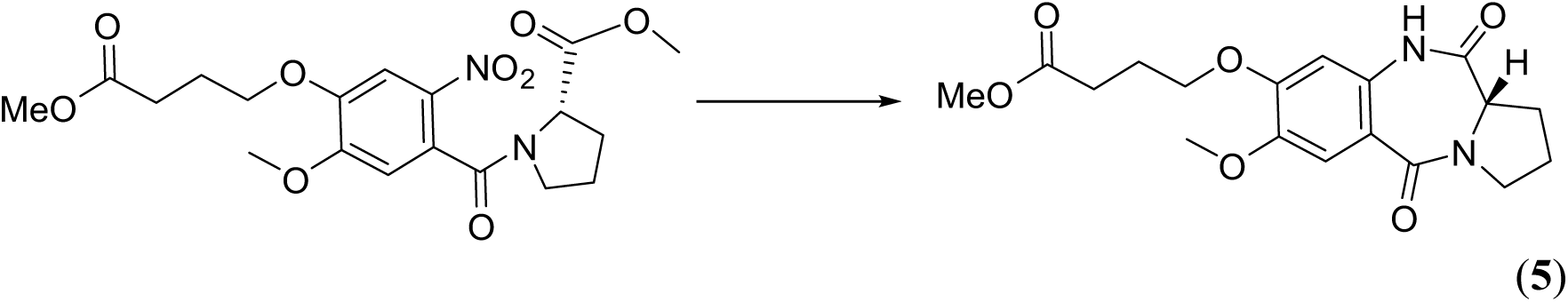

To a solution of methyl Methyl (5-methoxy-4-(4-methoxy-4-oxobutoxy)-2-nitrobenzoyl)-*L*-prolinate (**4**) (1.5 g, 2.36 mmol) in ethanol (20 mL) and ethyl acetate (50 mL) was added palladium on activated charcoal (10% wt.) (100 mg). The reaction mixture was hydrogenated at 35 psi for 16 h in a Parr apparatus. The reaction mixture was filtered through celite and the resulting cake was washed with ethyl acetate. The filtrate was concentrated *in vacuo* to give the title compound (775 g, 91%) as an orange oil. The product was carried through to the next step without any further purification^1^H NMR (400 MHz, CHLOROFORM-*d*) *δ* 8.91 (s, 1H), 7.42 (s, 1H), 6.52 (s, 1H), 4.04 (td, *J* = 3.08, 5.92 Hz, 3H), 3.87 (s, 3H), 3.71 - 3.78 (m, 1H), 3.67 (s, 3H), 3.55 - 3.62 (m, 1H), 2.70 - 2.76 (m, 1H), 2.53 (t, *J* = 7.05 Hz, 2H), 2.15 – 2.09 (m, 2H), 1.96 - 2.02 (m, 3H). ^13^C NMR (101 MHz, CHLOROFORM-*d*) *δ* 173.5, 171.2, 165.3, 151.5, 146.6, 129.8, 119.3, 112.3, 105.1, 67.8, 56.9, 56.2, 51.7, 47.3, 30.2, 26.5, 24.1, 23.6. [M+H]^+^ LCMS C_18_H_22_NO_8_ calculated 363.15, found 363.1.

(*S*)-4-((7-methoxy-5,11-dioxo-2,3,5,10,11,11a-hexahydro-1*H*-benzo[*e*]pyrrolo[1,2-*a*][1,4]diazepin-8-yl)oxy)butanoic acid (**6**)

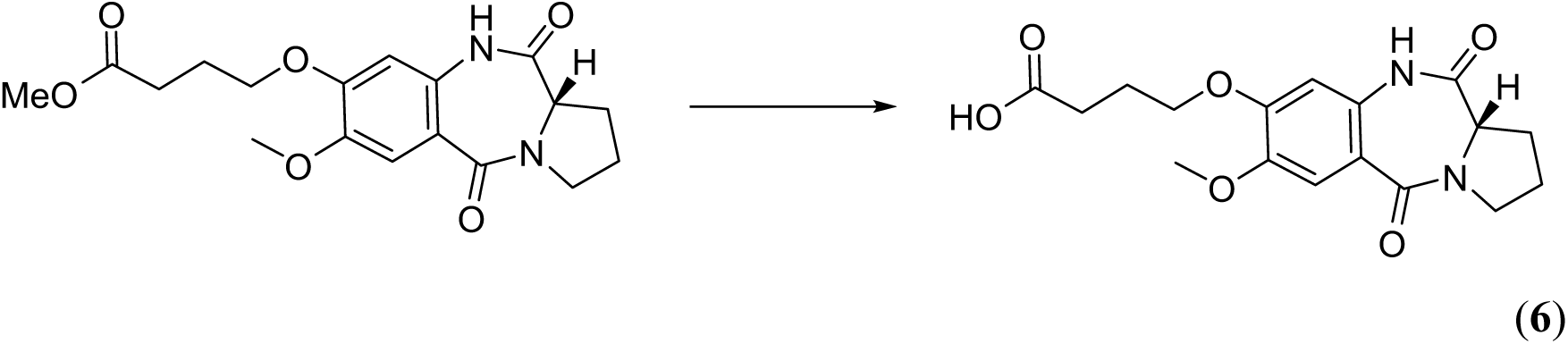

To a solution of allyl Methyl (*S*)-4-((7-methoxy-5,11-dioxo-2,3,5,10,11,11a-hexahydro-1*H*- benzo[*e*]pyrrolo[1,2-*a*][1,4]diazepin-8-yl)oxy)butanoate (**5**) (770 mg, 2.12 mmol) in 1,4-dioxane (10 mL) was added a 1 M aqueous solution of sodium hydroxide (10.0 mL). The reaction mixture was stirred at room temperature for 2 h and was then concentrated *in vacuo*, after which water (20 mL) was added and the aqueous layer was acidified with a 5 M aqueous solution of acetic acid (10 mL, 50 mmol). The aqueous layer was extracted with ethyl acetate (2 x 50 mL). The combined organic extracts were washed with a saturated aqueous solution of sodium chloride (50 mL), dried over sodium sulfate, filtered and concentrated to give the title compound (690 mg, 93%) as a white solid. The product was carried through to the next step without any further purification.

^1^H NMR (400 MHz, MeOH-*d*_4_) δ: 7.27 (s, 1H), 6.57 (s, 1H), 4.10-4.05 (m, 1H), 3.99 (t, *J* = 6.4Hz, 2H), 3.75 (s, 3H), 3.69-3.59 (m, 1H), 3.53-3.42 (m, 1H), 3.24-3.17 (m, 2H), 2.59-2.51 (m, 1H), 2.42 (t, *J* = 7.2Hz, 2H), 2.03-1.90 (m, 4H). ^13^C NMR (101 MHz, MeOH-*d*_4_): 176.8, 172.4, 167.6, 153.5, 147.9, 132.3, 120.0, 113.1, 106.5, 69.1, 58.4, 56.6, 31.2, 27.0, 25.5, 24.6. [M-H]^-^ LCMS C_17_H_20_N2O_6_ calculated 347.12, found 347.1.

Methyl 5-(4-((*tert*-butoxycarbonyl)amino)-1-methyl-1*H*-pyrrole-2-carboxamido)benzo[*b*]thiophene-2-carboxylate (**7**)

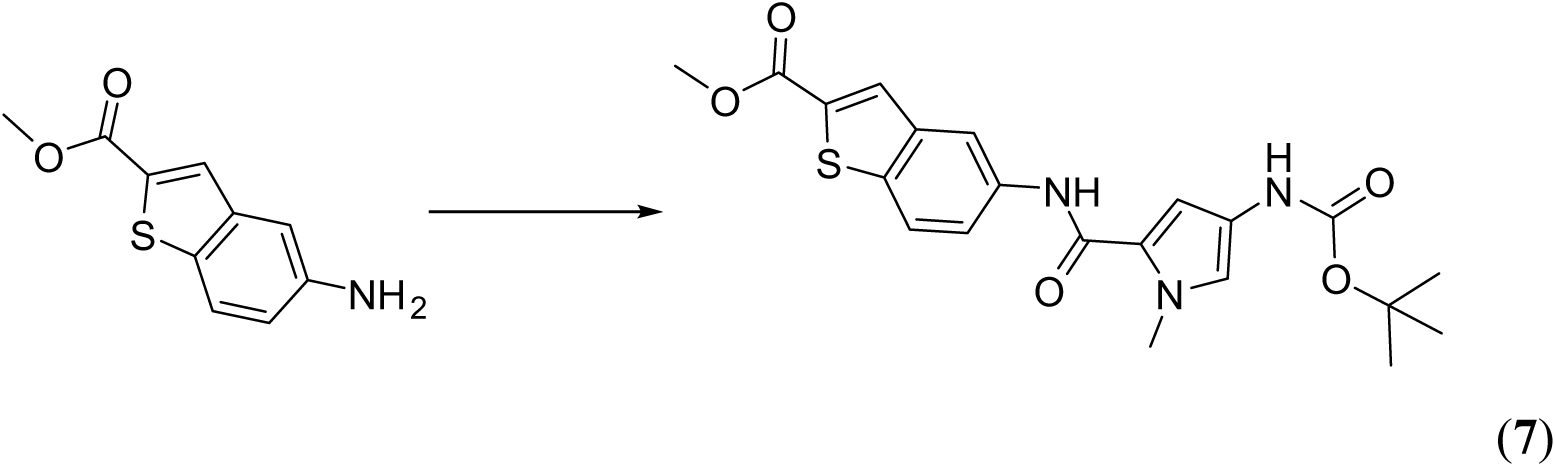

4-((*Tert*-butoxycarbonyl)amino)-1-methyl-1*H*-pyrrole-2-carboxylic acid (150 mg, 0.624 mmol, 1.0 eq.) was dissolved in DMF (2.5 mL) and activated with EDC (2.0 eq, 239.37 mg, 1.25 mmol) and DMAP (2.5 eq, 190.69 mg, 1.56 mmol) for 30 min with stirring. Amine 1.2 (1.5 eq, 194.08 mg, 0.936 mmol) was added, and the mixture stirred overnight. The reaction was quenched with ice-cold water (50mL) and extracted with ethyl acetate (3x 50 mL). The combined organic fractions were washed sequentially with 1 M citric acid, saturated sodium bisulfate, water, and brine, dried over magnesium sulfate, filtered, and solvent evaporated. The crude was purified by flash column chromatography (95% dichloromethane and 5% ethyl acetate) to give product **7** (214 mg, 79.8%). ^1^H NMR (400 MHz, DMSO-d6) δ 9.98 (s, 1H), 9.13 (br. s., 1H), 8.45 - 8.51 (m, 1H), 8.17 (s, 1H), 7.97 (d, *J* = 8.80 Hz, 1H), 7.79 (dd, *J* = 1.83, 8.80 Hz, 1H), 6.97 (d, *J* = 8.99 Hz, 2H), 3.89 (s, 3H), 3.83 (s, 3H), 1.46 (s, 9H). 13C NMR (101 MHz, DMSO-d6) δ 162.4, 159.8, 152.9, 138.8, 137.0, 135.8, 133.1, 131.0, 122.8, 122.5, 122.4, 121.5, 115.9, 78.3, 52.6, 36.2, 28.2. [M+H]^+^ LCMS C_21_H_24_N_3_O_5_S calculated. 430.50 found 430.1.

5-(4-((*tert*-butoxycarbonyl)amino)-1-methyl-*1H*-pyrrole-2-carboxamido)benzo[*b*]thiophene-2-carboxylic acid (**8**)

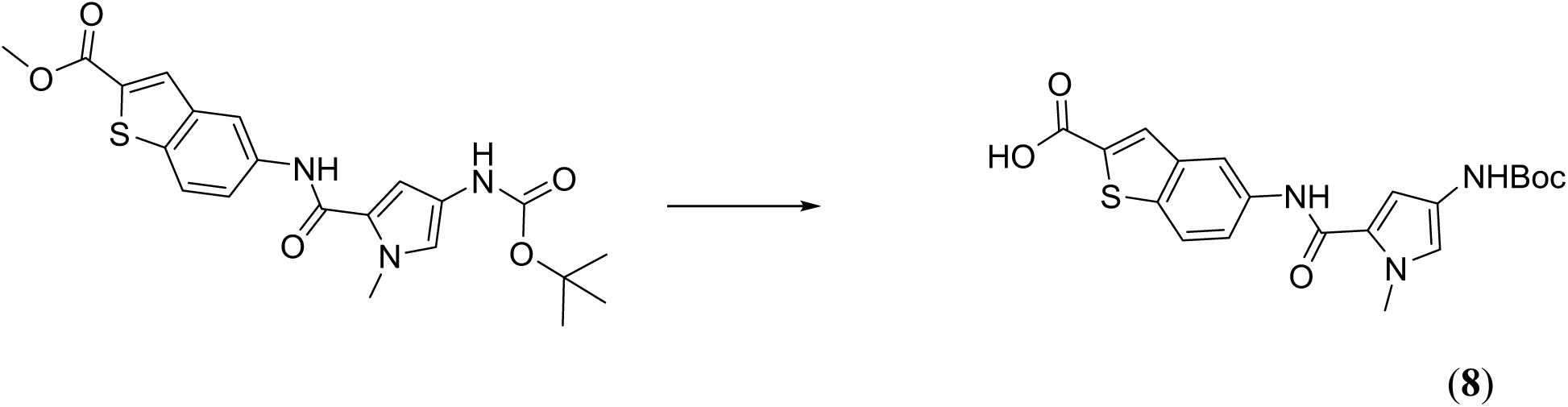

Compound **7** (78 mg, 0.182 mmol, 1 eq.) was dissolved in dioxane (2 mL) and sodium hydroxide (72.64 mg, 1.82 mmol; 10 eq.) dissolved in water (1 mL) was added dropwise. After 2 h, the dioxane was evaporated, and the residue dissolved in water (20 mL) and (**8**) acidified by the dropwise addition of 1 M HCl to pH 2.0. The crude was extracted using ethyl acetate (3x 30 mL), the combined organic fractions dried over magnesium sulfate and filtered, and the solvent evaporated to give compound **8** (42 mg, 56%). ^1^H NMR (400 MHz, DMSO-d6) δ 9.96 (s, 1H), 9.13 (s, 1H), 8.44 (d, *J* = 1.83 Hz, 1H), 8.05 (s, 1H), 7.94 (d, *J* = 8.80 Hz, 1H), 7.76 (dd, *J* = 2.02, 8.80 Hz, 1H), 6.97 (d, *J* = 6.24 Hz, 2H), 3.83 (s, 3H), 1.46 (s, 9H). 13C NMR (101 MHz, DMSO-d6) δ 163.6, 159.9, 152.9, 139.1, 136.9, 135.9, 130.2, 122.8, 122.6, 122.5, 121.1, 115.8, 104.8, 48.6, 40.2, 40.0, 39.8, 39.6, 36.2, 28.2. [M+H]^+^ LCMS C*20*H*22*N*3*O*5*S calculated. 416.47 found 416.1.

Ethyl 4-(5-(4-((*tert*-butoxycarbonyl)amino)-1-methyl-*1H*-pyrrole-2-carboxamido)benzo[*b*]thiophene-2-carboxamido)-1-methyl-*1H*-imidazole-2-carboxylate (**9**)

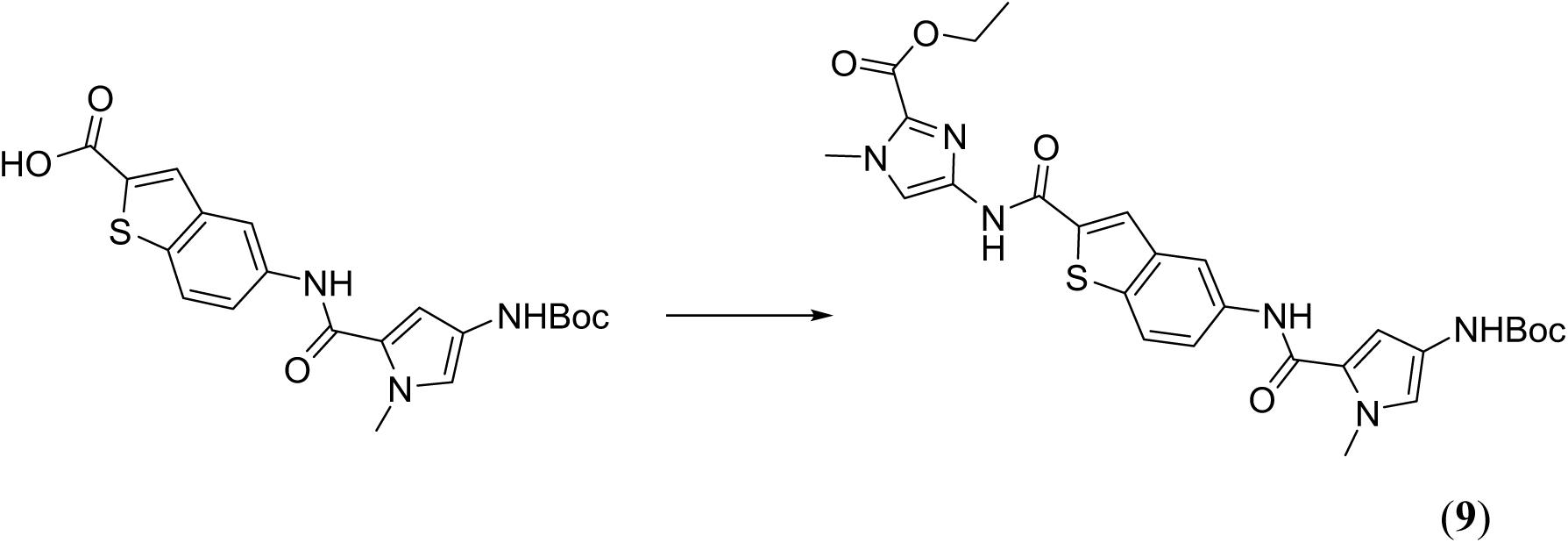

Compound **8** (40 mg, 0.096 mmol, 1.0 eq.) was dissolved in DMF (2.5 mL) and activated with EDC (2.0 eq, 36.91 mg, 0.193 mmol) and DMAP (2.5 eq, 29.41 mg, 0.241 mmol) for 30 min with stirring. Ethyl 4-amino-1-methyl-1*H*-imidazole-2-carboxylate (1.5 eq, 24.43 mg, 0.144 mmol) was added, and the mixture stirred overnight. The reaction was quenched with icecold water (50mL), extracted with ethyl acetate (3x 50 mL), dried over magnesium sulfate, filtered, and solvent evaporated. The crude product was dissolved in DCM (2.5 mL) and TFA (1 mL) was added. The mixture was stirred for 1 h, solvent evaporated, and purified by flash column chromatography (95% ethyl acetate and 5% methanol) to give product **9** (18 mg, 55%). ^1^H NMR (400 MHz, DMSO-d6) δ 11.52 (s, 1H), 9.69 (s, 1H), 8.37 (s, 2H), 7.92 (d, *J* = 8.80 Hz, 1H), 7.67 - 7.78 (m, 2H), 6.52 (d, *J* = 2.02 Hz, 1H), 6.36 (d, *J* = 1.83 Hz, 1H), 4.30 (q, *J* = 7.15 Hz, 2H), 3.96 (s, 3H), 3.76 (s, 3H), 1.33 (t, *J* = 6.42 Hz, 3H). 13C NMR (101 MHz, DMSO-d6) δ 159.2, 158.5, 139.7, 137.2, 137.1, 134.9, 131.8, 122.6, 116.1, 115.9, 115.8, 104.2, 41.6, 35.8, 35.6, 31.6, 29.4, 22.1, 14.1. [M+H]^+^ LCMS C_22_H_23_N_6_O_4_S calculated. 467.53 found 467.1.

Ethyl (*S*)-4-(5-(4-(4-((7-methoxy-5,11-dioxo-2,3,5,10,11,11a-hexahydro-1H-benzo[e]pyrrolo[1,2-a][1,4]diazepin-8-yl)oxy)butanamido)-1-methyl-1H-pyrrole-2-carboxamido)benzo[b]thiophene-2-carboxamido)-1-methyl-1H-imidazole-2-carboxylate (**10**)

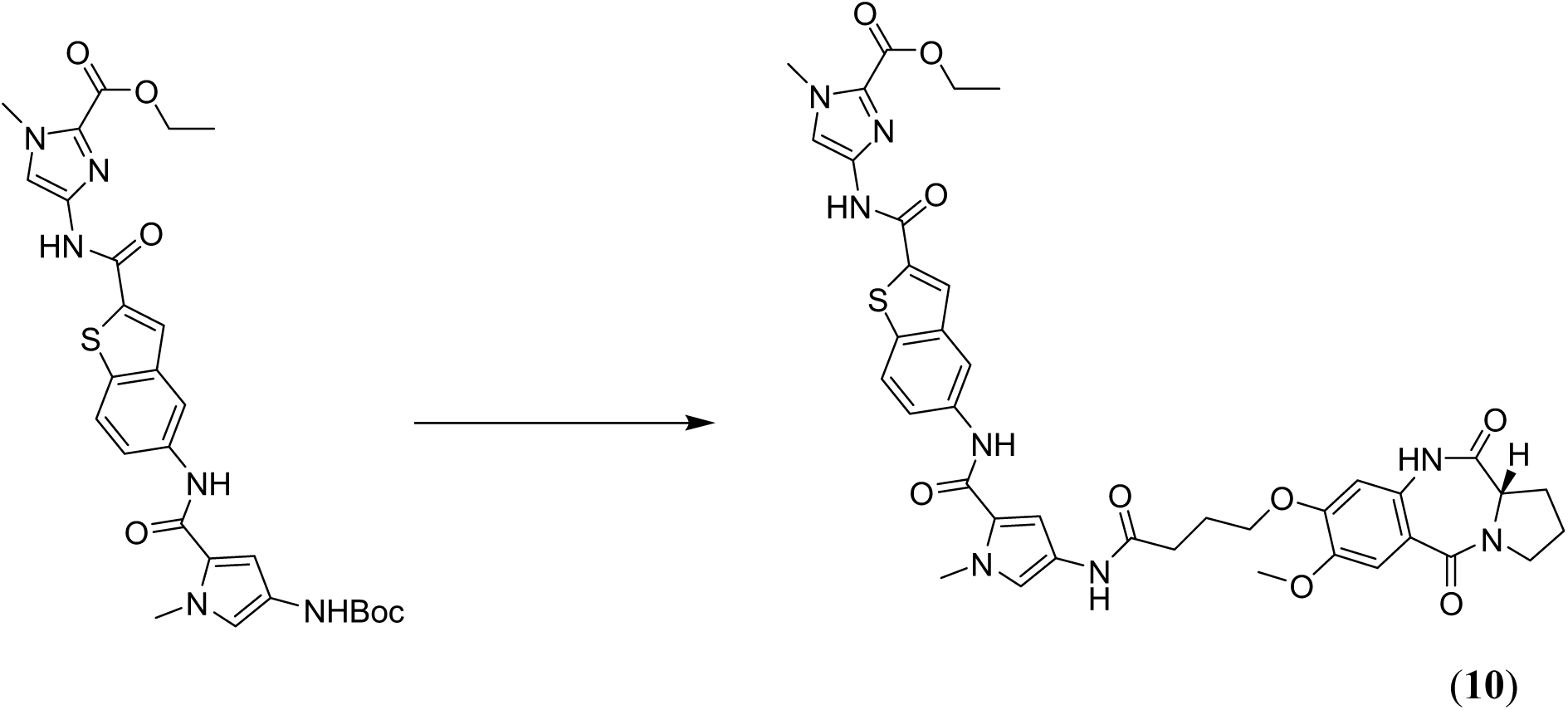

To a solution of compound 9 (50 mg, 0.11 mmol) in dioxane (3 mL), a 1 M solution of hydrochloric acid in dioxane was added. The mixture was stirred **at** room temperature for 3 hours and then concentrated in vacuo. The resulting solid was re-dissolved in DMF (2 mL) and added to a mixture of compound 6 (42 mg, 0.12 mmol), EDC (58 mg, 0.3 mmol), and DMAP (44 mg, 0.36 mmol) in DMF (1 mL). The resulting reaction mixture was stirred at room temperature for 16 hours. This was then diluted with brine (30 mL) and the product was extracted into ethyl acetate (3 × 10 mL). The combined organic fractions were dried over anhydrous Na_2_SO_4_, filtered, and concentrated in vacuo. The crude residue was purified by flash column chromatography eluting with dichloroethane and acetone (from 0% to 40%) to afford 40 mg of pure product as a light pink solid (66% yield).

_1_H NMR (400 MHz, DMSO) δ 11.54 (s, 1H), 10.21 (s, 1H), 9.98 (s, 1H), 9.94 (s, 1H), 8.40 (d, *J* = 1.8 Hz, 2H), 7.96 (d, *J* = 8.7 Hz, 1H), 7.77 (dd, *J* = 8.8, 2.1 Hz, 1H), 7.73 (s, 1H), 7.28 – 7.23 (m, 2H), 7.04 (d, *J* = 1.9 Hz, 1H), 6.70 (s, 1H), 4.31 (q, *J* = 7.1 Hz, 2H), 4.13 – 3.98 (m, 4H), 3.98 (s, 3H), 3.87 (s, 3H), 3.80 (s, 3H), 3.55 (td, *J* = 8.6, 4.5 Hz, 1H), 3.51 – 3.40 (m, 1H), 2.47 (t, *J* = 7.3 Hz, 3H), 2.08 (p, *J* = 6.9 Hz, 2H), 1.99; ^13^C NMR (101 MHz, DMSO) δ 170.82, 169.32, 164.83, 160.32, 159.57, 158.91, 151.41, 145.77, 140.25, 140.13, 137.68, 137.32, 135.64, 131.75, 131.25, 126.44, 123.14, 123.00, 122.61, 120.90, 119.51, 118.96, 116.22, 115.89, 112.41, 105.68, 105.43, 68.25, 61.17, 56.81, 56.14, 47.30, 36.73, 36.02, 32.28, 26.19, 25.16, 23.61, 14.57; [M+H]^+^ LCMS C_22_H_23_N_6_O_4_S calculated. 797.27 found 797.2. ^1^H NMR and ^13^C NMR spectra are shown in (**Suppl. Fig. 18b, c**).

### Expression and purification of AlbAS, AlbA, and AlbA^F143A^

Expression and purification of *K. oxytoca* His_6_-AlbAS (UniProt Q8KRS7, residues 1-221) has been described previously^2^. For AlbA expression, a pET-28a(+)-TEV plasmid encoding codon-optimized (for *E. coli* expression) *K. oxytoca* His_6_-AlbA (UniProt A0A9P0U153, residues 1-348) was obtained from Genscript (USA). The AlbA^F143A^ variant was generated by site-directed mutagenesis using the Q5 Site-Directed Mutagenesis Kit (NEB). Primers are reported in **Suppl. Table 6**. All constructs were verified by plasmid sequencing.

*E. coli* BL21 cells were transformed with the appropriate plasmid, and small overnight cultures in LB medium supplemented with kanamycin were grown at 37°C and used to inoculate larger-scale cultures in TB medium (1-4 L), which were grown at 37°C to an OD600 of 0.4-0.6. The temperature was then lowered to 18°C and protein synthesis was induced with 200 μM IPTG. After overnight expression, cultures were centrifuged at 6000 g for 30 min at 4°C, and the pellet was either processed immediately or stored at -80°C. The pellet was resuspended in Buffer A (50 mM Tris-HCl, 500 mM NaCl, 20 mM imidazole, 10% glycerol, 1 mM DTT, pH 7.5) supplemented with a tablet of protease inhibitor cocktail (Roche), and cells were lysed by sonication. Insoluble material was removed by centrifugation at 35,000 g for 30 min at 4°C, and the clarified lysate (filtered through a 0.45 μm membrane) was loaded onto a 1 mL HisTrap column (Cytiva) and eluted with a 0-100% gradient of Buffer B (50 mM Tris-HCl, 500 mM NaCl, 500 mM imidazole, 10% glycerol, 1 mM DTT, pH 7.5) on an ÄKTA Pure chromatography system. Selected fractions were buffer-exchanged into SEC buffer (50 mM Tris-HCl, 100 mM NaCl, 1 mM DTT, pH 7.5) using a HiPrep 26/60 Desalting column (Cytiva), and the His_6_-tag was cleaved overnight at 20°C with TEV protease added at a 1:100 protease:protein mass ratio. The sample was then loaded onto a 1 mL HisTrap column, the flow-through was collected, concentrated and loaded onto a Superdex 16/60 200 column. Selected fractions were pooled, concentrated, flash-frozen in liquid nitrogen and stored at -80°C.

### Expression and purification of RNA polymerase (RNAP)

*E. coli* RNAP was encoded on the PVS10 plasmid obtained from the Addgene repository (Addgene plasmid #104398)^4^. *E. coli* BL21 cells were transformed with the plasmid, and small overnight cultures in LB medium supplemented with ampicillin were grown at 37°C and used to inoculate larger-scale cultures in TB medium (2 L), which were grown at 37°C to an OD600 of 0.7-0.8. The temperature was then lowered to 30°C and protein synthesis was induced with 1 mM IPTG. After 5 h of expression, cultures were centrifuged at 6000 g for 30 min at 4°C, and the pellet was either processed immediately or stored at -80°C. The pellet was resuspended in Buffer A (50 mM Tris-HCl, 500 mM NaCl, 20 mM imidazole, 5% glycerol, 1 mM DTT, pH 6.9) supplemented with a tablet of protease inhibitor cocktail (Roche), and cells were lysed by sonication. Insoluble material was removed by centrifugation at 35,000 g for 30 min at 4°C, and the clarified lysate (filtered through a 0.45 μm membrane) was loaded onto a 1 mL HisTrap column (Cytiva) and eluted with a 0-100% gradient of Buffer B (50 mM Tris-HCl, 500 mM NaCl, 500 mM imidazole, 5% glycerol, 1 mM DTT, pH 6.9) on an ÄKTA Pure chromatography system. Selected fractions were buffer-exchanged into 95% Buffer AX (50 mM Tris-HCl, 5% glycerol, 0.5 mM EDTA, 1

mM DTT, pH 6.9) and 5% Buffer BX (50 mM Tris-HCl, 1.5 M NaCl, 5% glycerol, 0.5 mM EDTA, 1 mM DTT, pH 6.9) (AX95-BX5-buffer) using a HiPrep 26/60 Desalting column (Cytiva). The sample was then loaded onto a 5 mL Heparin Trap column (Cytiva) pre-equilibrated in AX95-BX5-buffer, washed and eluted with a gradient from 5% to 100% Buffer BX over 10 CV. Selected fractions were pooled, concentrated, and buffer-exchanged into AX95-BX5-buffer. The sample was then loaded onto a 6 mL RESOURCE Q column (Cytiva) pre-equilibrated in AX95-BX5-buffer, washed and eluted with a gradient from 5% to 100% Buffer BX over 20 CV. Selected fractions were pooled and buffer-exchanged into 86.7% Buffer AX and 13.3% Buffer BX, then supplemented with an equal volume of 100% glycerol to give a final storage buffer of 25 mM Tris-HCl, 100 mM NaCl, 0.25 mM EDTA, 0.5 mM DTT, 52.5% glycerol, pH 6.9. The protein was then flash-frozen in liquid nitrogen for long-term storage at -80°C.

### Expression and purification of σ^70^ subunit

*E. coli* σ^70^ factor was encoded on the PIA586 plasmid, obtained from the Addgene repository (Addgene plasmid #104399)^4^. *E. coli* BL21 cells were transformed with the plasmid, and small overnight cultures in LB medium supplemented with kanamycin were grown at 37°C and used to inoculate larger-scale cultures in TB medium (2 L), which were grown at 37°C to an OD600 of 0.7-0.8. The temperature was then lowered to 30°C and protein synthesis was induced with 1 mM IPTG. After 5 h of expression, cultures were centrifuged at 6000 g for 30 min at 4°C, and the pellet was either processed immediately or stored at -80°C. The pellet was resuspended in Buffer A (50 mM Tris-HCl, 500 mM NaCl, 20 mM imidazole, 5% glycerol, 1 mM DTT, pH 6.9) supplemented with a tablet of protease inhibitor cocktail (Roche), and cells were lysed by sonication. Insoluble material was removed by centrifugation at 35,000 g for 30 min at 4°C, and the clarified lysate (filtered through a 0.45 μm membrane) was loaded onto a 1 mL HisTrap column (Cytiva) and eluted with a 0-100% gradient of Buffer B (50 mM Tris-HCl, 500 mM NaCl, 500 mM imidazole, 5% glycerol, 1 mM DTT, pH 6.9) on an ÄKTA Pure chromatography system. Selected fractions were buffer-exchanged into 95% Buffer AX (50 mM Tris-HCl, 5% glycerol, 0.5 mM EDTA, 1 mM DTT, pH 6.9) and 5% Buffer BX (50 mM Tris-HCl, 1.5 M NaCl, 5% glycerol, 0.5 mM EDTA, 1 mM DTT, pH 6.9) (AX95-BX5-buffer) using a HiPrep 26/60 Desalting column (Cytiva). The sample was then loaded onto a 6 mL RESOURCE Q column (Cytiva) pre-equilibrated in AX95-BX5-buffer, washed and eluted with a gradient from 5% to 100% Buffer BX over 20 CV. Selected fractions were pooled and further polished by SEC using a Superdex 16/60 200 column (Cytiva) in 66.6% Buffer AX and 33.3% Buffer BX. The final sample was concentrated with a 10 kDa cutoff concentrator, supplemented with 100% glycerol to a final concentration of 5% glycerol, and flash-frozen in liquid nitrogen for long-term storage at -80°C.

### Reconstitution of the RNAP(σ^70^):DNA:AlbA complex

The RNAP(σ^70^):DNA:AlbA complex was reconstituted by incubating 375 μL of RNAP (0.67 mg/ml), 23 μL of σ^70^ (3.3 mg/ml), 15 μL of *palbA* (50 μM), 10 μL AlbA (22 mg/ml) for 1 h at 4°C. The approximate ratio between the components is 1:2:1.5:5. The sample was then purified on a Superose 6 10/300 Increase (Cytiva) in S6 buffer (10 mM HEPES, 100 mM KCl, 5 mM MgCl, 1 mM DTT, pH 7.5) and selected fractions were concentrated to about 5 mg/ml.

### SEC-MALS

150 μL of AlbA (2.4 mg/mL) were loaded on an Agilent WTC030S5 column equilibrated with 50 mM Tris-HCl pH 7.5, 100 mM NaCl using an Agilent 1260 Infinity II LC system coupled to DAWN MALS detector and Optilab dRI detector (Wyatt). Data were analyzed using the ASTRA software.

### Minimum Inhibitory Concentration (MIC) assay

MICs for each antibiotic were assessed by the broth microdilution method, following a modified protocol from Surani *et al*^2^. AlbAS expression in *E. coli* was induced with 200 μM IPTG at OD_600_ = 0.1 and protein synthesis was induced at 20°C for 1 hour. After that, the culture was diluted to OD_600_ = 0.08-0.13 (approximated McFarland standard of 0.5) and used to prepare the microdilution plate according to the protocol, except for the presence of 200 μM IPTG in each well and the growth at 28°C to minimize protein misfolding. Experiments were performed in triplicate.

### Equilibrium dissociation constant (*K*_D_) determination by fluorescence quenching

Ligand binding was monitored by intrinsic protein fluorescence quenching. For each ligand, a fixed concentration of protein was titrated with increasing total ligand concentration spanning approximately 0.1 nM to 10–50 µM, and emission spectra were recorded at each titration point (typically three replicate spectra per point). Two protein constructs were analyzed under an identical protocol: monomeric AlbAS and dimeric AlbA. A protein-only control and a ligand-only (protein-free) control were recorded to confirm the origin of the signal and to assess intrinsic ligand fluorescence at high concentration.

For each titration series, the emission peak wavelength was determined from the mean spectrum of the lowest ligand-concentration point, where quenching is minimal and the protein signal is maximal; all subsequent analysis used the fluorescence intensity at that fixed peak wavelength. At each titration point, the mean and standard deviation of the replicate intensities were computed and used as the observation and its weight, respectively.

Because several dissociation constants approach or fall below the total binding-site concentration, the standard hyperbolic model (which assumes free ligand equals total ligand) is invalid. Binding was therefore fit with the Morrison quadratic (tight-binding) equation, which explicitly accounts for ligand depletion. The concentration of bound complex [PL] at each total ligand concentration [*L*]*_t_* is

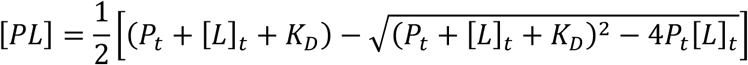

and the observed fluorescence F is modeled as

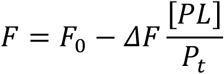

where *F*_0_ is the fluorescence of the free protein, *ΔF* the total quench amplitude at saturation, *K*_D_ the dissociation constant, and *P_t_* the total binding-site concentration.

*P_t_* was set from the concentration of protein binding sites. Crystallographic analysis shows two ligand molecules bound per AlbAS molecule; however, intrinsic tryptophan quenching reports total site occupancy through a single observable and does not resolve the individual sites. The titration data were well described by a single binding transition, and the fitted parameter is therefore reported as an apparent (macroscopic) dissociation constant (*K*_D,app_) for ligand binding to AlbAS rather than a site-resolved constant. The relationship between the two sites whether equivalent, independent, or cooperative cannot be determined from these data, and no site-resolved (*K*_D1_, *K*_D2_) or cooperative model was fit, because such parameters are not identifiable from a single occupancy signal measured over the accessible concentration range. For AlbAS, *P_t_* was set to 50 nM, i.e. the concentration of protein molecules, so that the reported *K*_D,app_ is expressed on a per-molecule basis.

For the highest-affinity ligands, *K*_D,app_ is comparable to or below *P_t_* (tight-binding / stoichiometric regime). In this regime the fitted value is influenced by the titration stoichiometry and depends on the assumed binding-site concentration: refitting against a two-site basis (100 nM sites) changed the fitted *K*_D,app_ by approximately 2- to 5-fold for these ligands. Such values are therefore interpreted as upper bounds rather than precise constants, whereas for the lower-affinity ligands (*K*_D,app_ above the site concentration) the fitted value is well determined and essentially independent of this choice. The rank order of ligand affinities across the series is preserved under either assumption and is the robust, model-independent output of the analysis. The second, lower-occupancy binding event could not be characterized. The ligand concentrations required to populate it coincide with strong intrinsic ligand fluorescence and inner-filter attenuation (see below), which corrupt the fluorescence signal in precisely that concentration range and preclude determination of a second dissociation constant. The inability to observe this event is an optical limitation of the assay and does not, in itself, constrain the affinity of the second site. For the dimeric AlbA construct, the two protomers each contribute one binding site on separate polypeptide chains. These were treated as independent and equivalent, an assumption appropriate to two chemically identical, spatially separate subunits which collapse to a single apparent *K*_D_ fit against *P_t_* = 50 nM total sites (dimer measured at 25 nM). No two-site or cooperative model was required for this construct.

*F*_0_, *ΔF*, and *K*_D_ were fit by non-linear least squares (Levenberg–Marquardt with parameter bounds; scipy.optimize.curve_fit), weighting each point by its replicate standard deviation with absolute error scaling (absolute_sigma = True), so that reported parameter errors are absolute rather than rescaled to the fit residuals. The asymptotic standard error (SE) on *K*_D_ was taken as the square root of the corresponding diagonal element of the covariance matrix, and the 95% confidence interval as

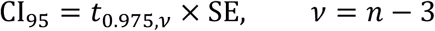

where *ν* = (number of points fit − 3) degrees of freedom. Goodness of fit is reported as the coefficient of determination *R²*.

High ligand concentrations are subject to intrinsic ligand fluorescence and inner-filter effects that corrupt the measured signal in one of two ways: fluorescence rising back above the quenched minimum (ligand autofluorescence), or continuing to fall below the true binding plateau (inner-filter attenuation). A single objective exclusion rule was applied identically to all datasets of both constructs. Working inward from the highest concentration, a titration point was excluded if it either (i) exceeded the interior minimum of the retained points by more than three standard deviations, or (ii) fell below the fitted saturation plateau (F_0_ − ΔF, estimated from the lower-concentration points) by more than 8% of the quench amplitude. Exclusion stopped at the first outermost point satisfying neither criterion, and a minimum of six points was always retained.

### X-ray crystallography

Purified AlbAS was concentrated to ∼30 mg/mL and a 100% DMSO stock of each ligand was added to the concentrated protein solution to reach a nominal stoichiometric excess of 3-5:1 and a final DMSO percentage of 1-2%. The suspension was incubated overnight at 4°C. Before setting up crystallization drops, the suspension was centrifuged for at least 30 minutes at 13,000g to separate the insoluble ligand. Crystallization plates were set up with an Oryx8 crystallization robot (Douglas Instruments). Crystals of AlbAS in complex with KMR-04-154, KMR-04-163, AlbAS:KMR-04-177, AlbAS:KMR-28-27, were obtained by vapor diffusion in sitting drops in 1.0-2.0 M ammonium sulfate. Crystals of AlbAS:KMR-04-161 and AlbAS:KMR-04-165 were obtained also by vapor diffusion in the presence of 0.15 M ammonium sulfate, 0.1 M MES pH 6.5, 25% PEG 4000. For data collection, crystals were cryoprotected by the addition of either 2 M sodium malonate or 25% (v/v) ethylene glycol and cryo-cooled in liquid nitrogen. Data were collected remotely at the European Synchrotron Radiation Facility (ESRF, Grenoble) and at Diamond Light Source (DLS, Didcot). Data were processed using AUTOPROC^5^ and DIALS^6^ via automated pipelines available at the beamlines. The structures were solved by the molecular replacement (MR) technique using the software package PHASER^7^ starting from protein coordinates of the AlbAS-KMR-14-14 complex (PDB code 8RKY). Model building was performed using *COOT*^8^ and crystallographic refinement was carried out with either REFMAC5^9^ or PHENIX.REFINE^10^. Stereochemical restraints for the organic ligands were generated using GRADE2 (Global Phasing Ltd.). A summary of data collection and refinement statistics are shown in **Suppl. Table 2**.

### AI-driven model generation of the AlbA dimer

Models for the AlbA dimer were initially generated using the web-based versions of AlphaFold2^11^ (v1.5.2, Colab implementation) and AlphaFold3^12^ by providing the *K. oxytoca* sequence for this protein (UniProt A0A9P0U153, residues 1-348). Default parameters were used. To produce models restrained to the experimental cryo-EM structure of the AlbA LBD dimer we initially utilized Chai-1^13^ (v0.6.1) via its web-interface. The following intermolecular distance restraints were provided (V140-T215, T215-V140, 4 Å; N151-Q139, A192-Q189, Q139-N151, 6 Å; A192-A192, 7 Å; G193-G193, 13 Å; E312-E312 20 Å; R340-R340 42 Å; R256-R256 50 Å). Next, we generated a pool of 2,000 dimeric models sampling alternative relative domain orientations using Boltz-2^14^ (v2.2.1) on the Leonardo HPC system (CINECA). Of these, 1,000 models were generated without a template, 500 models were restrained to the experimental apo cryo-EM structure (LBDs only), and 500 were restrained to the Chai-1 model (DBD-out/LBD-close) that had itself been restrained to the cryo-EM structure. Calculations were performed using Python v3.11.7, PyTorch v2.6.0+cu124, and the CUDA 12.4 runtime. The multiple sequence alignment (MSA) was generated using ColabFold v1.6.2 and its public MMseqs2 server via ‘colabfold.colabfold.run_mmseqs2’.

The search was performed with default settings, environmental sequences and ColabFold’s MSA filter enabled. The same locally generated ColabFold MSA was used for all predictions. All three pools were generated using the same inference settings and sampling strategy.

Predictions were performed in independent sampling campaigns consisting of arrays of 25 tasks, with each one generating 20 diffusion samples. Models were generated using 10 recycling steps and 200 diffusion sampling steps. Independent campaigns used distinct random seeds while retaining identical input sequences, MSA, and inference settings; the only difference between pools production was the structural template information supplied during prediction. The directly restrained set reproduces the apo arrangement to a median 1.95 Å over the LBDs, the indirectly restrained set to 4.68 Å, and the unrestrained set comes within 9.89 Å of it. Models were then screened for CC integrity by taking, for each CC residue of one chain, the distance to the nearest coiled-coil residue of the partner. Any model in which that distance exceeded 14 Å at more than a fifth of positions was rejected, which removed 15 dimers whose two helices had dissociated and left a final pool of 1,985 models.

### Small angle X-ray scattering (SAXS)

For AlbA, we collected size-exclusion chromatography-coupled SAXS (SEC-SAXS) data at the P12 bioSAXS beamline (DESY, Hamburg) using a Pilatus 6M (Dectris) detector at a wavelength of 0.124 nm. A volume of 100 μl at the concentration of 10 mg/ml in 50 mM Tris-HCl pH 7.5, 150 mM NaCl, 1 mM DTT was injected onto a Superdex 200 Increase 10/300 column (Cytiva) and eluted at 0.5 mL/min. SAXS data for AlbA^F143A^ (1.25 mg/ml in the same buffer) were collected in batch mode at the BM29 BioSAXS beamline (ESRF, Grenoble) using a Pilatus3 2M (Dectris) detector at a wavelength of 0.099 nm. Both experiments were performed at 20°C. All data processing and analyses were carried out with software tools from the ATSAS suite^15^. AlbA data in the *q* range (0.0020-0.4198 Å^−1^) were processed at the beamline with the CHROMIXS package while AlbA^F143A^ scattering data in the *q* range (0.0050-0.5575 Å^−1^) were corrected for the buffer contribution using PRIMUS with the high-angle region as reference (scale factor 0.9950). The radius of gyration and forward intensity were obtained with AUTORG, and the pair distance distribution with GNOM. Molecular weight was estimated from the volume of correlation with DATVC, using the AUTORG *R*_g_ and *I*(0) as required inputs, and shape classification was performed with DATCLASS. Uncertainties are quoted only where the program reports one. Where a *R*_g_ is compared between the two proteins the same estimator is used for both.

Theoretical scattering for single models was calculated with CRYSOL while theoretical scattering for the ensemble computed with FFMAKER, which applies a common hydration-shell contrast to every model. The solvent electron density was fixed at 0.334 e Å^−3^ throughout. The shell contrast was set to 0.060 e Å^−3^ for AlbA and 0.030 e Å^−3^ for AlbA^F143A^, each chosen as the minimum of a contrast scan over the fit quality of the resulting ensemble. Mixtures were selected from the pool in two independent ways: by non-negative least squares using NNLSJOE and by the genetic algorithm GAJOE. Composition of the ensemble was assessed over all independent fits of each protein rather than from a single run, comprising 26 fits for AlbA and 10 for AlbA^F143A^. These combine Monte Carlo replicates drawn within the measurement uncertainties, repeated GAJOE runs from independent seeds, and both selection algorithms. They also include a jackknife in which each selected conformer was withheld from the pool in turn and the fit repeated. Quoted means and standard deviations are taken over that set.

For conformational classification, each model was placed on two axes computed from coordinates. The DBD axis is the fraction of DNA-binding domain surface buried in the dimer. It is one minus the ratio of the accessible surface area of residues (5-70) of both subunits in the assembled dimer to the area of the same residues in isolation. Buried fractions above 0.341 define DBD-in. That boundary sits in an empty region of the pool distribution, and it agrees with a contact-based criterion for all but four of the models. Boundaries were chosen by visual inspection and well describe the bimodal distribution along both axes.

### Cryo-EM

#### Grid preparation

For AlbA, freshly purified AlbA at 2 mg/mL was used to prepare cryo-EM grids. An EM GP2 Plunge Freezer (Leica) was used to prepare UltrAuFoil grids, which were glow discharged for 30 seconds at 15 mA using an EM ACE200 (Leica). 3 μL of the sample were applied to each grid at 90% humidity level, waiting for 1 minute before front blotting for 2 s. Immediately after, the grid was plunged in liquid ethane at -180°C to vitrify the sample. The grid was then carefully transferred to liquid nitrogen at -196°C and clipped using autogrid rings and C-clips.

For the AlbA:KMR-14-14 and AlbA:KMR-28-27 complexes, a 2 mg/mL protein solution was incubated with a 10:1 ligand ratio (0.6% DMSO in the final solution) for 3 h at 4°C. The sample was then centrifuged for 30 minutes at 13,200 rpm before preparing grids as for AlbA.

For RNAP(σ^70^):DNA:AlbA, to 20 μL of the complex at 5 mg/ml we added 2.5 μL of AlbA (22 mg/ml), 0.8 μL of KMR-28-27 (125 μM in 25% DMSO) and 0.5 μL CHAPSO (10% w/v, 160 mM) was added. A 3 μl volume of the sample was blotted on a Quantifoil C-Au using a Vitrobot (ThermoFisher) at 4°C, 100% humidity and plunged in liquid ethane.

#### Data collection

AlbA and AlbA:KMR-14-14 datasets were collected with a Krios G3i (ThermoFisher) operating at 300 kV, with a Gatan K3 camera and a BioQuantum imaging filter. Movies were collected with EPU (Thermo Fisher) software with a super-resolution mode sampling of 0.54 Å/px (165 kX magnification). A total of 16987 micrographs were collected for AlbA and 24100 micrographs for AlbA:KMR-14-14, using a defocus range of -0.6 to -1.8 μm and a total dose of 70 e−/Å² distributed over 70 frames. AlbA:KMR-28-27 dataset was collected at University of Padova on a Glacios (ThermoFisher) microscope operating at 200 kV with a Falcon 4i camera. Movies were collected with EPU (Thermo Fisher) software at 1.2 Å/px (120 kX magnification). A total of 5239 micrographs were collected using a defocus range of -0.8 to -2 μm and a total dose of 60 e−/Å² distributed over 60 frames.

Screening of RNAP(σ70):DNA:AlbA(KMR-28-27) grids was performed at the DSB-CEM facility at the University of Padova on a 200 kV Glacios (ThermoFisher) microscope equipped with a Falcon 4i camera (ThermoFisher) and collected at eBIC on a Titan Krios (ThermoFisher) microscope operating at 300 kV with a Gatan K3 camera and a BioQuantum imaging filter. Movies were collected with EPU (ThermoFisher) software at 0.825 Å/px (105 kX magnification). A total of 24521 micrographs were collected using a defocus range of -0.8 to -2 μm and a total dose of 40 e−/Å² distributed over 40 frames.

#### Data processing

Near-identical processing pipelines in CryoSPARC^16^ were used for AlbA, AlbA:KMR-14-14 and AlbA:KMR-28.27 datasets. These are schematized in **Suppl. Fig. 9, 12, 13**, respectively. Movies were motion corrected using the Patch Motion Correction implementation and the CTF was estimated using the Patch CTF implementation. Manual picking and subsequent 2D classification were used to generate templates for template picking. After a few rounds of 2D classification, *ab initio* reconstruction followed by heterogeneous refinement of each class was performed. The most promising classes were subjected to several rounds of HR-HAIR^17^ and local refinement. Reference-based motion correction was then performed before a final round local refinement.

For the RNAP(σ70):DNA:AlbA(KMR-28-27) complex processing was also carried out using CryoSPARC^16^ (**Suppl. Fig. 19**). Briefly, movies were subjected to patch motion correction and patch CTF estimation. After manual picking of a subset of particles, template-based picking and particle extraction, RNAP particles were selected after several rounds of 2D classification. *Ab initio* reconstruction into 5 classes and heterogeneous refinement were used to reconstruct a volume, which was then subjected to 3D classification into 10 classes. The classes containing visible AlbA density were further processed through non-uniform refinement, heterogeneous refinement, masked 3D classification and reference-based motion correction, yielding the consensus reconstruction at 2.43 Å (gold-standard FSC, 0.143 criterion). Particle subtraction of the RNAP core and masked local refinement focused on the AlbA-containing region then produced the focused reconstruction of AlbA at 3.03 Å resolution.

#### Model building

For apo AlbA and AlbA:PBD complexes, the starting model was taken from PDB 8RKY after removal of all heteroatoms. For the RNAP(σ70):DNA:AlbA(KMR-28-27) complex we used the RNAP core of PDB 6XL5 while for the DBD/CC regions of AlbA we employed AlphaFold2 predictions. The DBD region was taken from 8RKY. Restraints for KMR-14-14 and KMR-28-27 were generated using GRADE2. Regions not included in the starting compostite models were built manually into density using Coot. The models were refined using a combination of Servalcat, Phenix.refine and ISOLDE^83^.

### Structural analysis of promoter spacer DNA

UCSF Chimera was used to measure the spacer lengths and kink angles. The spacer length is defined as the centroid-to-centroid distance between the first and last base pairs of a promoter spacer DNA. The kink angle is defined as the angle between the centroids of the first six base pairs, central seven base pairs and last six base pairs, respectively, of a 19-bp spacer.

Promoter spacer DNA conformation was analyzed by W3DNA 2.0 web server^18^. The total helical twist angles of the promoter spacers were calculated by extrapolating the average base-pair step helical twist (h-twist) parameter over 19 baes-pair steps. The under-twist value of each spacer DNA was calculated by subtracting its total helical twist angle from the total helical twist angle of an ideal B-form 19-bp DNA.

### SELEX

An 82 bp oligonucleotide library, containing a 42 bp random region flanked by two 20 bp primer regions containing BamHI and EcoRI restriction sites (ssDNA-SELEX) was obtained from Eurogentec (Belgium) and complementary strand synthesis was carried out using the Klenow fragment (ThermoFisher). The dsDNA-SELEX library produced was purified using the Monarch PCR and DNA cleaning kit (Neb) following the manufacturer’s instructions.

The dsDNA-SELEX library was then amplified by PCR using forward primer (SELEX-fwd) and reverse primer (SELEX-rev) using Vent DNA polymerase (Neb). The PCR program was as follows: 95°C 3 min, 25 cycles of 95°C 30 sec and 60°C 15 sec; 72°C 5 min. The annealing temperature of 60°C was selected after screening temperatures between 50°C and 70°C as it maximized the correct amplification product. For subsequent rounds of selection, an annealing temperature gradient between 52-60°C was tested and the cleanest PCR products were purified using the Monarch PCR and DNA Cleaning Kit (Neb) and used for the next selection round. When needed, agarose gel purification (2% agarose) using the Monarch DNA Gel Extraction Kit (Neb) was carried out to purify the correct band after PCR amplification due to buildup of side products.

The dsDNA-SELEX library (500 ng) was incubated with AlbA at protein:DNA ratios of 5:1, 20:1 and 40:1 for a varying amount of time and with increasingly stringent conditions during subsequent rounds of selection, up to 500 mM NaCl and 500 ng/μL sssDNA. 2 μL of 5×DNA-binding buffer (5×DBB, 50 mM Tris-HCl pH 7.5, 250 mM NaCl, 2.5 mM EDTA, 15 mM MgCl2, 25% glycerol, 1 mM DTT) were used to prepare 10 μL of each protein:DNA sample, which was then loaded on a 6% polyacrylamide gel (made in 0.5×TBE, pre-ran for 1 hour at 4°C) and run for 1 hour in 0.5×TBE. The gel was then stained with ethidium bromide in 0.5×TBE for 30 minutes. The protein:DNA gel bands were then excized and soaked in 100 μL diffusion buffer (500 mM NH4Ac, 10 mM Mg(Ac)2, 1 mM EDTA, 0.1% SDS, pH 8.0) overnight at 37°C. The diffusion buffer was then purified using the Monarch PCR and DNA Cleaning Kit and the obtained selected *palbA*-SELEX DNA was amplified by PCR and checked using a 2% agarose gel. Selected and purified *palbA*-SELEX DNA after each cycle was used for the subsequent selection cycle.

The selected *palbA*-SELEX library was then cloned in the pUC19 plasmid for sequencing. pUC19 vector was double digested using EcoRI-HF (Neb) and BamHI-HF (Neb). 500 ng of digested pUC19 was treated with FastAP Thermosensitive Alkaline Phosphatase (ThermoFischer). Digested and dephosphorylated pUC19 was purified by agarose gel electrophoresis and Monarch DNA Gel Extraction Kit following the manufacturer’s instructions. *palbA*-SELEX after Round 7 of selection was also double digested with EcoRI-HF (Neb) and BamHI-HF (Neb). After that, digested DNA was purified using agarose gel electrophoresis and Monarch DNA Gel Extraction Kit following the manufacturer’s instructions. Digested and dephosphorylated pUC19 vector and digested *palbA*-SELEX (10:1 insert:vector ratio) were ligated using T4 DNA ligase (Promega). The ligation mixture was used to transform *E. coli* TOP10 competent cells, and blue-white screening was used to select 95 white colonies, which were picked and grown overnight before Direct Colony Sequencing (Eurofins Genomics). The M13rev-49 primer (GAGCGGATAACAATTTCACACAGG) was used for sequencing. Motif analysis was performed with the MEME Suite.

### Fluorescence Polarization (FP)

3’-TAMRA-derivatized DNA oligonucleotides (Merck) for the 5’-3’ strands of *palbA* and *palbA*-SELEX-A8 were annealed with their non-fluorescent complementary oligonucleotides using a thermocycler (95°C for 2 minutes, then slowly cooling to 25 °C for 1 hour). For the FP experiment, the fluorescent duplexes were incubated for 1 hour at 4 °C with increasing concentrations of AlbA and analyzed using a TECAN Spark (TECAN). 20 μL of sample were dispensed in a black 384-well plate.

Polarization values were baseline-corrected against the DNA-only signal, and each titration was fitted to a one-site binding model incorporating a linear nonspecific term:

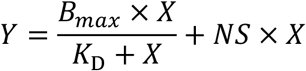

where *X* is the total protein concentration, *B_max_* the amplitude of specific binding, *K*_D_ the dissociation constant of the saturable component, and *NS* the slope of the nonspecific contribution. The background term was constrained to zero for all fits, since the data were baseline-corrected. Replicate wells were treated as individual observations and fits were unweighted. Reported *K*_D_ values therefore describe the saturable component only. Individual wells whose robust standardized residual exceeded 3 against the corresponding fit were excluded (10 of 249 measurements). The residual scale was estimated as 1.4826 × the median absolute deviation from a soft-L_1_ robust fit to the complete dataset, so that the exclusion threshold was not itself influenced by the outlying points. Fits obtained without any exclusions are reported in Supplementary Table 5. Three *palbA* titrations (AlbA, AlbA+AF-CD-007, AlbA+KMR-28-27) were acquired under matched conditions and were adequately described by a single nonspecific slope (F-test for a shared versus three independent *NS* values, P = 0.97). They were therefore fitted globally with *NS* as one shared parameter (*NS* = 0.73 ± 0.04 mP µM^−1^), which reduces the covariance between *B_max_* and *NS* and narrows the confidence interval on each *K*_D_. AlbA^F143A^ and the *pSELEX* duplex were fitted individually with their own nonspecific slopes, because in each case *NS* differed significantly from the wild-type *palbA* value. Differences in *K*_D_ and *B_max_* between conditions were assessed by F-tests comparing nested models in which the parameter of interest was either shared between two conditions or fitted independently. Confidence intervals on *K_D_* were obtained by profile likelihood, since the likelihood is asymmetric on a logarithmic concentration scale; the symmetric standard errors given in (**Fig. 5d, e**) are provided for comparison and are narrower than the profile intervals at the upper bound. Non-linear regression was performed in Python 3.11 using the *least_squares* routine of SciPy 1.17 (Trust Region Reflective algorithm), with all parameters constrained to be non-negative.

### *In vitro* transcription activity assay

To prepare the 178 bp and 176 bp DNA sequences containing the *palbA* promoter, PCR amplification from the *palbA* and *palbA-17* plasmids was used. 5.0 μL 10×ThermoPol Reaction buffer (NEB), 1 μL 10 mM dNTPs (Promega), 0.25 μL 100 μM forward primer (AlbA-SELEX-fw), 0.25 μL 100 μM reverse primer (AlbA-SELEX-rev, Sigma), 0.25 μL Vent DNA polymerase (2 U/μL, NEB), 1 μL of *palbA*-TA (0.3 ng/μL), and 42 μL sterile ddH2O. The PCR program was as follows: 95°C 3 min, 30 cycles of 95°C 1 min and 57°C 45 sec; 72°C 5 min. The PCR product was checked by agarose gel electrophoresis (2% agarose) and then purified using the Monarch PCR and DNA Cleaning Kit (NEB). The *palbA*-17 plasmid was obtained from the *palbA* plasmid using the Q5 Mutagenesis Kit (NEB) using *palbA*-TA-fwd and *palbA*-TA-rev plasmids. The PCR program was as follows: 98°C 30 sec, 25 cycles of 98°C 10 sec, 69°C 30 sec and 72°C 70 sec; 72°C 2 min. For each sample, to a master mix composed of 12 μL of deionized water, 3 μL of 10X TA buffer (1X: 40 mM Tris-HCl, 150 mM NaCl, 2 mM MgCl_2_, 1 mM DTT, pH 7.5), 6 μL of RNAP holoenzyme (1 μM) and 1.5 μL DNA (350 nM), 1.5 μL of AlbA (30 μM) were added. In the case of ligands, these were incubated with AlbA prior to addition to the master mix. The transcription reaction was then initiated with 6 μL of rNTPs (500 μM, ThermoFisher) and incubated at 37°C for 2 hours. The reaction was then stopped with the addition of 2X RNA Loading Buffer (ThermoFischer), incubated at 70°C for 10 minutes before loading into a 10% urea-PAGE gel. The gel was run at 180 V for 1 hour before staining with ethidium bromide and visualized with a Uvitec NineAlliance documentation station.

